# A Novel Subtype of Myeloproliferative Neoplasms Driven by a MYC-Alarmin Axis

**DOI:** 10.1101/2023.09.25.559138

**Authors:** Nicole D. Vincelette, Xiaoqing Yu, Andrew T. Kuykendall, Jungwon Moon, Siyuan Su, Chia-Ho Cheng, Rinzine Sammut, Tiffany N. Razabdouski, Hai Vu Nguyen, Erika A. Eksioglu, Onyee Chan, Najla Al Ali, Parth C. Patel, Dae Hyun Lee, Shima Nakanishi, Renan B. Ferreira, Qianxing Mo, Suzanne Cory, Harshani R. Lawrence, Ling Zhang, Daniel J. Murphy, Rami S. Komrokji, Daesung Lee, Scott H. Kaufmann, John L. Cleveland, Seongseok Yun

## Abstract

Despite advances in understanding the genetic abnormalities in myeloproliferative neoplasms (MPNs) and the development of JAK2 inhibitors, there is an urgent need to devise new treatment strategies, particularly for triple negative myelofibrosis (MF) patients whose MPNs lack mutations in the JAK2 kinase pathway and have very poor clinical outcomes. Here we report that *MYC* copy number gain and increased MYC expression frequently occur in triple negative MF, and that MYC-directed activation of S100A9, an alarmin protein that plays pivotal roles in inflammation and innate immunity, is necessary and sufficient to drive development and progression of MF. Notably, the MYC-S100A9 circuit provokes a complex network of inflammatory signaling that involves various hematopoietic cell types in the bone marrow microenvironment. Accordingly, genetic ablation of *S100A9* or treatment with small molecules targeting the MYC-S100A9 pathway effectively ameliorates MF phenotypes, highlighting the MYC-alarmin axis as a novel therapeutic vulnerability for this subgroup of MPNs.

**SIGNIFICANCE:** This study establishes that MYC expression is increased in triple negative MPNs via trisomy 8, that a MYC-S100A9 circuit manifest in these cases is sufficient to provoke myelofibrosis and inflammation in diverse hematopoietic cell types in the BM niche, and that the MYC-S100A9 circuit is targetable in triple negative MPN.

## INTRODUCTION

Myelofibrosis (MF) is an aggressive myeloproliferative neoplasm (MPN) characterized by constitutional symptoms, cytopenias, splenomegaly, extramedullary hematopoiesis, bone marrow (BM) fibrosis, and a propensity for transformation to acute myeloid leukemia (AML)^1–3^. Approximately 85% of MF cases are driven by recurrent mutations in the *JAK2*, *CALR*, and *MPL* genes^4–9^ that result in constitutive activation of JAK/STAT, PI3K/AKT, and MEK/ERK signaling pathways and a robust production of pro-inflammatory cytokines, leading to chronic inflammation and BM fibrosis^10,11^. Accordingly, the FDA has approved the JAK2 inhibitors ruxolitinib, fedratinib, and pacritinib, which provide significant improvement in constitutional symptoms, splenomegaly and overall survival (OS) for MF patients^12–15^. The clinical benefits of JAK2 inhibition largely reflect anti-inflammatory effects, as current JAK2 inhibitors are not disease modifying agents and subgroups of MF patients have inferior response to these drugs, particularly in triple negative MF (TN-MF) cases that lack *JAK2/CALR/MPL* mutations^12–14,16–18^. Thus, the field is focused on developing new therapeutic strategies to improve clinical outcomes in MF, especially TN-MF^19^.

MYC is a basic-helix-loop-helix leucine zipper (bHLH-Zip) transcription factor that coordinates expression of genes controlling cell proliferation, survival, and metabolism^20–22^. Although MYC was originally identified as a transforming oncogene in lymphoid neoplasms^23–25^, our studies and those of others have shown that MYC has important oncogenic roles in several myeloid malignancies^26–34^. For example, copy number gain of the *MYC* gene, located on chromosome (chr) 8q24, frequently occurs across all myeloid malignancies via trisomy 8^35–39^, a genomic abnormality associated with increased MYC expression and adverse outcomes in AML and myelodysplastic syndrome (MDS)^26,27,34^. Further, as a direct downstream target of JAK/STAT signaling, MYC has been shown to play key roles in MPN cell survival and JAK2 inhibitor resistance^40–42^. However, whether MYC serves as an independent oncogenic driver in MPN is not known.

Here we report that *MYC* copy number gain frequently occurs in TN-MF, a subgroup of MPN with the worst clinical outcomes, and is associated with increased MYC levels in hematopoietic stem cells (HSCs). Further, using transgenic mouse models that conditionally overexpress low levels of MYC in HSCs, we demonstrate that MYC can provoke MF-like disease independent of JAK2 pathway mutations, that this requires upregulation of S100A9, an alarmin protein that plays a key role in inflammation and innate immunity^43–48^, and that the MYC-S100A9 axis represents a therapeutic vulnerability for MF

## RESULTS

### *MYC* copy number gain frequently occurs in TN-MF

To identify potential oncogenic drivers of TN-MF, we first performed cytogenetic analyses and targeted exome sequencing of 98 genes commonly mutated in myeloid malignancies in DNA from 584 MF patients who were identified in the Moffitt Cancer Center Total Cancer Care database as previously described^49^. Our MF cohort includes total 379 (65.0%) primary MF, 83 (14.2%) post-PV (polycythemia vera) MF, and 122 (20.9%) post-ET (essential thrombocythemia) MF (Figure 1A; Table S1), with the most common somatic mutation in *JAK2* followed by *ASXL1, TET2, CALR*, and *SRSF2* (Figures 1B-C and S1A). Consistent with previous reports^12–14,16–18^, TN-MF patients in our cohort showed significantly shorter leukemia free survival (LFS) and OS compared to *JAK2/CALR/MPL* mutant patients (Figures S1B-C). Notably, there was no significant difference in mutation profiles (other than JAK2 activating mutations) between TN *vs. JAK2/CALR/MPL* mutant MF (Figure S1A). However, we observed that trisomy 8 occurs more frequently in TN-MF *vs.* other subtypes (Figures 1D and S1D). Supporting a potential pathogenic role of trisomy 8 in TN-MF, patients with trisomy 8 had significantly shorter LFS (median LFS 21.4 *vs.* 32.9 years, p=0.0005) and OS (median OS 3.1 *vs.* 8.9 years, p=0.0011) (Figures 1E-F and S1E-F).

**Figure 1.**
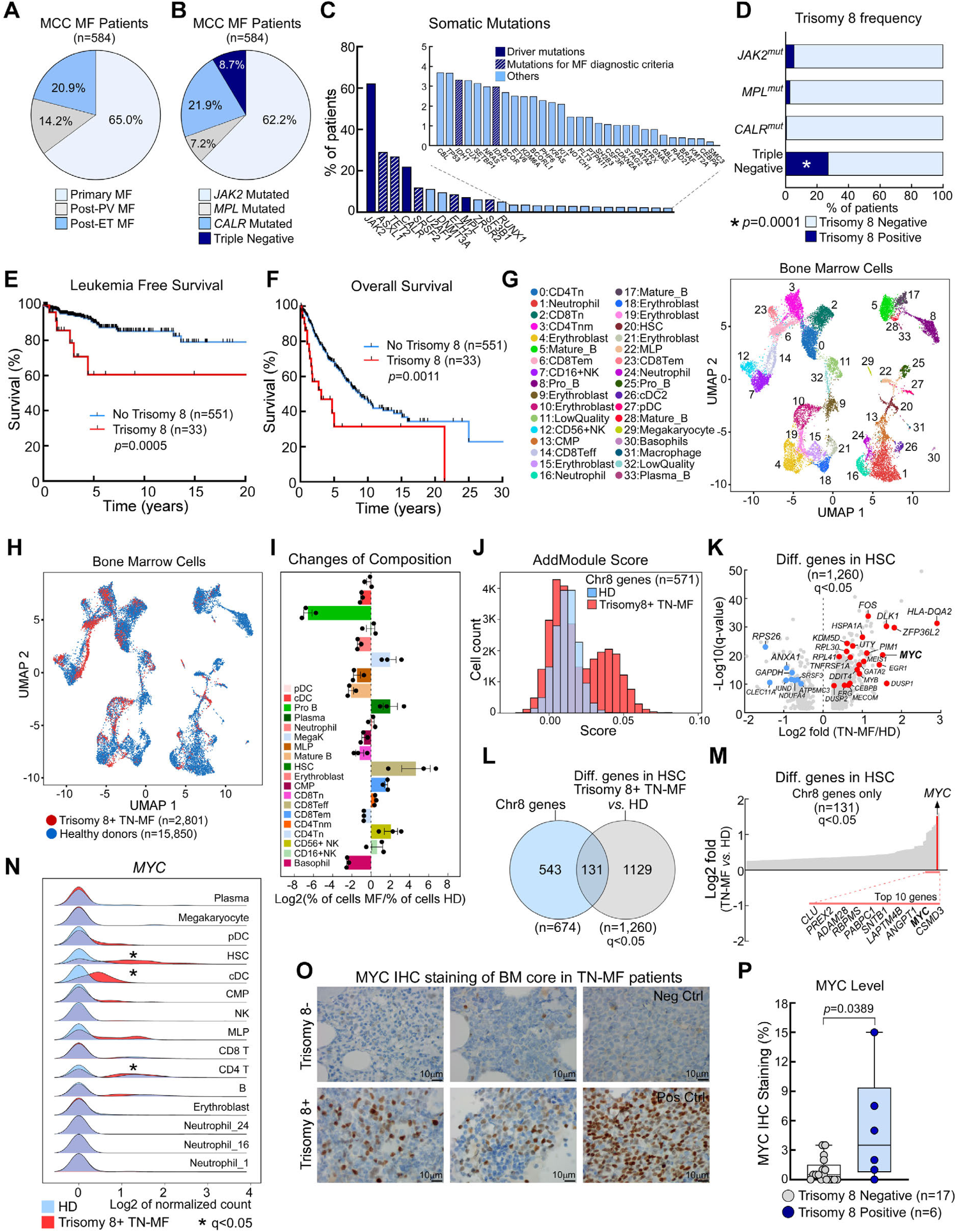
Trisomy 8 is associated with elevated MYC expression in BM HSCs of TN-MF patients. **A,** The proportion of primary, post-PV, and post-ET MF in Moffitt MF dataset (n=584). **B,** The frequency of *JAK2/MPL/CALR* somatic mutations in the Moffitt MF cohort. Demographic and laboratory profiles are described in Table S1. **C,** Bar plot shows percentages of recurrently mutated genes; solid blue, driver mutations; and stripes, mutations included in the major diagnostic criteria for MF based on the WHO classification. **D,** The percentages of trisomy 8 in each molecular subtype of MF. **E-F,** Kaplan-Meier (KM) curves showing the LFS and OS based on presence of trisomy 8 in MF. G-H, UMAP plots of hematopoietic single cells in BM of normal HD (n=3) *vs.* trisomy 8+ TN-MF (n=1). Demographic and laboratory profiles are described in Table S2. **I-J,** Comparison of BM major cell types (I) and AddModule Score (J) of HD *vs.* trisomy 8+ TN-MF. AddModule Score was calculated based on a total of 571 coding genes located on chr8 and that were detected in scRNA-seq analysis. **K,** Volcano plot showing a total of 1,260 genes that are differentially regulated (q<0.05) in HSCs of trisomy 8+ TN-MF *vs.* HD. **L,** Venn diagram of coding genes on chr8 (n=674) and genes with differential changes in HSCs. **M,** Ranked list of log2-fold change of 131 chr8 genes differentially regulated in HSCs of trisomy 8+ TN-MF. **N,** Ridgeline plots comparing *MYC* mRNA levels in BM cells for each major cell types of trisomy 8+ TN-MF (red) *vs.* HD (blue). **O,** MYC IHC staining of BM core biopsy samples from TN-MF patients. **P,** Percentages of MYC positive cells in trisomy 8 negative (n=17) *vs.* positive (n=6) TN-MF. Demographic and laboratory profiles of TN-MF patients included in MYC IHC analysis are shown in Table S4.

To screen oncogenic drivers in trisomy 8+ TN-MF, scRNA-seq analysis was performed comparing BM cells collected from healthy donors (HD) (n=3) and a treatment-naïve TN-MF patient with trisomy 8 (n=1) (Figures S1H-I; Table S2). Analysis of 18,651 cells (n=15,850 from HD; n=2,801 from the TN-MF patient) identified a total of 34 clusters that exhibit distinct gene expression profiles (Figures 1G and S1J-K). Distributions of these clusters were not significantly different between these two groups in the UMAP analyses (Figure 1H), indicating that hematopoietic cells in this TN-MF patient maintain gene expression profiles that are similar to normal cells. Cell component analyses, however, revealed expansion of HSCs, megakaryocytes, CD8^+^ T-cells, and CD56^+^ NK-cells in TN-MF, and reductions in multipotent lymphoid progenitors (MLPs), B-cells, and basophils (Figures 1I and S1L-M). In addition, the percentages of cells in S or G2/M phase were markedly increased in HSCs and common myeloid progenitors (CMPs) of trisomy 8+ TN-MF patient compared to HD (Figure S1N).

Copy number variation was also inferred using the scRNA-seq dataset. As expected, there was a distinct pattern of chr8 gain across major cell types (e.g., HSCs, CMPs, MLPs, erythroblasts, dendritic cells, and NK-cells) in trisomy 8+ TN-MF, but not in HD (Figures S1O-P). Of note, trisomy 8 was associated with bimodal distribution of AddModule score that was calculated based on mRNA expression levels of 571 chr8 genes detected in our scRNA-seq analysis (Figure 1J). These results suggest that the additional copy of chr8 increases levels of chr8 gene expression. Subsequent comparison of 674 coding genes on chr8 *vs.* 1,260 genes that are differentially regulated in HSCs of trisomy 8+ TN-MF (compared to HD) identified a total of 131 chr8 genes (Figures 1K-L). Among these, *MYC* ranked second based on mRNA levels (Figure 1M) and was preferentially upregulated in HSCs (Figure 1N). Further, the MYC pathway was more active in the majority of major cell types (including HSCs/CMPs) in trisomy 8+ TN-MF based on PROGENy analysis (Figure S1Q; Table S3). Importantly, a comparison of trisomy 8+ *vs.* trisomy 8-cells in the same TN-MF patient, revealed that *MYC* levels were also significantly higher in trisomy 8+ HSCs/CMPs (Figure S1R); thus, increased *MYC* levels in trisomy 8+ TN-MF patient cells do not result from individual variation, but instead reflect gene copy number.

These findings were validated using immunohistochemistry (IHC) staining. In particular, BM MYC protein levels were significantly higher in trisomy 8+ *vs.* trisomy 8-TN-MF cells (Figures 1O-P and S1G). Of note, we did not observe any significant difference in somatic mutation profiles, cytogenetics, or clinical parameters between MYC negative *vs.* positive cases that potentially contribute to different levels of MYC expression other than trisomy 8 (Figure S1S; Table S4). Collectively, these results support the hypothesis that elevated MYC expression driven by *MYC* copy number gain may play an oncogenic role in MF independent of JAK2 pathway mutations.

### MYC provokes an MPN that resembles MF *in vivo*

To assess the effects of MYC in HSCs and potential roles of MYC in MPN development, we established two independent transgenic mouse models: Mx1-Cre^+/-^;Rosa26^LSL-MYC/LSL-MYC^ and Scl-CreERT^+/-^;Rosa26^LSL-MYC/LSL-MYC^ that inducibly overexpress human *MYC* in HSCs following Cre enzyme-mediated removal of a *LoxP-stop-LoxP* transcriptional stop cassette (Figures 2A-C and S2A)^50^. MYC protein levels in the Mx1-Cre^+/-^;Rosa26^LSL-MYC/LSL-MYC^ mice were initially compared with those expressed in lymphomas of Eµ-*Myc* transgenic mice^51,52^ and in MYC10 VavP-*MYC* mice that express low levels of MYC and develop myeloid disease^53^. BM MYC levels in Mx1-Cre^+/-^;Rosa26^LSL-MYC/LSL-MYC^ mice were significantly lower than levels in Eµ-*Myc* B lymphoma cells (Figure 2D) but were similar to those in MYC10 BM cells (Figure 2E). These findings indicate that Mx1-Cre^+/-^;Rosa26^LSL-MYC/LSL-MYC^ mice express biologically relevant levels of MYC to study myeloid neoplasms^25,53^.

**Figure 2.**
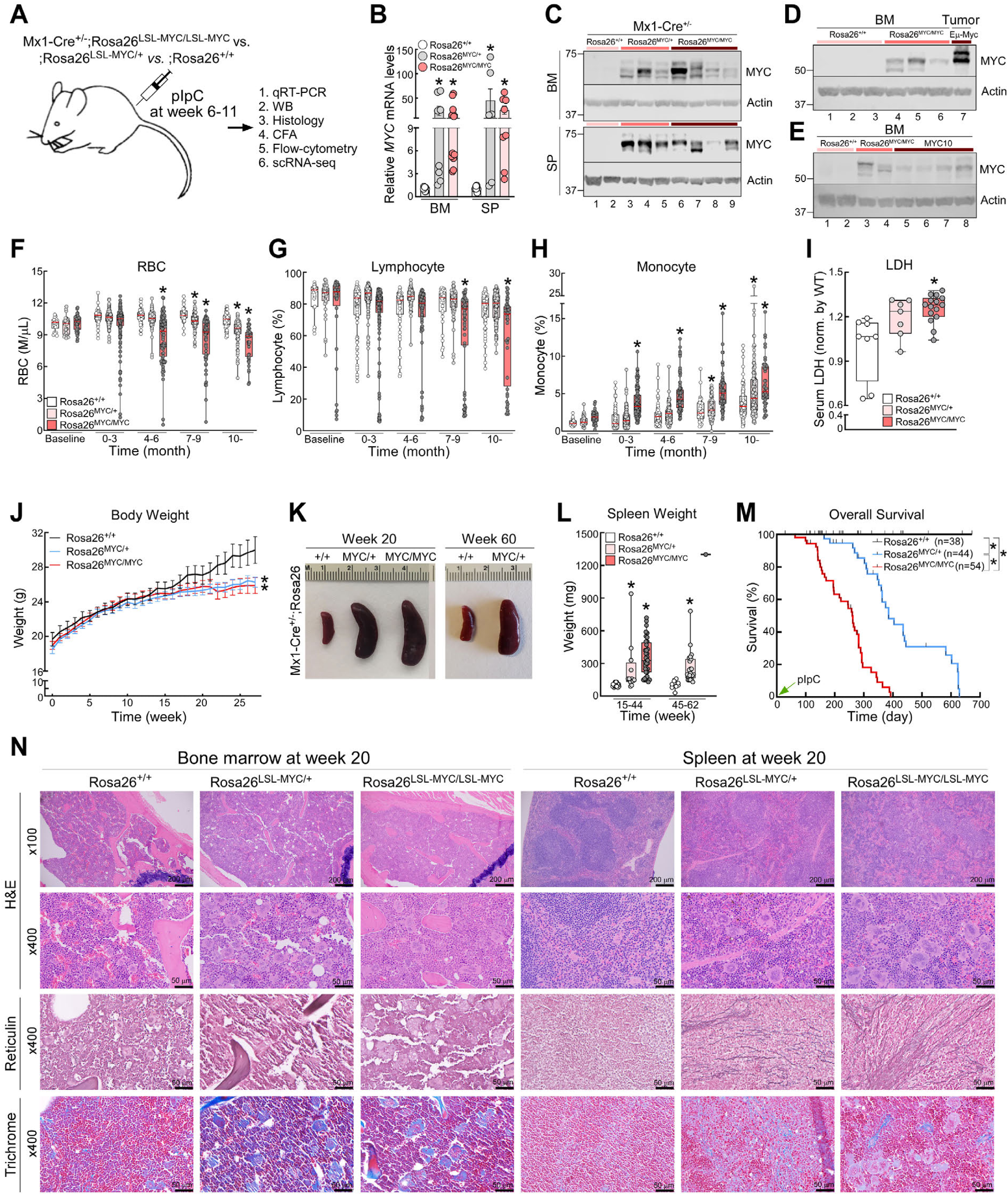
MYC overexpression in HSCs provokes MF. **A,** Schematic of mouse *in vivo* studies. **B-C,** MYC mRNA (B) and protein (C) levels in BM and spleen cells of Mx1-Cre^+/-^;Rosa26^+/+^, Mx1- Cre^+/-^;Rosa26^LSL-MYC/+^, and Mx1-Cre^+/-^;Rosa26^LSL-MYC/LSL-MYC^ mice, respectively, at 20 weeks post-pIpC treatment. **D-E,** Comparison of MYC protein levels in Mx1-Cre^+/-^;Rosa26^LSL-MYC/LSL-MYC^ BM cells *vs.* Eµ-*Myc* lymphoma cells (D) or VavP-MYC10 mice BM cells (E) at age 8 weeks. **F-H,** Peripheral blood (PB) CBC analyses. Baseline CBC was performed 1 week prior to pIpC injection. **I,** LDH activity in serum samples from each experimental group. **J,** Comparison of body weight changes of experimental groups. Baseline body weights were determined 1 week prior to pIpC treatment. **K-L,** Spleen weight at endpoints. Mx1-Cre^+/-^;Rosa26^+/+^ mice were sacrificed as control when Mx1- Cre^+/-^;Rosa26^LSL-MYC/+^ and Mx1-Cre^+/-^;Rosa26^LSL-MYC/LSL-MYC^ mice were at their endpoints. Endpoints criteria is described in “Animal studies” section. **M,** KM curves of OS, calculated from the date of pIpC treatment. **N,** H&E, reticulin, and trichrome stained images of BM (left 3 columns) and spleen (right 3 columns) at week 20 post-pIpC. Demographic and laboratory profiles are shown in Table S5. Box plots in (F-I, L) represent data from 38-54 mice in each group. Error bars in (B) indicate mean ±SEM of at least 6 independent mice. *, *P*<0.05 compared with control group.

The consequences of MYC overexpression in HSCs of Mx1-Cre^+/-^;Rosa26^LSL-MYC/LSL-MYC^ mice were evident following examination of peripheral blood (PB) of these mice, which displayed profound anemia and low grade monocytosis, as well as mild leukocytosis without significant changes in the neutrophil percentages and platelet counts (Figures 2F-H, S2B-D; Table S5). Mx1-Cre^+/-^;Rosa26^LSL-MYC/LSL-MYC^ mice also had significant increases in serum lactate dehydrogenase (LDH), slower weight gain, larger spleens, and shorter OS (median OS 258 *vs.* 385 days *vs.* not reached (NR) in MYC homozygous *vs.* heterozygous *vs.* wild type (WT), respectively, p<0.0001) (Figures 2I-M). Further, MYC homozygous mice had marked increases in atypical megakaryocytes^54,55^, extramedullary hematopoiesis, and fibrosis of the spleen, liver, and, to a lesser extent, BM (Figures 2N and S2E-G). Notably, there was no evidence of acute leukemia (e.g., increased blasts), MDS (e.g., dysplastic cells) or lymphoma (e.g., bulky lymph nodes). An extensive comparison of MYC heterozygous *vs.* homozygous mice revealed that both develop the same disease phenotypically, with heterozygous mice having delayed disease onset and lower disease burden (Figures 2F-N and S2B-G).

In independent experiments using Scl-CreERT^+/-^;Rosa26^LSL-MYC/LSL-MYC^ mice, we observed the same phenotypes as Mx1-Cre^+/-^;Rosa26^LSL-MYC/LSL-MYC^ mice, but with slower disease onset and progression (Figures S2H-T; Table S6). Collectively, these results indicate that modest levels of MYC overexpression in HSCs are sufficient to provoke a chronic myeloid neoplasm that is phenotypically and pathologically most similar to primary MF based on WHO diagnostic criteria^56–58^.

### MYC increases HSCs and myeloid progenitors with limited self-renewal capacity

Experiments characterizing hematopoietic sub-populations of Mx1-Cre^+/-^;Rosa26^LSL-MYC/LSL-MYC^ mice revealed that MYC promotes expansion of HSCs, multipotent progenitors (MPPs), myeloid progenitors such as CMPs, granulomonocytic progenitors (GMPs), and megakaryocyte erythrocyte progenitors (MEPs), as well as Gr-1^+^/CD11b^+^ mature myeloid cells. In contrast, there were reductions in the percentages of B- and T-lymphocytes in the BM and spleen (Figures 3A-C and S3A-C). Although MYC increased the percentages of HSCs and progenitors in the Lin^-^ population, there was no significant change in the proportion of individual components (i.e., LT-HSC, ST-HSC, MPP2, and MPP3) and there was no significant lineage bias in myeloid progenitors (Figures S3B-C). A similar expansion of HSCs/MPPs without changes in the proportion of individual components was observed in Scl-CreERT^+/-^;Rosa26^LSL-MYC/LSL-MYC^ and Scl-CreERT^+/-^;Rosa26^LSL-MYC/+^ mice compared to control mice (Figure S3D). Notably, MYC homozygous progenitors had increased monocytic colony forming units (M-CFU) and granulomonocytic CFU (GM-CFU) in BM cells and all CFU in spleen cells, but these displayed limited self-renewal capacity ex vivo (Figures 3D-E and S3E-J); thus, this MYC-driven myeloid neoplasm is not acute leukemia. Conversely, in myeloid progenitors isolated from the BM of Rosa26-CreERT2^+/-^;*Myc*^fl/fl^ mice, *Myc* loss provoked marked reductions in CFU, specifically in GM-CFU (Figure S3K-M).

**Figure 3.**
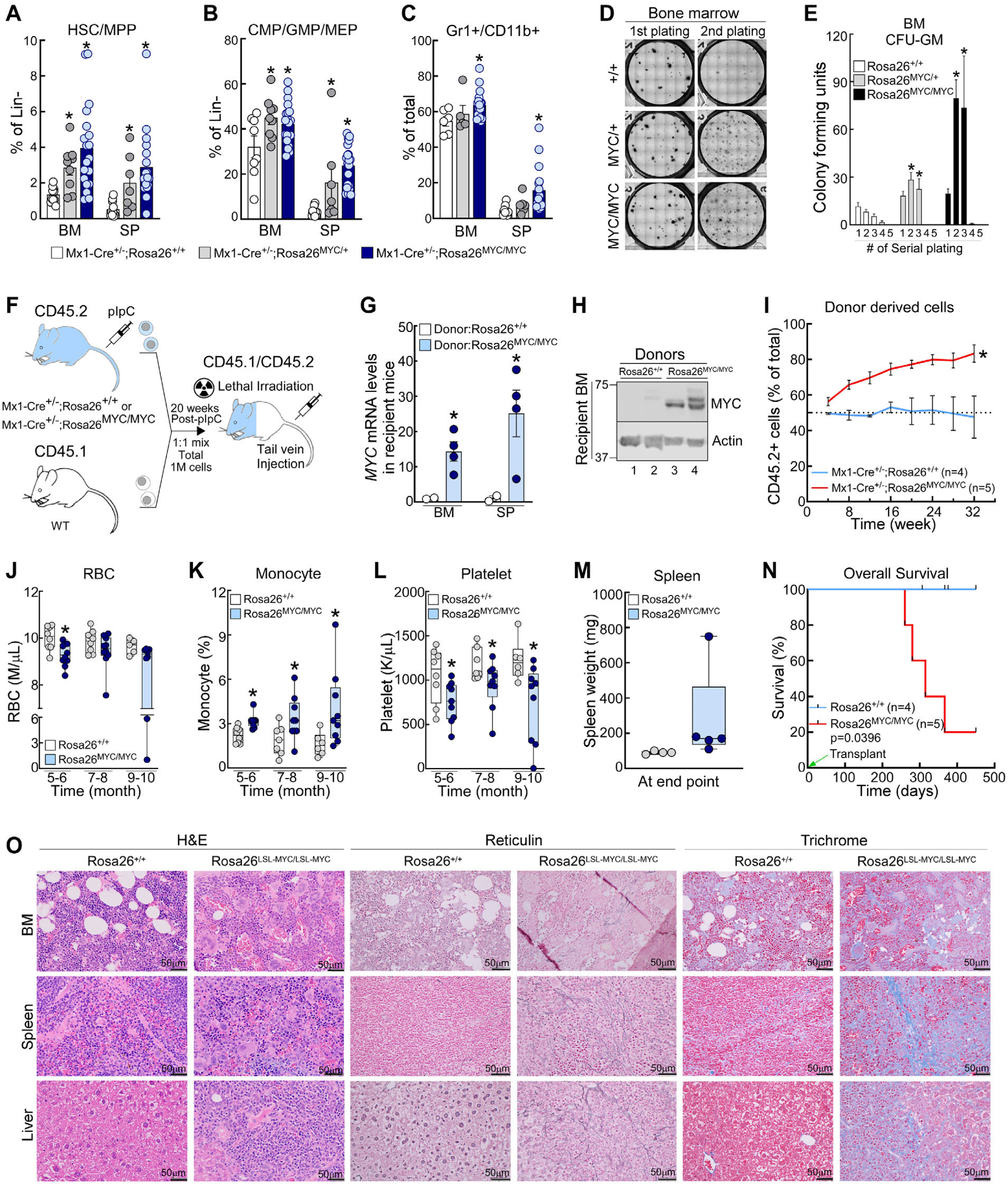
Forced MYC expression provokes expansion of HSCs and myeloid progenitors with limited self-renewal capacity. **A-C,** The percentages of HSCs/MPPs (A), myeloid progenitors (B), and Gr1^+^/CD11b^+^ mature myeloid cells (C) in each experimental group. **D-E,** Serial colony forming assays using primary BM cells collected from Mx1-Cre^+/-^;Rosa26^+/+^, Mx1-Cre^+/-^;Rosa26^LSL-MYC/+^ and Mx1-Cre^+/-^;Rosa26^LSL-MYC/LSL-MYC^ mice at 20 weeks post-pIpC. **F,** Competitive transplant using BM cells from CD45.2^+^ Mx1-Cre^+/-^;Rosa26^+/+^ *vs.* Mx1-Cre^+/-^;Rosa26^LSL-MYC/LSL-MYC^ mice. BM cells were harvested from these mice 20 weeks post-pIpC injection, mixed 1:1 with CD45.1^+^ WT BM cells, and a total of 1 million cells were injected via tail vein into lethally irradiated CD45.1^+^/CD45.2^+^ recipient mice. **G-H,** MYC mRNA (G) and protein (H) levels in BM and spleen cells from recipient mice at endpoints. **I,** Percentages of CD45.2^+^ cells were assessed by serial PB flow cytometry analyses. **J-L,** CBC analyses at indicated times following transplantation. **M,** Spleen weight at endpoints. **N,** KM curves of OS, calculated from the date of transplant. **O,** H&E, reticulin, and trichrome stained images of BM, spleen, and liver. Mice transplanted with Mx1- Cre^+/-^;Rosa26^+/+^ BM cells were sacrificed as controls when mice transplanted with Mx1-Cre^+/-^;Rosa26^LSL-MYC/LSL-MYC^ BM cells approached their endpoints. Error bars in (A-C, E, G, I) indicate mean ±SEM of at least 4 independent mice. *, *P*<0.05 compared with control group.

In further studies, CD45.1^+^/CD45.2^+^ mice were transplanted with CD45.2^+^ Mx1-Cre^+/-^;Rosa26^LSL-MYC/LSL-MYC^ and CD45.1^+^ WT *vs.* CD45.2^+^ Mx1-Cre^+/-^;Rosa26^+/+^ and CD45.1^+^ WT BM cells that are mixed at 1:1 ratio (Figure 3F). In these competitive transplant studies, the proliferative advantage associated with MYC overexpression was maintained in vivo (Figure 3G-I). Further, the transplanted MYC homozygous BM cells provoked a phenotypically indistinguishable disease in recipient mice and reduced their OS (median OS 315 days in Mx1-Cre^+/-^;Rosa26^LSL-MYC/LSL-MYC^ *vs.* NR in control transplanted group, p=0.0396) (Figures 3J-O; Table S7), indicating that the transplanted cells were sufficient to induce an MF-like chronic myeloid malignancy in the normal BM niche.

### MYC-driven expansion and proliferation of HSCs and myeloid progenitors is independent of JAK/STAT signaling

To identify the mechanism by which MYC drives MF, scRNA-seq analysis was performed comparing BM cells harvested from Mx1-Cre^+/-^;Rosa26^LSL-MYC/LSL-MYC^ *vs.* Mx1-Cre^+/-^;Rosa26^+/+^ mice at 20 weeks following pIpC injection (Figure S4A). By analyzing 25,232 cells (n=13,552 from control and n=11,680 from Mx1-Cre^+/-^;Rosa26^LSL-MYC/LSL-MYC^ mouse), we identified a total of 33 clusters and 23 major cell types (Figures 4A and S4B-D). As in analyses of human TN-MF and healthy human donors (Figure 1H), distributions of these clusters were not significantly different between MYC homozygous vs. control mice (Figure 4B). Cell component analyses revealed that the percentages of HSCs, myeloid progenitors, and mature myeloid cells, including monocytes and macrophages, were increased, whereas erythroblasts, B- and T-lymphocytes were reduced in MYC homozygous mice (Figures 4C-D). Notably, MYC homozygous mouse has an increased percentages of cells in S or G2/M phase, specifically in HSCs, myeloid progenitors, monoblasts, and monocytes (Figure S4E). These findings are consistent with the results from complete blood counts (CBC) with differential and flow-cytometry (Figures 2H, S2O, 3A-C, 3K, S3C, S3E, S3I, and S3M), further supporting the role of MYC in driving myeloid proliferation.

**Figure 4.**
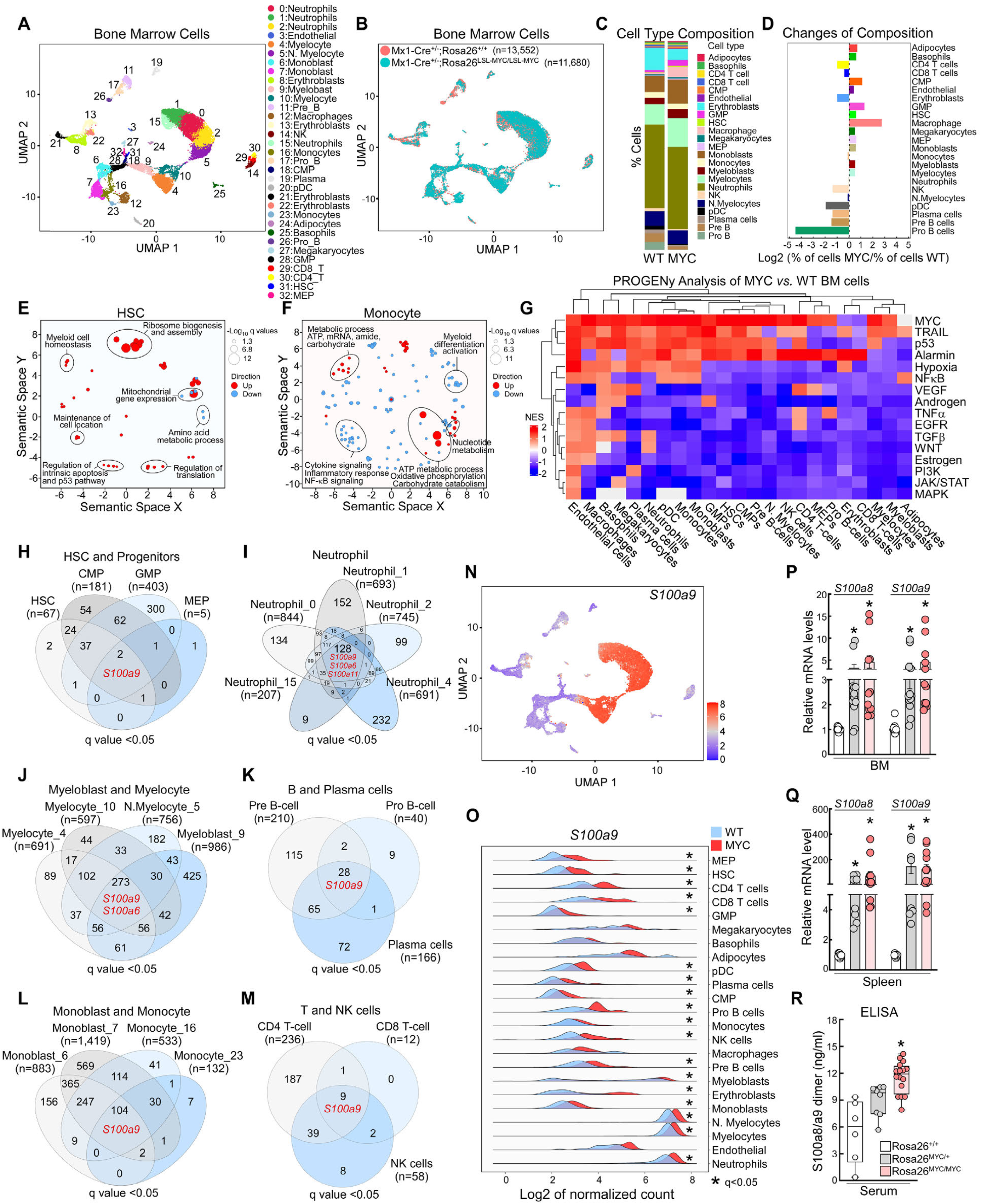
Characterization of MYC-driven MF at the single cell level. **A-B,** UMAP plots of single hematopoietic cells in BM of Mx1-Cre^+/-^;Rosa26^+/+^ (control) *vs.* Mx1-Cre^+/-^;Rosa26^LSL-MYC/LSL-MYC^ (MYC) mice at 20 weeks post-pIpC injection. **C-D,** Bar plots of percentages of major cell types in control *vs.* MYC mouse. **E-F,** Semantic plots of HSCs (E) and monocytes (F). Upregulated and downregulated pathways in MYC *vs.* control cells are highlighted in red and blue, respectively. **G,** PROGENy analysis comparing MYC *vs.* control cells. Individual column and row are major cell types and pathways known to contribute to MPN pathogenesis, respectively. **H-M,** Venn diagram of MYC-controlled, differentially expressed genes by scRNA-seq (log2fold change>0.25, q<0.05) in major cell types. **N,** Expression levels of *S100a9* in individual cells are projected onto the UMAP space. **O,** Ridgeline plots comparing *S100a9* mRNA levels in MYC (red) vs. control (blue) BM cells in each major cell type. **P-Q,** qRT-PCR assays were used to assess the levels of *S100a8/a9* mRNA in BM and spleen cells. **R,** ELISA was used to assess the levels of S100a8/a9 heterodimer in the serum at their end points. Error bars in (P-Q) indicate mean ±SEM of at least 6 independent mice. Box plot in (R) represents data from 6-16 mice in each group. *, *P*<0.05 compared with control group.

To assess MYC-driven transcriptomic changes in each cellular component, we performed differential gene expression analysis followed by semantic analysis in individual major cell types (Figures S4F-G; Table S8). Many of the well-known MYC target pathways (e.g., ribosome biogenesis, regulation of translation, p53 and intrinsic apoptosis^22^) were significantly activated in MYC homozygous HSCs (Figure 4E). In mature myeloid cells such as monocytes, MYC activated genes involved in metabolic processes, including oxidative phosphorylation and carbohydrate catabolism, and suppressed genes involved in myeloid differentiation (Figure 4F). Unexpectedly, MYC also suppressed cytokine signaling and inflammatory response pathways (Figure 4F). Indeed, PROGENy analysis revealed that the JAK/STAT, PI3K/AKT, and MAPK pathways as well as TNF-α and TGF-β signaling that play critical roles in chronic inflammation in the *JAK2/CALR/MPL* mutant MPN^10,11^ are inactive in most of the major cell types that overexpress MYC, whereas p53 and alarmin pathways are substantially activated by MYC (Figure 4G; Table S9). These findings were validated by immunoblotting that showed a marked increase in p53 in MYC overexpressing BM and spleen cells *vs*. control cells but no significant differences in the levels of phosphorylated Stat3, Akt, and Erk1/2 (Figures S4H-I).

Further underscoring a negligible role of the JAK pathway in MYC-driven MF, ruxolitinib treatment had minimal effects on the levels of phosphorylated Stat3, Akt, and Erk1/2 as well as apoptosis in homozygous MYC BM cells, although the same ruxolitinib concentrations induced robust cell death and/or downregulated levels of phospho-Stat3, -Akt, and -Erk1/2 in the *JAK2^V617F^*mutant SET-2 and HEL MPN cells (Figures S4J-M). Collectively, these findings suggest that MYC provokes MF independent of the JAK pathway.

### MYC induces S100A9-mediated chronic inflammation

Because the alarmin pathway upregulated by MYC (Figures 4G) can induce inflammation independent of the JAK/STAT pathway, we next focused on defining MYC targets that play critical roles in the alarmin pathway. Among genes that are differentially regulated by MYC in individual clusters (Figures S4F-G), increased MYC was accompanied by *S100a9* upregulation in most of the major cell types as well as increased expression of *S100a8* and *ASC* in several of the cell types (Figures 4H-O, S4N-Q; Table S8). These findings were validated by qRT-PCR, ELISA, and immunoblotting (Figures 4P-R, S4H-I, S4R). Supporting the clinical relevance of S100A9 in MF pathogenesis, *S100A9* levels in CD34^+^ BM cells were significantly elevated in MF patients *vs.* HD regardless of *JAK2^V617F^* mutation status (Figures 5A-B)^59,60^. Importantly, in the TN-MF cohort, cells with *MYC* expression showed significantly higher S100A9 levels in their BM cells compared to cells not expressing *MYC* (Figures 1O-P and 5C-E). In addition, *S100A8/A9* levels in megakaryocytes, neutrophils (i.e., cluster 16), as well as *ASC* levels in CMPs, neutrophil, NK-, T-, and B-cells were markedly increased in trisomy 8+ TN-MF compared to HD (Figures S5A-C). Finally, MYC induced a marked increase in cleaved caspase-1 and dimerized ASC (Figures S4H-I and S4S), which are hallmarks of activation of S100a8/a9-dependent inflammation^61^. Collectively, these observations suggest that increased MYC in HSCs induces S100a9-mediated inflammation.

**Figure 5.**
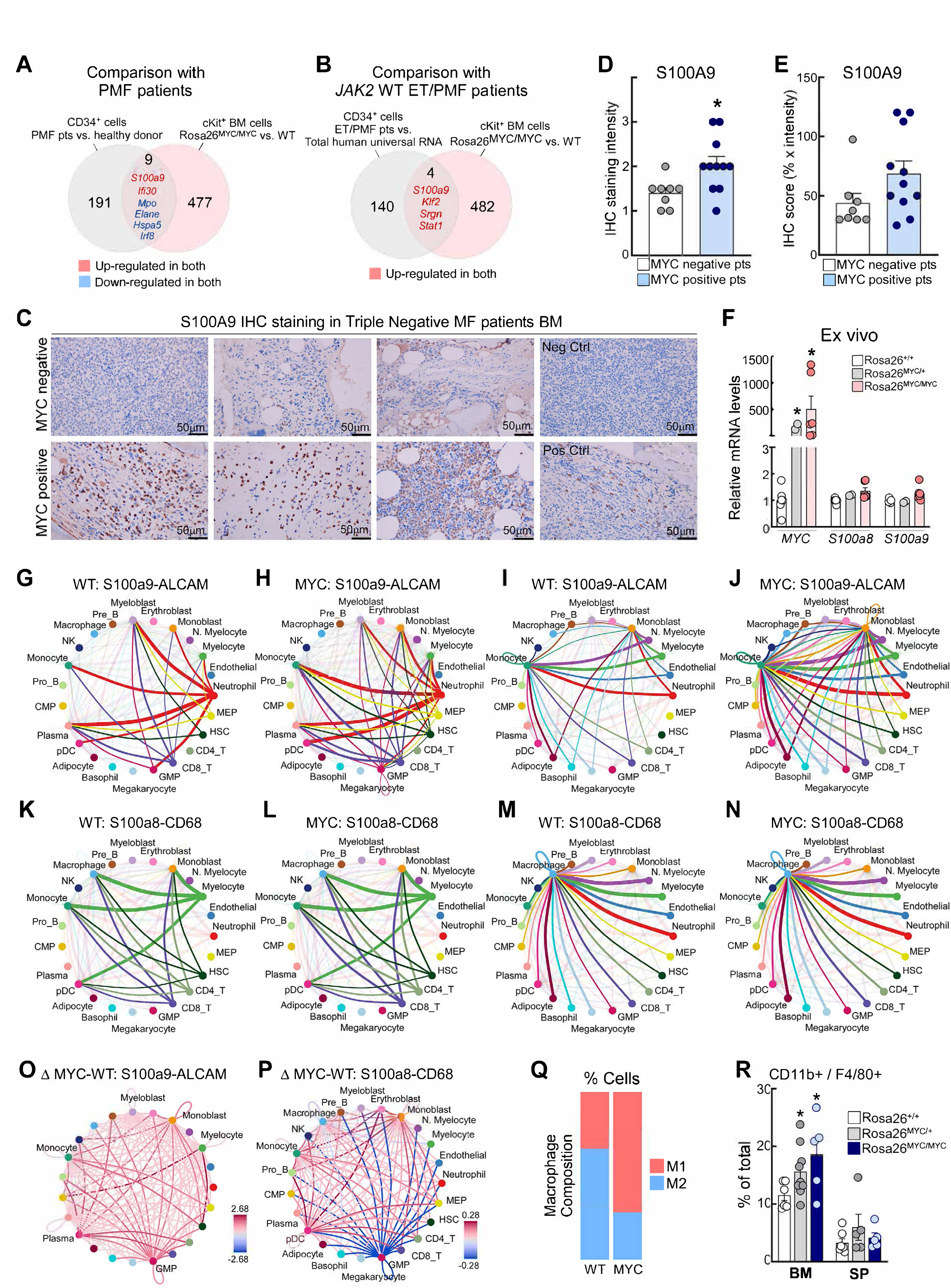
MYC-directed changes in cell-cell interactions in the BM niche. **A,** Comparison of CD34^+^ BM cells of *JAK2^V617^* mutant MF patients (n=9) *vs.* HD (n=6) identified 200 differentially regulated genes. Among these, 9 are also regulated by MYC, including *S100A9*. **B,** A total of 144 genes were identified as upregulated in CD34^+^ BM cells from *JAK2* WT MF patients (n=5) vs. control (total human universal RNA), and *S100A9* is one of 4 genes that is also increased by MYC. **C-E,** IHC staining analysis of BM demonstrates increased levels of S100A9 protein in TN-MF patients with positive MYC expression vs. patients lacking MYC expression in their BM. **F,** Levels of *MYC, S100a8*, and *S100a9* in *ex vivo* cultured (for 9∼10 days) primary BM cells harvested from the indicated mice. **G-P,** Network plots of ligands (S100a8, S100a9) and receptors (ALCAM, CD68) interactions. S100a8/a9 signals originating from HSCs, GMPs, MEPs, myelocytes, neutrophils, and T-cells in WT (G, K) and MYC (H, L) mice, and S100a8/9 signals coming to monoblasts, monocytes, or macrophages in WT (I, M) and MYC (J, N) mice are presented in different colors. Thicker line in **G-N,** indicates a more frequent interaction between cell types and differences in intensity of interactions are shown in (O-P); higher and lower signal activity in MYC cells (*vs.* WT) are in red and blue, respectively. **Q,** Percentages of M1 *vs.* M2 macrophages in MYC *vs.* WT mice based on scRNA-seq analysis. R, Percentages of macrophages (CD11b^+^ and F4/80^+^ cells) in BM and spleen were assessed by flow cytometry at 20 weeks post-pIpC injection as indicated. Error bars in (D-F, R) indicate mean ±SEM of at least 3 independent samples or assays. *, *P*<0.05 compared with control group.

### A complex network of S100A9-mediated inflammatory signaling is manifest in MYC-driven MF

Interestingly, MYC-directed S100a8/a9 upregulation was completely lost when cells were cultured ex vivo (Figure 5F), indicating that an intact BM niche is critical for the MYC-driven inflammation. Notably, following the induction of MYC expression there were several marked changes in cell-cell interactions. First, a significant portion of S100a8/a9 signal was derived from cells in the granulocyte lineage (i.e., myeloblasts, myelocytes, neutrophils) (Figures 4N-O, S4N, S4P, 5G-N, S5D-F; Table S10). However, for HSCs, myeloid progenitors (i.e., CMPs, GMPs, MEPs), and lymphocytes (i.e., T-, B-, NK cells) that are normally less inflammatory, MYC expression also resulted in elevated levels of *S100a9*, thus affecting many other cell types expressing S100a9 receptors such as ALCAM^62^ (Figures 5G-J). Second, even though MYC did not affect the frequency or pattern of interactions of some of the ligand-receptor pairs such as S100a8/a9-CD68 (Figures 5K-N, S5E-F), the intensity of S100a8/a9-mediated interactions was markedly increased in cells with elevated MYC levels (Figures 5O-P, S5G). Third, cells of myelomonocytic lineage (i.e., myeloblasts, myelocytes, monoblasts, monocytes, and macrophages) in MYC homozygous mice were most affected by S100a8/a9 signaling (Figures 5G-N, S5D-F). Accordingly, levels of the inflammatory marker ASC were significantly increased in these cell types in MYC homozygous mice (Figures S4O, S4Q) and there was a marked expansion of monocytes and macrophages (Figures 2H, S2O, 3C-E, 3K, S3E, S3I, and S3M), especially M1 macrophages (Figures 5Q-R) that are known to be expanded by activation of S100A9 and CD68 and play pivotal roles in MF pathogenesis^63–65^. Finally, TNFα- and CSF-1-mediated cell-cell interactions were downregulated in MYC homozygous cells (Figures S5H-L; Table S10), further supporting negligible roles for the JAK2 pathway in this MYC-driven model of TN-MF. Collectively, these findings suggest that MYC provokes a unique inflammatory network that is mediated by S100A8/A9 within the BM niche.

### *S100a9* contributes to development of MYC-driven MF

To test whether S100a9 upregulation is required for MYC-driven MF development, we generated Mx1-Cre^+/-^;Rosa26^LSL-MYC/LSL-MYC^;*S100a9*^-/-^ mice and compared them with Mx1-Cre^+/-^;Rosa26^LSL-MYC/LSL-MYC^ mice (Figure 6A). MYC levels in BM and spleen cells of these two cohorts were similar and were significantly higher than MYC levels in WT or *S100a9*^-/-^ mice (Figures 6B-C). Conversely, S100a9 (in BM and spleen), as well as S100a8/a9 heterodimers (in serum) were undetectable in both *S100a9*^-/-^ and Mx1-Cre^+/-^;Rosa26^LSL-MYC/LSL-MYC^;*S100a9*^-/-^ mice (Figures 6C-D). *S100a9* deficiency impaired MYC-induced anemia and monocytosis, and increased platelet counts (Figures 6E-F, S6A-C; Table S11), indicating that S100a9 contributes to MYC-induced MF. Further, Mx1-Cre^+/-^;Rosa26^LSL-MYC/LSL-MYC^;*S100a9*^-/-^ mice displayed significant improvements in weight gain, reduced spleen size, and prolonged OS compared to Mx1-Cre^+/-^;Rosa26^LSL-MYC/LSL-MYC^ mice (median OS NR *vs.* 225 days, p=0.0345) (Figures 6G-H, S6D; Table S11). Flow cytometric analyses revealed that *S100a9* loss in Mx1-Cre^+/-^;Rosa26^LSL-MYC/LSL-MYC^ mice (i) substantially reduced the percentages of HSCs, myeloid progenitors, and macrophages in BM and/or spleen; (ii) significantly reduced the percentages of Gr1^+^/CD11b^+^ myeloid cells in spleen; and (iii) restored the percentages of B220^+^ and CD3^+^ cells close to normal levels in spleen (Figures 6I-N). Notably, *S100a9* deficiency alone did not affect normal hematopoiesis, as CFU in *S100a9*^-/-^ progenitors were similar to those of WT mice (Figures S6E-G). Finally, loss of *S100a9* effectively suppressed MYC-driven megakaryocytic atypia, fibrosis, and extramedullary hematopoiesis (Figures 6O, S6H). However, the size of spleens and the percentages of HSCs/progenitors in Mx1-Cre^+/-^;Rosa26^LSL-MYC/LSL-MYC^;*S100a9*^-/-^ mice were still higher than those of WT or Mx1-Cre^+/-^;Rosa26^+/+^ mice (Figures 6G and 6I-J), and there was residual megakaryocytic atypia and extramedullary hematopoiesis in Mx1-Cre^+/-^;Rosa26^LSL-MYC/LSL-MYC^;S100a9^-/-^ mice (Figures 6O, S6H). Thus, S100a9 plays a major role in the pathogenesis of MYC-driven MF, but additional targets downstream of MYC might also contribute to myeloid proliferation and MF pathogenesis.

**Figure 6.**
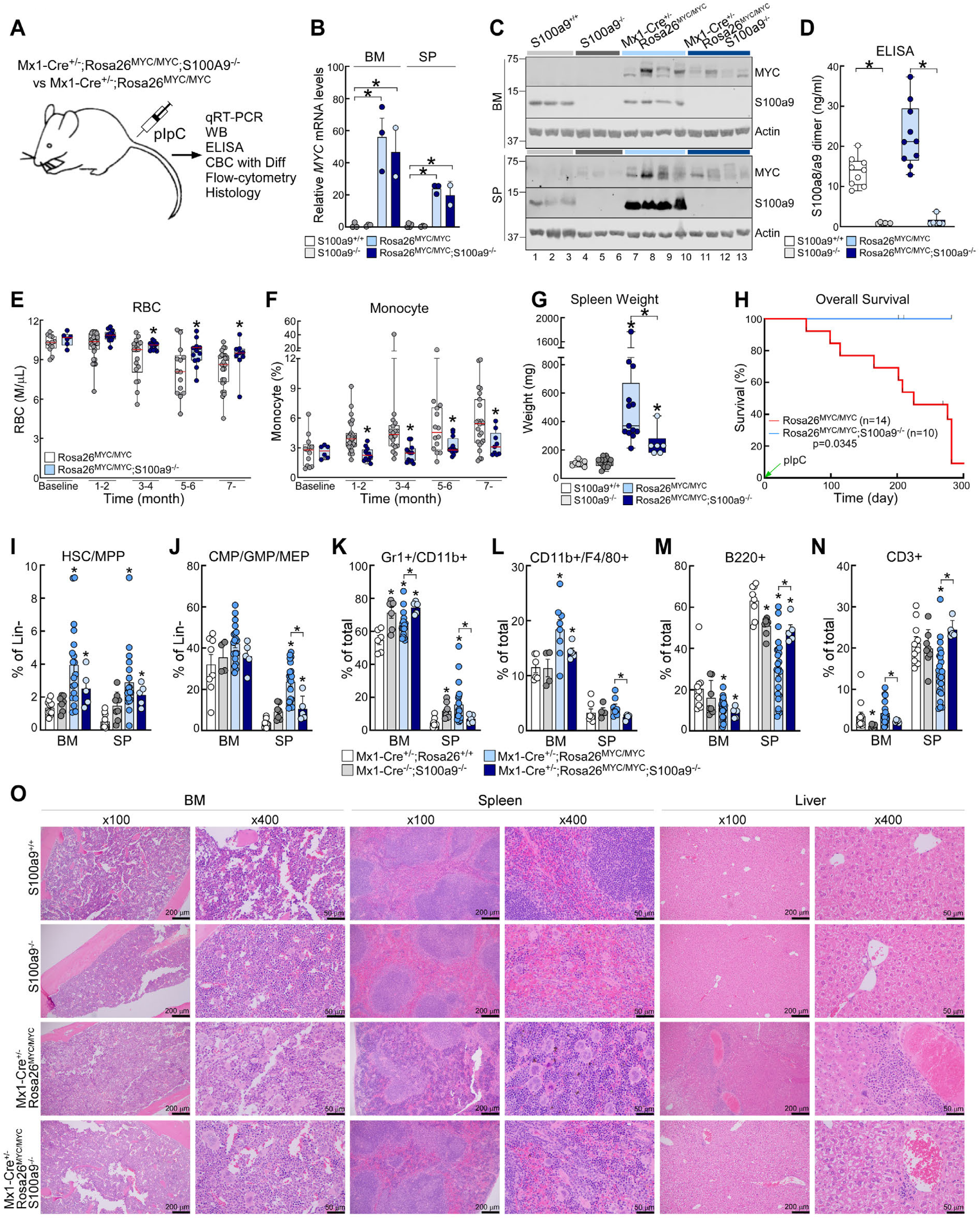
*S100a9* loss impairs MYC-driven MF. **A,** Schematic of in vivo mouse studies. **B-D,** Levels of MYC, S100a9, and S100a8/a9 heterodimers in BM, spleen, and serum, respectively, in *S100a9*^+/+^, *S100a9*^-/-^, Mx1-Cre^+/-^;Rosa26^LSL-MYC/LSL-MYC^, and Mx1-Cre^+/-^;Rosa26^LSL-MYC/LSL-MYC^;*S100a9*^-/-^ mice. *S100a9*^+/+^ and *S100a9*^-/-^ mice were sacrificed as age matched controls when Mx1-Cre^+/-^;Rosa26^LSL-MYC/LSL-MYC^ mice were at their endpoints. Mx1-Cre^+/-^;Rosa26^LSL-MYC/LSL-MYC^;*S100a9*^-/-^ mice were sacrificed at ∼32 weeks post-pIpC treatment, which is equivalent to the median OS of Mx1- Cre^+/-^;Rosa26^LSL-MYC/LSL-MYC^ mice (see panel H). **E-F,** PB CBC analyses. Baseline CBC was performed 1 week prior to pIpC injection. **G,** Comparison of spleen weight. Mice were sacrificed as described in panels (B-D). **H,** KM curves of OS. **I-N,** Percentages of HSCs/MPPs (I), myeloid progenitors (J), Gr1^+^/CD11b^+^ myeloid cells (K), macrophages (L), B-cells (M), and T-cells (N) in each group. **O,** H&E stained images of BM, SP, and liver at week 32 post-pIpC injection. Demographic and laboratory profiles are described in Table S11. Actin blots in (C) were from the same samples run on different gels. Box plots in (D-G) represent data from at least 6 mice in each group. Experiments in (I-N) were performed simultaneously with experiments in Figure 3A-C and Figure S3B, thus, control and some of experimental groups are shared among these experiments. Error bars in (B, I-N) indicate mean ±SEM of at least 2 independent assays. *, *P*<0.05 compared with control group.

### Enforced S100a9 expression is sufficient to provoke MF phenotypes

To test whether S100a9 overexpression would phenocopy the effects of MYC in driving MF, we assessed the phenotypes manifest in S100a9 transgenic (S100a9Tg) mice that overexpress *S100a9* under the control of the H2-K promoter (Figure S6I)^66^. Although there was no significant difference in *S100a9* mRNA levels in BM of S100a9Tg and WT mice (Figure S6J), most likely due to the abundance of neutrophils that express very high levels of *S100a9* in BM (Figures 4A and 4N), there were significantly higher levels of S100a9 mRNA and protein in the spleen of S100a9Tg mice (Figures S6K-L). At age >12 months, serum levels of S100a8/a9 dimers in S100a9Tg mice were also significantly elevated compared to age-matched control mice (Figure S6M). There was no positive or negative correlation between S100a9 and MYC protein levels in S100a9Tg (Figure S6L), indicating S100A9 does not affect MYC expression.

Notably, S100a9 overexpression alone was sufficient to provoke anemia, lymphopenia, monocytosis, neutrophilia, and thrombocytopenia in ∼60% of S100a9Tg mice, but these phenotypes only became prominent after 11 months of age (Figures S6N-R; Table S12). Forced expression of S100a9 also induced marked splenomegaly and significantly reduced OS (median OS 371 days *vs.* NR, p=0.0104) (Figure S6S-U). Further, S100a9 promoted expansion of HSCs, MPPs, and mature myeloid cells, and provoked decreases in B- and T-cells in BM and spleen (Figure S6V). Finally, S100a9 overexpression induced marked increase in megakaryocytic atypia, extramedullary hematopoiesis, and fibrosis of BM, spleen, and liver (Figure S6W). Collectively, these data confirm that S100A9 plays a pivotal role in MYC-driven MF.

Based on these findings, we assessed effects of pharmacologic inhibition of S100a9 in vivo using Tasquinimod, a small molecule that binds to S100a9 and prevents its interaction with its receptors^67,68^ (Figure S7A). Phenocopying some of the effects of *S100a9* deletion, treatment of Mx1-Cre^+/-^;Rosa26^LSL-MYC/LSL-MYC^ mice with Tasquinimod (i) increased platelet counts, (ii) reduced spleen size, and (iii) partially reversed MYC-driven expansion of myeloid progenitors in spleen and of macrophages in BM (Figures S7B-H; Table S13). Further, Mx1-Cre^+/-^;Rosa26^LSL-MYC/LSL-MYC^ mice treated with Tasquinimod showed reduced megakaryocytic atypia, fibrosis, and extramedullary hematopoiesis (Figure S7I). Nonetheless, Tasquinimod failed to improve OS (median OS 251 *vs.* 208 days, p=0.7715) (Figure S7J). Instead, prolonged treatment with Tasquinimod (>3 months) provoked profound anemia and neutrophilia (Figures S7B, S7D), which were not seen in Mx1-Cre^+/-^;Rosa26^LSL-MYC/LSL-MYC^;*S100a9*^-/-^ mice (Figures 6E, S6B). These changes, which are likely off-target effects of Tasquinimod, may explain the failure of this agent to phenocopy *S100a9* knockout and significantly improve OS.

### Targeting of the MYC-S100A9 circuit with a small molecule antagonizing MYC

In further studies, we examined the impact of directly targeting MYC using MYCi975, a small molecule that directly binds to the HLH domain of MYC, reducing MYC protein stability and transcriptional activity^69^. MYCi975, at concentrations that downregulate MYC protein in *JAK2^V617F^*mutant MPN cells and MYC-dependent AML cells, induced robust PARP cleavage and apoptosis in primary BM cells from Mx1-Cre^+/-^;Rosa26^LSL-MYC/LSL-MYC^ mice ex vivo (Figures S7K-O), indicating that this model of TN-MF is addicted to MYC.

To test whether MYC inhibition can suppress disease progression in vivo, lethally irradiated CD45.1^+^/CD45.2^+^ WT mice were transplanted with BM cells harvested from CD45.2^+^ Mx1-Cre^+/-^;Rosa26^LSL-MYC/LSL-MYC^ mice that were treated with pIpC 20 weeks prior to the transplant (Figure 7A). Engraftment of donor-derived CD45.2^+^ BM cells was confirmed by flow-cytometry at 4 weeks post-transplant, and mice were then randomized to either MYCi975 or vehicle treatment and monitored for disease progression (Figure 7A). MYCi975 effectively reduced MYC and S100a8/a9 levels in BM and spleen cells in vivo (Figure 7B). Consistent with these results, MYCi975 slowed disease progression and even caused disease regression in some mice (Figure 7C). Further, MYCi975 effectively suppressed MYC-driven MF phenotypes (Figures 7D-F, S7P; Table S14). Finally, these changes were associated with significantly improved OS (median OS NR *vs.* 252 days in MYCi975- *vs.* vehicle-treated group, respectively, p=0.0109) (Figure 7G).

**Figure 7.**
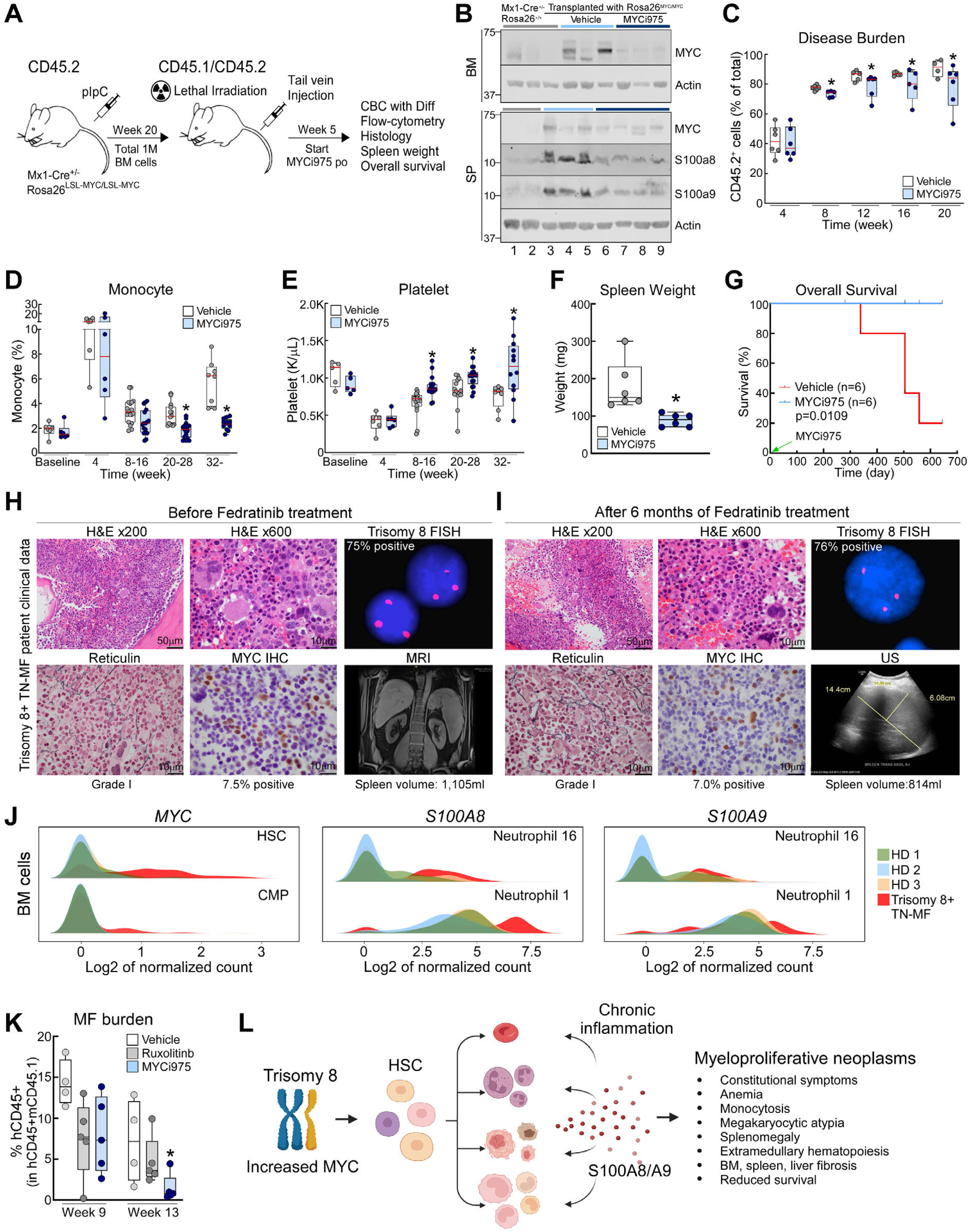
Inhibition of MYC effectively suppresses TN-MF disease progression. **A,** Schematic of in vivo mouse studies. **B,** MYC, S100a8, and S100a9 protein levels in the BM and spleen cells of vehicle- *vs.* MYCi975- treated mice (that were transplanted with BM cells from a Mx1-Cre^+/-^;Rosa26^LSL-MYC/LSL-MYC^ mouse) were compared to protein levels in Mx1-Cre^+/-^;Rosa26^+/+^ control mice. **C,** Disease burden (CD45.2^+^ cells as % of total CD45.1^+^+CD45.2^+^ cells) was serially assessed by flow cytometry at the indicated time points using PB samples. **D-E,** PB CBC analyses. Baseline CBC was performed 1 week prior to transplant. **F,** Spleen weight at endpoints. MYCi975-treated mice were sacrificed at the same time when vehicle-treated mice approach their endpoints for paired analyses. **G,** KM curves of OS. Demographic and laboratory profiles are described in Table S14. **H-I,** H&E, reticulin, and MYC IHC staining, as well as cytogenetics and spleen size in a trisomy 8+ TN-MF patient before and after 6 months treatment of fedratinib (400mg po daily). **J,** *MYC* and *S100A8/A9* levels in HSCs/CMPs and neutrophils, respectively, in trisomy 8+ TN-MF *vs.* HD. **K,** PB disease burden (expressed as % of hCD45^+^ cells in hCD45^+^+mCD45.1^+^ cells) in recipient mice treated with vehicle, ruxolitinib, or MYCi975. Mice were randomized at 9 weeks post-transplant, and disease burden was compared at 13 weeks post-transplant. **L,** Graphical summary of the MYC-S100A8/A9 circuit in TN-MF. Actin blots in (B) were from the same samples run on different gels. Box plots in (D-F, K) represent data from 6 and 4-5 mice in each group, respectively. *, *P*<0.05 compared with control group.

Based on these results, the effects of MYC inhibition were assessed in a patient-derived xenograft (PDX) from a trisomy 8+ TN-MF patient. First, BM samples were collected at the time of diagnosis from a trisomy 8+ TN-MF patient who had splenomegaly and profound constitutional symptoms (Figure 7H; Table S2). Staining of the BM biopsy revealed a hypercellular marrow with megakaryocytic atypia, grade 1 reticulin fibrosis, and elevated MYC protein levels (Figure 7H). scRNA-seq analysis of BM cells confirmed increased *MYC* and *S100A8/A9* in HSCs and neutrophils, respectively (Figure 7J). BM cells from this patient were then transplanted via intratibial injection into sublethally irradiated NSG-SGM3 mice. Following confirmation of engraftment of human CD45^+^ cells at 9 weeks post-transplant, mice were randomized to treatment with MYCi975, ruxolitinib, or vehicle (Figure 7K). MYCi975 treatment effectively suppressed disease progression as evidenced by a significantly lower percentage of hCD45^+^ cells in PB compared to mice treated with ruxolitinib or vehicle (Figure 7K). Importantly, treatment of the patient with fedratinib for 6 months showed only a modest improvement in spleen size without significant changes in BM cellularity, the percentage of trisomy 8+ and MYC+ cells, and reticulin fibrosis (Figures 7H-I, S7Q). These results indicate MYC as a clinically actionable target in MYC-driven TN-MF (Figure 7L). Further, our studies suggest that a small molecule targeting MYC can potentially irradicate malignant trisomy 8+ TN-MF clones, which is a major advance as malignant clones in MF are not eradicated by current JAK2 inhibitors^18,19^.

## DISCUSSION

While increased MYC expression in lymphoid malignancies is mainly driven by *MYC* gene rearrangements^23–25^ and *MYC* somatic mutations have been identified in a variety of cancers^4–9,39,70–76^, these changes are rare in myeloid neoplasms^39,71–76^. However, the *MYC* gene is located on chr 8q24 where copy number gain is frequently observed across all myeloid malignancies^35–39^, and we and others have shown that *MYC* copy number gain is associated with increased MYC levels and inferior survival outcomes in AML and MDS^26,27,34^. Although *MYC* copy number gain is also frequent in MF patients^36–38,77^, its oncogenic role in MF is largely unknown. Here we have shown that trisomy 8 commonly occurs in TN-MF, but not in other molecular subtypes, and is associated with adverse clinical outcomes. Further, our scRNA-seq and IHC analysis of BM cells revealed significantly higher levels of MYC mRNA and protein in HSCs from trisomy 8+ TN-MF patients compared to normal donors. These observations, along with subsequent findings discussed below, identify MYC as an oncogenic driver of TN-MF that is independent of JAK2 pathway mutations.

MYC transcriptional programs are activated as HSCs differentiate into myeloid progenitors^78^, and these programs control the balance of HSCs self-renewal and differentiation into mature myeloid cells^79^. In accord with these findings, there was a dramatic reduction in GM-CFU following *Myc* silencing in Rosa26-CreERT2^+/-^;*Myc*^fl/fl^ mice, and the opposite effect upon MYC overexpression in Mx1-Cre^+/-^;Rosa26^LSL-MYC/LSL-MYC^ mice. Although these effects of MYC up- or down-regulation on myeloid progenitors ex vivo have been observed across many pre-clinical studies, the in vivo effects of increased MYC expression in HSCs have been variable^26,53,79–81^. For example, studies using retroviral-mediated MYC overexpression in HSCs have shown that MYC can provoke AML in vivo, and recent studies using human CD34^+^ cord blood stem cells have shown that lentiviral-mediated MYC overexpression in HSCs drives AML only in the presence of continuing IL-3/GM-CSF co-stimulus^32,33^. In contrast, culture of HSCs ex vivo following MYC transduction yields highly homogeneous committed myeloid progenitors^82–86^. These varied findings might reflect the effects of different MYC expression levels. Previous in vivo studies using the VavP-*MYC* transgenic mouse model established that MYC levels govern hematopoietic tumor type, with high HSCs MYC levels favoring lymphoma development and lower MYC levels inducing the formation of marrows having megakaryocyte atypia and cytokine hypersensitivity, all of which are hallmarks of MPN, although the precise disease phenotypes and underlying mechanisms were not fully characterized^25,53^.

Using two independent transgenic mouse models that inducibly increase MYC expression to levels found in myeloid disease-prone VavP-*MYC* transgenic mice, we have shown that low levels of MYC overexpression in HSCs induce a myeloid malignancy that is most consistent with MF^56,57^. Further, using in vivo transplant studies, pharmacologic inhibition of MYC was shown to effectively reverse MYC-driven MF phenotypes and significantly improve OS. Importantly, the MYC inhibitor also suppressed disease progression in a PDX model using primary BM cells from a TN-MF patient with trisomy 8. Accordingly, this study is the first to show that MYC levels are increased in a subgroup of TN-MF patients, specifically in patients with trisomy 8 and that MYC provokes MF independent of JAK2 pathway mutations, and suggest that this MYC circuit is targetable using a small molecule inhibitor.

Chronic inflammation is a hallmark of MF pathogenesis. While pro-inflammatory cytokines induced by constitutive activation of JAK/STAT pathway play pivotal roles in MF development in patients with *JAK2/CALR/MPL* mutations, the oncogenic drivers that promote inflammation in TN-MF have heretofore been unclear. Our studies establish that increased MYC expression in HSCs induces S100A8/A9-mediated chronic inflammation, and that *S100a9* deficiency significantly impairs MYC-driven MF development, leading to improved OS. Importantly, MYC-driven MF cells are not sensitive to JAK2 inhibitors ex vivo, a finding supported by additional PDX experiments and clinical studies performed in parallel. Collectively, these findings suggest that MF patients whose disease depends on MYC activation need alternative therapeutic strategies using agents targeting the MYC-alarmin axis rather than JAK2 inhibitors.

One of the most striking findings is the complex nature of the inflammatory network driven by the MYC-alarmin axis. This inflammatory network requires an intact BM niche and likely involves various hematopoietic and non-hematopoietic cells. Even though S100a8/a9 levels are significantly increased by MYC in BM and/or spleen cells in vivo, MYC-directed upregulation of S100a8/a9 is completely abolished in BM cells cultured ex vivo. scRNA-seq analyses also suggest that this inflammatory network likely includes contributions of various hematopoietic and non-hematopoietic cells. Indeed, a wide range of hematopoietic cells that are minimally inflammatory under normal conditions (e.g., HSCs, myeloid progenitors, T-lymphocytes) can affect diverse cell types via S100a8/a9 when MYC is activated. This is especially manifest in myelomonocytic lineage cells (e.g., monoblasts, monocytes, macrophages) that are predominantly affected by S100a8/a9 signals through various receptors (e.g., ALCAM, CD68) to promote the selective expansion of highly inflammatory M1 macrophages^87,88^. Notably, MF patients have increased macrophages in their BMs compared to patients with other MPN subtypes or normal BM^89^, and depletion of macrophages effectively prevents MF development in a model of *JAK2^V617F^* driven MF^63^. Taken together, our data suggest that expansion of inflammatory monocytes and macrophages plays a key role in MF development regardless of JAK2 pathway mutations. If so, selective targeting of these sub-populations using small molecules or monoclonal antibodies could be an alternative therapeutic approach.

Our studies raise several important questions. First, MYC, as a direct downstream target of the JAK/STAT pathway, also plays key roles in MPN cell survival and resistance to JAK2 inhibitors^40–42,90,91^. Although *S100A9* levels were elevated in the CD34^+^ cells of *JAK2^V617F^* mutant MF patients^59,60^, it is unclear whether the MYC-S100A9 circuit also plays an active role in MPN driven by *JAK2* pathway mutations. The answer to this clinically important question will have consequences for eligibility criteria in future clinical trials using agents targeting the MYC-S100A9 pathway and for the design of optimal combination therapies. Second, S100A9 from mesenchymal stromal cells (MSCs) is known to play an essential role in the JAK2^V617F^- and TPO-driven MF pathogenesis^45^. Indeed, Gli1^+^ or LepR^+^ MSCs have been identified as fibrosis-driving myofibroblasts in the context of *JAK2* pathway mutations^92–94^, but it remains to be determined whether MYC in HSCs can also activate Gli1^+^ or LepR^+^ MSCs. Third, consistent with observations from other MPN mouse models^95–101^, we observed significant fibrosis in the spleen and liver, but less in the BM in MYC-driven murine MF. While it is possible that inherently higher levels of *S100A8/A9* in the normal mouse BM niche *vs.* human BM reflect a tolerability of higher S100A8/A9 levels in murine BM before fibrosis occurs, other differences in the BM niche between mouse and human, as well as between mouse BM and spleen, also need to be explored as potential explanations for the differences in fibrosis phenotypes. Fourth, *S100a9* deficiency only partially abrogated MYC-driven MF phenotypes, suggesting that additional targets downstream of MYC also contribute to MF pathogenesis. As S100A9-directed agents become available, it will be important to see whether inhibition of S100A9-mediated inflammation will translate into improved survival outcomes in patients with pre-existing MF and to address this question separately in patients with MYC-driven MF vs. *JAK2/CALR/MPL*-driven MF. Fifth, even though Tasquinimod acutely inhibits S100A9 signaling and has been well studied in both mice and humans^67,68^, this agent showed modest effects in suppressing MYC-driven MF phenotypes and, upon long-term treatment, was associated with effects such as anemia and neutrophilia that were not seen with *S100a9* knockout, indicating the need for developing S100A9 inhibitors with improved efficacy and safety profiles. Sixth, because TN-MF is associated with higher risk of transformation into AML *vs*. other molecular subtypes, identifying secondary genetic alterations and defining how they cooperate with MYC to promote AML transformation will be critical to improve clinical outcomes in these patients. Finally, trisomy 8 also occurs in HSCs in the normal population, with a prevalence that increases with aging^102–105^ in much the same fashion as somatic mutations (e.g., *JAK2, DNMT3A, TET2*) in HSCs that drive clonal hematopoiesis and provoke a pro-inflammatory state linked to other co-morbidities such as cardiovascular disease^106–109^. Thus, the systemic impact of inflammatory signals that can arise from MYC-dependent clonal hematopoiesis warrants future investigation.

In summary, the studies reported herein are the first to describe an oncogenic role of MYC in MF pathogenesis, reveal a previously undescribed circuit that connects MYC, alarmins, and inflammation, and validate the MYC-S100A9 axis as a novel therapeutic vulnerability in a subgroup of MF patients with very poor outcomes. Accordingly, our results provide a strong rationale for testing agents targeting MYC or S100A9 in early phase clinical trials in MF patients having increased MYC activity.

## METHODS

### Human subjects and clinical data

Using the Moffitt Cancer Center (MCC) Total Cancer Care (TCC) dataset, we retrospectively identified cases diagnosed with primary and secondary myelofibrosis (MF) from 2000 to 2021. The patients had provided written informed consent to be included in the dataset, and our study was approved by the MCC IRB (protocols MCC#14690 and MCC#18864). MYC and S100A9 protein expression were assessed by immunohistochemistry (IHC) staining as described below. Somatic mutations were assessed using targeted exome sequencing (developed at MCC) examining 54 or 98 myeloid genes as previously described^110,111^. Clinical variables and disease-related prognostic factors, including age, gender, cytogenetics, and treatment regimens, were characterized at the time of MF diagnosis, and were annotated using descriptive statistics. The LFS and OS were estimated with the Kaplan-Meier method and compared using log-rank test. All statistical analyses were performed using SPSS v24.0 and GraphPad Prism 7.

### Animal studies

All animal studies were performed in compliance with the National Institutes of Health Guidelines under a protocol approved by the H. Lee Moffitt Cancer Center & Research Institute and the University of South Florida Institutional Animal Care and Use Committee (IACUC). Mouse genotypes from tail biopsies were determined using real time PCR with specific probes designed for each gene (Transnetyx). All mice used in our in vivo studies were C57BL/6J background except PDX studies that used NSG-SGM3 mice (see Table S15).

Mx1-Cre^+/-^;Rosa26^LSL-MYC/LSL-MYC^ and Scl-CreERT^+/-^;Rosa26^LSL-MYC/LSL-MYC^ mice were generated by crossing Mx1-Cre^+/-^ or Scl-CreERT^+/-^ mice with Rosa26^LSL-MYC/LSL-MYC^ mice, respectively. Mx1- Cre^+/-^;Rosa26^LSL-MYC/LSL-MYC^;S100a9^-/-^ mice were generated by crossing Mx1-Cre^+/-^;Rosa26^LSL-MYC/LSL-MYC^ mice with S100a9^-/-^ mice. Mice at age 6∼11 weeks in both experimental and control groups were treated with pIpC (250µg/kg, d1 and d3, prepared in normal saline, i.p.) or tamoxifen (250mg/kg, d1-5, prepared in corn oil, po) to induce MYC gene expression. Using peripheral blood (PB) samples, complete blood counts (CBC) with differential were assessed 1 week prior to pIpC or tamoxifen treatment (baseline), and then every 4 weeks following drug treatment. Mice at pre-defined endpoints were sacrificed and primary BM, spleen, and liver tissues/cells were analyzed via the assays described below. Pre-defined endpoints used in our in vivo studies include (i) substantial weight loss (≥20% loss), (ii) abnormalities with movement or breathing, (iii) excessive lethargy, tremors, restlessness, failure to groom causing unkempt appearance, teeth grinding, or guarding (protecting the painful area), (iv) failure to show normal patterns of inquisitiveness, and (v) inability to urinate or defecate.

For competitive congenic transplant experiments, mice were placed on Baytril water (0.25 mg/ml) 72 hr prior to irradiation to prevent opportunistic bacterial infections. Lethally irradiated (1100cGy) CD45.1^+^/CD45.2^+^ WT recipient mice were transplanted with a total of 1 million CD45.2^+^ Mx1-Cre^+/-^;Rosa26^LSL-MYC/LSL-MYC^ or Mx1-Cre^+/-^;Rosa26^+/+^ (that were treated with pIpC 20 weeks prior to transplant) and CD45.1^+^ WT helper cells (mixed at 1:1 ratio) via tail vein injection 24 hr after irradiation. To confirm successful engraftment, PB samples were collected at 4 weeks following transplant, then incubated in ACK buffer twice for 5 min to lyse RBC. Disease burden was measured every 4 weeks by performing serial CBC with differential and flow cytometry as described below. Mice were harvested at pre-defined endpoints and primary BM and spleen cells were harvested for RNA extraction and qRT-PCR, immunoblotting, and ELISA assays. Femur, tibia, spleen, and liver tissues were also harvested. Tissues were incubated in Neutral Buffered Formalin for 24 hr, then stored in 70% EtOH until used. H&E, trichrome, and reticulin staining were performed as described^112^ or as noted below. Spleen and body weight were measured per protocol.

### Tissue culture

Cell lines were propagated at densities of <1 x 10^6^ cells/ml in RPMI 1640 medium containing 10% heat-inactivated fetal bovine serum (FBS), 100 units/ml penicillin G, and 2 mM glutamine (medium A) except for SET-2 cells, which were cultured in medium with 15% FBS. Primary BM cells were harvested from Mx1-Cre^+/-^;Rosa26^+/+^, Mx1-Cre^+/-^;Rosa26^LSL-MYC/+^, Mx1-Cre^+/-^;Rosa26^LSL-MYC/LSL-MYC^, Rosa26-CreERT2^+/-^;*Myc*^+/+^, and Rosa26-CreERT2^+/-^;*Myc*^fl/fl^ mice. BM cells were homogenized in PBS with 2% FBS, filtered through a 100-µm strainer and cultured in RPMI 1640 medium containing FBS (15%), glutamine (2 mM), mouse IL-3 (10 ng/ml), mouse IL-6 (10 ng/ml), mouse SCF (10 ng/ml), and penicillin G (100 units/ml).

### Colony forming assays

Primary mouse BM and spleen cells were harvested under sterile conditions. Cells were centrifuged for 5 min at 1,000 rpm and pellets were resuspended in RBC lysis buffer. Samples were then centrifuged for 5 min at 1,000 rpm and resuspended in 2% FBS IMDM media to 2 x 10^4^ cells/100 µl stock for BM cells and 2 x 10^5^ cells/100 µl for spleen cells. After mixing 400 µl of each stock with 4 ml of Methocult^TM^ GF M3434, 1.1ml of the mixture (2 x 10^4^ BM cells and 2 x 10^5^ spleen cells) was plated in triplicate into 6-well dishes (SmartDish^TM^, STEMCELL Technology). After incubation for 10 days, colonies were manually counted.

### Immunoblotting

Following treatment with the indicated concentrations of drugs or isolation of the primary cells from BM and spleen, cells (5x10^6^/aliquot) were washed with cold Dulbecco’s phosphate buffered saline (PBS), lysed with RIPA buffer (10 mM Tris [pH 7.4], 100 mM NaCl, 1 mM EDTA, 1 mM EGTA, 1% Triton X-100, 10% glycerol 0.1%SDS, 0.5% deoxycholate) or ARF buffer (50 mM HEPES [pH 7.5], 150 mM NaCl, 1 mM EDTA, 2.5 mM EGTA, 0.1% Tween-20) that contained a complete protease inhibitor mini-tablet (1 tablet/10 ml), PMSF (1 mM), beta-glycerophosphate (10 mM), sodium fluoride (1 mM), and sodium orthovanadate (1 mM). After lysates were sonicated, and centrifuged at 15,000 rpm for 30 sec or 2 min, supernatants were carefully collected. Protein concentration was determined using a BCA Assay. Protein was separated on SDS-PAGE, transferred to nitrocellulose membranes, and blotted with specific primary antibodies as listed in the Table S15. Images were captured using Odyssey Fc Imaging System (LI-COR).

### ASC crosslinking assays

Primary BM and spleen cells (1 x 10^7^ cells per mouse) were lysed with RIPA buffer following RBC lysis as described above. Cell lysates were sheared 30 times through a 21-guage needle in microcentrifuge tubes, then centrifugated for 8 min at 1,800 rpm. Supernatants were incubated on ice for 10 min, then split into two tubes; one for immunoblotting as described above and the other for ASC speck cross-linking. For cross-linking, lysates were centrifugated for 10 min at 5,000 rpm and supernatants were removed. After pellets were washed twice with PBS, freshly made disuccimidyl suberate solution (dissolved in anhydrous DMSO, final concentration 2 mM) was added into each pellet followed by incubation at room temperature for 30 min on a rotator. Samples were centrifugated for 10 min at 5,000 rpm, supernatants were discarded, pellets were resuspended with 30 µl of 1x Laemmli buffer (2% SDS, 5% 2-mercaptoethanol, 10% glycerol, 0.002% bromophenol blue, 0.0625M Tris-HCl, pH 6.8) and subjected to SDS-PAGE as described above. Loading volumes were adjusted based on protein concentration of whole cell lysates.

### Enzyme-linked immunosorbent assay

Serum was collected from mice at the specified times in each protocol. Following incubation of whole blood at room temperature for 30-40 min, clots were removed by centrifugating blood samples at 2,000 x *g* for 10 min in a refrigerated centrifuge. This step was repeated one more time to remove residual blood clots, then serum samples were stored at -80°C until used. ELISA was performed following the manufacturer’s protocol (R&D System). Briefly, a 96-well microplate was coated with capture antibody (100 µL per well) overnight at room temperature. After plates were blocked with blocking buffer (300 µl per well) for 1 hr at room temperature, 100 µl of serum samples (diluted at 1:10 ratio with reagent diluent) or standard were added to the plate, which was then incubated for 2 hr at room temperature. After washing and addition of 100 µl of detection antibody to each well, plates were incubated for 2 hr at room temperature. Streptavidin-HRP, substrate solution and stop solution were added per protocol. Optical density was immediately measured using a microplate reader. Values measured at 450 nm were subtracted from values measured at 540 nm.

### RNA preparation and qRT-PCR assays

RNA from primary BM or spleen cells, tumor tissues of Eµ-*Myc* mice, or human MPN cells was isolated using NucleoSpin RNA II kit. cDNA was prepared using iScript^TM^ cDNA synthesis kit and qRT-PCR was performed using a CFX96 Touch Real-Time PCR Detection System (BioRad). Data were analyzed using the following equations: ΔCt = Ct(sample) - Ct(endogenous control); ΔΔCt = ΔCt(sample) - ΔCt(untreated or control); Fold Change = 2^-ΔΔCt^. *Ubiquitin* and *Actin* served as the endogenous controls. Primers are listed in the Table S16.

### Lactate dehydrogenase activity

Serum lactate dehydrogenase (LDH) levels were quantified according to a protocol provided with the CyQUANT LDH Cytotoxicity Assay Kit. A 50 µl serum sample was aliquoted into 96-well microplate, then mixed with 50 µl of reaction mixture provided in the kit. Following incubation for 30 min at room temperature in the dark, 50 µl of stop solution was added into each well and mixed by gentle tapping. Absorbance was measured at 490 nm and 680 nm. LDH activity was calculated by subtracting the 680 nm absorbance value (background) from the 490 nm absorbance.

### Flow cytometry analyses

Primary BM and spleen cells were harvested as above. Following RBC lysis using ACK buffer, cells were resuspended in PBS with 2% FBS. To characterize changes in individual hematopoietic lineages, cells were stained with Zombie near IR (ZNIR, viability dye), mouse anti-CD16/32 (Fc block), -Ter119-V450, -B220-PE, -Ly-6G/Ly-6C-APC, -CD11b-BUV737, -CD3-BV786, and -F4/80-BUV395 fluorochrome-conjugated antibodies. To characterize the hematopoietic stem cells and progenitor populations, cells were stained with mouse Lin cocktail (BV421), anti-CD34-PE, - CD117-APC, -Ly-6A/Ly-6E-BB515, -CD150-PE-Dazzle 594, -CD48-Brilliant Violet 711, and - CD16/CD32-BUV395 conjugated antibodies. Cells were fixed with 4% paraformaldehyde for 10 min, washed with FACS buffer three times, and then subjected to flow cytometry.

In competitive transplant and xenograft experiments, cells were incubated with ZNIR and washed with PBS. After incubation with mouse FC blocking antibody for 10 min, cells were stained with anti-mouse CD45.1 and CD45.2 antibodies, fixed with 4% paraformaldehyde for 10 min, washed with FACS buffer three times, and subjected to flow cytometry as described^113^. To assess apoptosis, cells were treated with drugs as indicated, stained with Annexin-V and propidium iodide (PI), and then the percentage of AnnexinV/PI positive populations were quantified by flow cytometry as described^114^.

### Immunohistochemistry

Paraffin-embedded BM trephine biopsies were used for immunohistochemistry (IHC) analyses as described^34^. Blocks were sectioned to 4-µm in thickness. Unstained slides were deparaffinized using automated system with EZ Prep solution (Ventana Medical System) and stained with MYC or S100A9 antibody using a Ventana Discovery XT automated system (Ventana) per the manufacturer’s protocols. Slides were reviewed by an independent hematopathologist. Protein expression levels were calculated as % (0∼100%) of MYC or S100A9 positive cells multiplied by staining intensity (1^+^∼3^+^) as described^28^.

### Reticulin and trichrome staining

The reticulin and trichrome stains were based on the method of Gomori and Snook (Artisan Reticulin-Nuclear Fast Red Stain Kit) and the original Masson’s procedure (Artisan Masson’s Trichrome Stain Kit). Staining was performed by using Artisan Staining System following protocols provided by manufacturer.

### Single cell library production and sequencing

Primary mouse BM cells, healthy donor BM MNCs, and MF patient BM MNCs were prepared following the protocol provided by 10x Genomics (CG000392, Rev A). For single cell encapsulation and library production, we used the Chromium Single Cell 3’ Reagent Kits (v2) per user guideline (CG00052, Rev D). Sequencing was performed on a Nextseq 2000 platform (Illumina) aiming for a minimum of 50,000 reads/cell.

### Single-cell RNA-seq data processing, filtering, batch effect correction, and clustering

A customized reference genome was built by adding MYC sequences to the GRCm38 mouse transcriptome using the *mkref* module of Cell Ranger (v6.0, 10X Genomics). Raw sequencing reads from single cells were aligned against the customized mouse reference and processed using *count* module of Cell Ranger. Gene-barcode matrices containing only barcodes with UMI counts passing threshold were imported to Seurat^115^ for further analysis. Genes detected in less than 3 cells were excluded; cells with less than 200 genes detected or greater than 10% mitochondrial UMIs were also filtered out. Doublets were detected using Scrublet^116^, DoubletFinder^117^, scDblFinder^118^, and doubletCells implemented in scran^119^, assuming 0.08% doublet rate for every 1,000 sequenced cells. Cells identified as doublets by at least two algorithms were removed from further analysis. Raw UMI counts were log normalized and the top 5,000 variable genes were identified using “vst” method implemented in the *FindVaribleFeatures* function in Seurat. T cell receptor and immunoglobulin genes were removed from the variable genes to prevent clustering based on V(D)J transcripts. S and G2/M cell cycle phase scores were assigned to cells using *CellCycleScoring* function.

To further remove batch effects, individual samples were then integrated using *FindIntegrationAnchors* and *IntegrateData* functions^120^ with 8,000 anchor genes and 40 dimensions of canonical correlation analysis (CCA). Briefly, dimension reduction was performed on each sample using diagonalized CCA, and L2-normalization was applied to the canonical correlation vectors to project all samples into a shared space. The mutual nearest neighbors (MNS) across cells from different datasets were used as “anchors” to encode the cellular relationship between samples. Samples were integrated based on correction vectors for sample calculated form anchors.

Integrated data were further scaled using *ScaleData* function by regressing against total read count, percentages of mitochondrial UMIs, and cell cycle phase scores (S and G2/M). A shared nearest neighbor (SNN) based graph was constructed using the top 40 principal components, and clusters were identified using the Louvain algorithm using *FindCluster* function at resolution = 1 (33 clusters). Uniform Manifold Approximation and Projection (UMAP) were generated by *RunUMAP* function and used for visualization.

### Mouse bone marrow cluster annotation

Differential expression analysis comparing each cluster *vs.* all others was performed using *FindAllMarkers* function in Seurat with default settings. Genes with log2(fold-change) >0.25 and Bonferroni-corrected p-value <0.05 were considered differentially expressed. Clusters were annotated by comparing the cluster-specific genes to canonical markers for Hematopoietic Stem cells (HSCs) (*Myct1, Angpt1, Rgs1, Pde4b, Ncl*), Erythroblasts (*Car2, Gypa, Prdx2, Alas2, Slc4a*1), Monoblasts (*F13a1, Tmsb10, Ly86, Lgals1, Ccr2*), Monocytes (*Pld4, S100a4, Itgb7, Ahnak, Pid*1), Myeloblasts (*Elane, Mpo, Ms4a3, Ctsg, Prtn3*), Myelocytes (*Anxa1, Ltf, Lcn2, Fcnb, Camp*), Megakaryocytes (*Timp3, Rab27b, Pls1, P2rx1*), Basophils (*Ccl3, Fcer1a, Ms4a2, Hdc, Cyp4f18*), Macrophages (*Cd300e, Batf3, C1qb, Cd68*), M1 macrophages (*Cxcl9, Cxcl10, Cxcl11, Il1a, Tnf, Il6, Ccl5, Ccl2, Ccl4, Cxcl1, Ccl7, Il27, Il10, Cxcl3, Tnfsf15, Ccl3, Il11*), M2 macrophages (*Ccl24, Ccl8, Ccl12, Ccl9, Cxcl12, Ccl6*), Neutrophils (*S100a9, S1000a8, Mmp9, Il1rn, Cxcr2*), Granulocyte-monocyte progenitors (GMPs) (*Mpo, Cstg, Prtn3, H2afy*), Megakaryocyte-erythroid progenitors (MEPs) (*Pf4, Gata2, Cdk6, Gas5*), Common Myeloid progenitors (CMPs) (*Cd34, Npm1, Eef1g, H2afy*), T cells (*Cd3e, Cd3d, Cd4, Cd8a*), NK cells (*Klrd1, Ncr1, Klra4*), Pro-B cells (*Vpreb3, Akap12, Cd79b, Chchd10, Cecr2*), Pre-B cells (*H2-Ab1, Shisa5, March1, Cd74, H2-DMb2*), Plasma cells (*Igkc, Ighm, Jchain*), Endothelial cells (*Kdr, Lrg1, Cldn5, Fam167b, Osmr*), pDC (*Siglech, Ccr9, Bst2, Pacsin1, Tcf4*), and Adipose cells (*Cxcl12, Igfpb5, Bgn, Igfbp4*). To confirm cell annotation, enrichment scores were calculated using *AUCell* algorithm implemented in SCENIC^121^ for signature genes reported in previous single-cell RNA-seq studies on BM, AML, and hematopoietic cells^122–125^. Human signatures were converted to their mouse homologs using R package biomaRt before sending for enrichment analysis. Expression of marker genes was visualized on UMAP or by violin plot using log normalized UMI counts. Heatmaps were used to visualize the z-score normalized average expression across clusters. The 33 clusters were further grouped into 23 major cell types based on their annotation. Differential gene expression analysis was performed to generate cell type specific marker genes.

### Human bone marrow cluster annotation

Human bone marrow (BM) scRNA-seq data were analyzed as described above. A shared nearest neighbor (SNN) based graph was constructed using the top 40 principal components, and clusters were identified using the Louvain algorithm using *FindCluster* function at resolution = 1 (34 clusters). Clusters were annotated by comparing the cluster-specific genes to canonical markers for Hematopoietic Stem cells (HSCs) (*SPINK2, CRHBP, MEIS1, MLLT3*), Erythroblasts (*BROX2, AHSP, HBB, ALAS2*), Basophils (*GATA2, LTC4S, CLC, IL3RA*), Macrophages (*CD68, CD163, CD300E*), Neutrophils (*CXCL8, S1000A8, S1000A9, CSF3R, CXCR2*), Megakaryocytes (*PPBP, PF4, GP9, SDPR*), Common myeloid progenitors (CMPs) (*MPO, FLT3, CEBPA, ELANE*), multi-lymphoid progenitors (*DNTT*, *CXCR4*, *BLNK*, *IGLL1*, *EBF1*), T cells (*CD3E, CD3D, CD4, CD8A*), Naïve T cells (*TCF7*, *IL7R*, *TCF7*, *CCR7*), Naïve-memory T cells (*CCL5*, *IFITM1*, *LTB*, KLF2), Effector-memory T cells (*GZMK*, *EOMES*, *KLRG1*, *CD27*), Effector T cells (*CCL3*, CCL4, *GZMB*, *GZMA*, PRF1), CD16^+^ NK cells (*FCGR3A, KLRB1, KLRD1*), CD56^+^ NK cells (*NCAM1*, *KLRB1*, *KLRD1)*, Pro-B cells (*CD79A*, *SOX4, IL7R, RAG2, LEF1*), Mature-B cells (*CD79A*, *MS4A1, TNFRSF13C, ITGAM*), Plasma cells (*IGHA1, IGHG1, IGHG2, IGHG4*), pDC (*LILRA4, IRF7, IL3RA, TCF4, PACSIN1*), and cDC2 (*CD1C, FCER1A, CLEC10A, HLA-DQA1*). Cell annotation was further confirmed by AUCell scores of signature genes reported in previous scRNA-seq studies on human BM cells^122–126^.

### InferCNV analysis

Copy number variation patterns in human BM single cells were extracted using InferCNV v3.1.5. (https://github.com/broadinstitute/inferCNV, Trinity CTAT Project). 3000 cells from the three HD samples were randomly selected and served as reference normal cells for de-noise control. Cells from the TN-MF patient were selected as observations. InferCNV analysis was performed using “denoise” mode to correct for batch effects from different patients. Clusters of cells with copy number gain observed on chr8 were determined as trisomy 8+ cells in the TN-MF patient. The trisomy status was further confirmed by expression of chr8 genes. Briefly, 571 genes located on chromosome 8 were identified from Gencode annotation. Overall expression of these genes was calculated by *AddModuleScore* function in Seurat and visualized by histogram.

### Differential gene expression comparing MYC *vs.* WT mouse bone marrow cells

Composition (%) of the 23 cell types was calculated and compared between MYC *vs.* WT samples and was visualized using ggplot2 stacked bar plot. The composition difference of each cell type between MYC *vs.* WT was calculated as log2(% in MYC/% in WT) and visualized as a bar plot. Differential expression analysis was performed to compare MYC *vs.* WT cells in each cell type using *FindMarkers* function of Seurat with default settings. Genes with log2(fold-change) >0.25 and Bonferroni-corrected p-value <0.05 were considered differentially expressed. The MYC differential genes were compared among clusters within HSCs and Progenitors, Myeloblasts and Myelocytes, Monoblasts and Monocytes, Neutrophils, T and NK cells, and B and Plasma cells, and visualized by Venn Diagram.

### Semantic analyses

First, differential gene expression analysis was performed with Seurat (v 4.0.2)^120^ between MYC and NULL phenotypes for all 23 major types. Genes with adjusted p value < 0.05 were considered for GO.bp enrichment analysis using clusterProfiler (v4.4.4)^127^. The obtained GO.bp terms from up-reg (fold change > 1) and down-reg genes (fold change < 1) were summarized via the REVIGO web application (http://revigo.irb.hr/)^128^ using p value of GO term as input metric allowing a similarity of 0.7.

### Cell-cell interaction (ligand-receptor) analyses

To identify and visualize cell-cell (ligand-receptor) interactions in MYC and WT cells, we loaded the scRNA-seq normalized gene expression data with cell type information into the R package CellChat ^129^. We also built mouse reference for cell-cell interactions with the data from OmniPath database (https://omnipathdb.org/)^130^ using R packages liana^131^ and OmniPathR^132^. A total of 5,964 mouse ligand-receptor interactions were used as a priori network information. The standard CellChat pre-processing steps were applied to identify overexpressed ligand-receptor interactions (identifyOverExpressedInteractions), compute the communication probability/strength between any interacting cell groups (computeCommunProb) and the communication probability/strength on the signaling pathway level, by summarizing all of the related ligands/receptors (computeCommunProbPathway); this resulted in 800 and 814 significant interactions for the MYC and WT groups, respectively. The selected interactions are visualized in a circular plot and the thicker edge line indicates a stronger signal based on the number of interactions. To visualize the difference in intensity of cell-cell interactions between the MYC and WT groups, we calculated both ligand and receptor expression means for the selected interaction in each pair of cell type, subtracted WT from MYC, and then visualized the difference of interaction intensity using the significant interactions (p<0.05).

### Pathway activity inference

A panel of 16 pathways was constructed with the 14 signaling pathways derived by PROGENy^133^ (Androgen, Estrogen, EGFR, Hypoxia, JAK-STAT, MAPK, NF-κB, PI3K, p53, TGF-β, TNF-α, Trail, VEGF, and WNT), known MYC target genes^134^, and the Alarmin pathway^135,136^. From the differential expression analysis comparing MYC *vs*. WT in each cell type, genes were ranked based on -log10(p-value)*(sign of log2(fold-change)), with the most up-regulated genes ranked at the top and the most down-regulated genes ranked at the bottom of the list. Pre-ranked Gene Set Enrichment Analysis (GSEA) was performed on these ranked gene lists against the 16 pathways using R package fgsea^137^ with 10,000 permutations. The normalized enrichment scores (NES) were used to denote the activity of each pathway and visualized by hierarchically clustered heatmap using R package ComplexHeatmap^138^, where positive NES presented up-regulated activity while negative NES presented down-regulated activity in MYC compared to WT.

### Synthesis of Tasquinimod

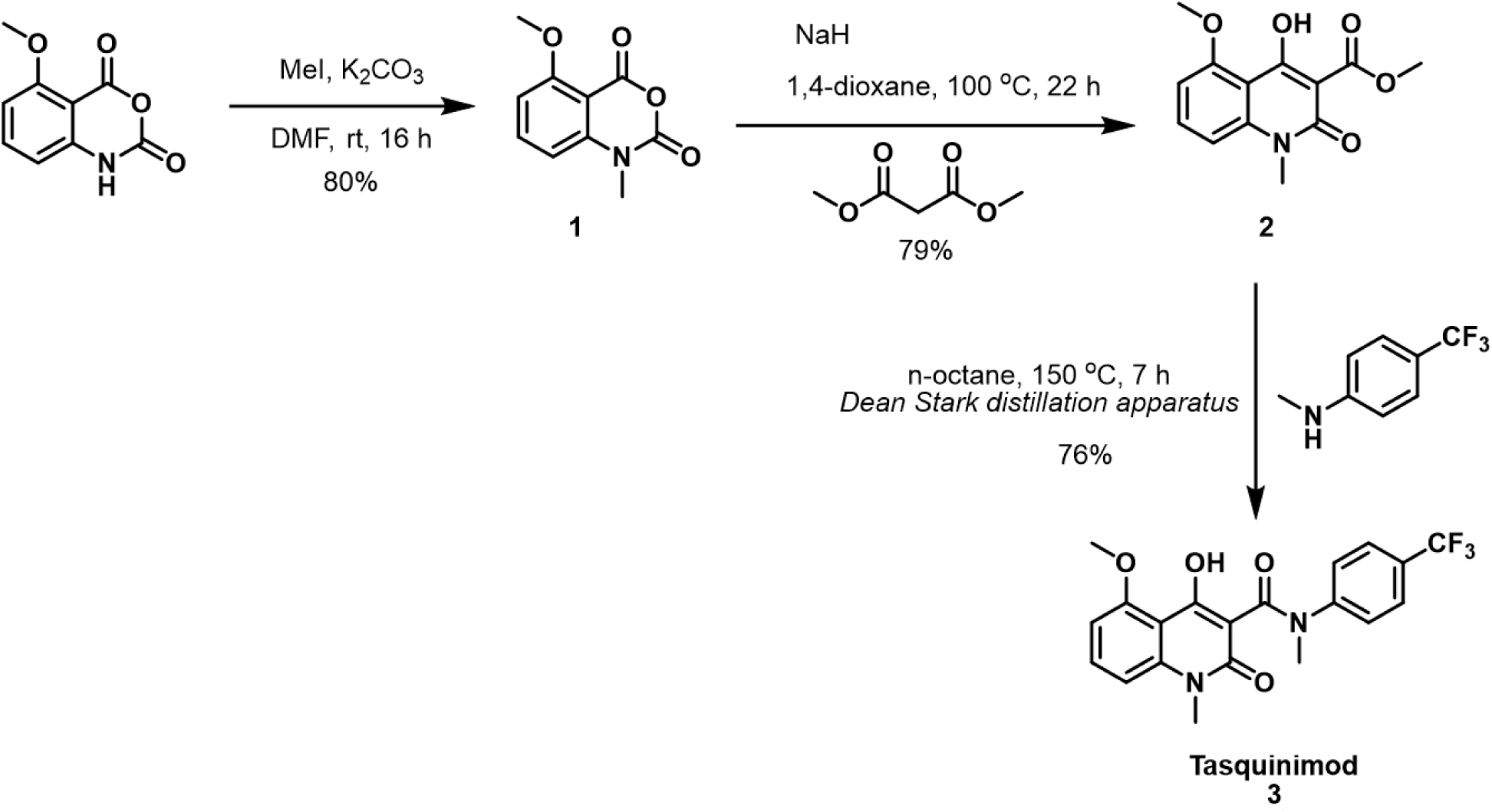

Synthesis of 5-Methoxy-1-methyl-2H-benzo[d][1,3]oxazine-2,4(1H)-dione (compound **1**): A dry 100 mL round-bottom flask with a stirring bar under inert (argon) atmosphere, was charged with 5-methoxy-2H-benzo[d][1,3]oxazine-2,4(1H)-dione (3.00 g, 15.53 mmol, 1 equiv) and DMF (32 mL). K_2_CO_3_ (2.73 g, 19.7 mmol, 1.27 equiv) and MeI (1.04 mL, 16.8 mmol, 1.08 equiv) were added to the resulting solution and the mixture was stirred at room temperature for 16 hr. After this period, HCl (1 M, 66 mL) was added slowly and carefully. After complete addition of HCL, the mixture was stirred for 30 min and the solid obtained was filtered. The solid was washed with water (3 x 20 mL) and dried under vacuum to afford intermediate dione 1 (2.57 g, 12.4 mmol, 80% yield) as a white solid. ^1^H NMR (500 MHz, DMSO) δ 7.76 (t, *J* = 8.5 Hz, 1H), 6.95 (t, *J* = 8.6 Hz, 2H), 3.91 (s, 3H), 3.42 (s, 3H).^13^C NMR (126 MHz, DMSO) δ 161.4, 154.7, 148.1, 144.0, 137.8, 106.6, 106.4, 100.5, 56.4, 32.2. HPLC–MS (ESI+): *m/z* 208.1 (M+H)^+^. *m/z* calculated for C_10_H_10_NO_4_^+^ (M+H)^+^ 208.06.

Methyl-4-hydroxy-5-methoxy-1-methyl-2-oxo-1,2-dihydroquinoline-3-carboxylate (compound **2**): A dry 100 mL round-bottomed flask, equipped with a stirring bar and a condenser, under inert (argon) atmosphere, was charged with 1 (2.00 g, 9.65 mmol, 1 equiv) and 1,4-dioxane (13 mL), followed by NaH (60% dispension in mineral oil, 425 mg, 10.6 mmol, 1.1 equiv). Dimethyl malonate (1.21 mL, 10.6 mmol, 1.1 equiv) was added dropwise and the resulting mixture was stirred at 100 °C for 22 hr. After this period, the reaction mixture was cooled to room temperature and quenched with water (61 mL). The aqueous solution was acidified with 1 M aq HCl to pH 3 and left in the refrigerator for 4-16 hr to precipitate the product. The solid obtained was collected by filtration, washed with water (3x 20 mL) and dried under vacuum to afford intermediate 2 (2.02 g, 7.67 mmol, 79% yield) as a pale yellow solid. ^1^H NMR (500 MHz, CDCl_3_) δ 7.57 (t, *J* = 8.5 Hz, 1H), 6.95 (d, *J* = 8.7 Hz, 1H), 6.74 (d, *J* = 7.6 Hz, 1H), 4.01 (s, 4H), 4.00 (s, 3H), 3.64 (s, 3H). ^13^C NMR (126 MHz, CDCl_3_) δ 171.8, 170.9, 159.9, 159.8, 143.2, 134.4, 107.4, 105.2, 104.3, 99.1, 56.6, 52.9, 30.1. HPLC–MS (ESI+): *m/z* 264.1 (M+H)^+^. *m/z* calculated for C_13_H_14_NO_5_^+^ (M+H)^+^ 264.09.

4-Hydroxy-5-methoxy-N,1-dimethyl-2-oxo-N-(4-(trifluoromethyl)phenyl)-1,2-dihydroquino-line-3-carboxamide (Tasquinimod, Compound **3**): A dry 100 mL round bottom flask equipped with a stirring bar, a Dean Stark distillation apparatus, and a condenser, under inert (argon) atmosphere, was charged with intermediate 2 (1.50 g, 5.70 mmol, 1 equiv), *N*-methyl-4- (trifluoromethyl) aniline (1.58 mL, 11.4 mmol, 2 equiv), and *n*-octane (40 mL). The reaction mixture was heated to 150°C (oil bath temperature) to reflux, and the mixture was stirred at this temperature for 7 hr, approximately 20 mL of volatiles were collected in the Dean Stark apparatus. The reaction mixture was cooled down to room temperature and the solvent was removed under vacuum. The crude product was purified by column chromatography (SiO_2_, 50-100% EtOAc in hexanes) to afford the final compound **3** (Tasquinimod) (1.77 g, 4.36 mmol, 76% yield) as an off-white solid.

### Characterization of Tasquinimod

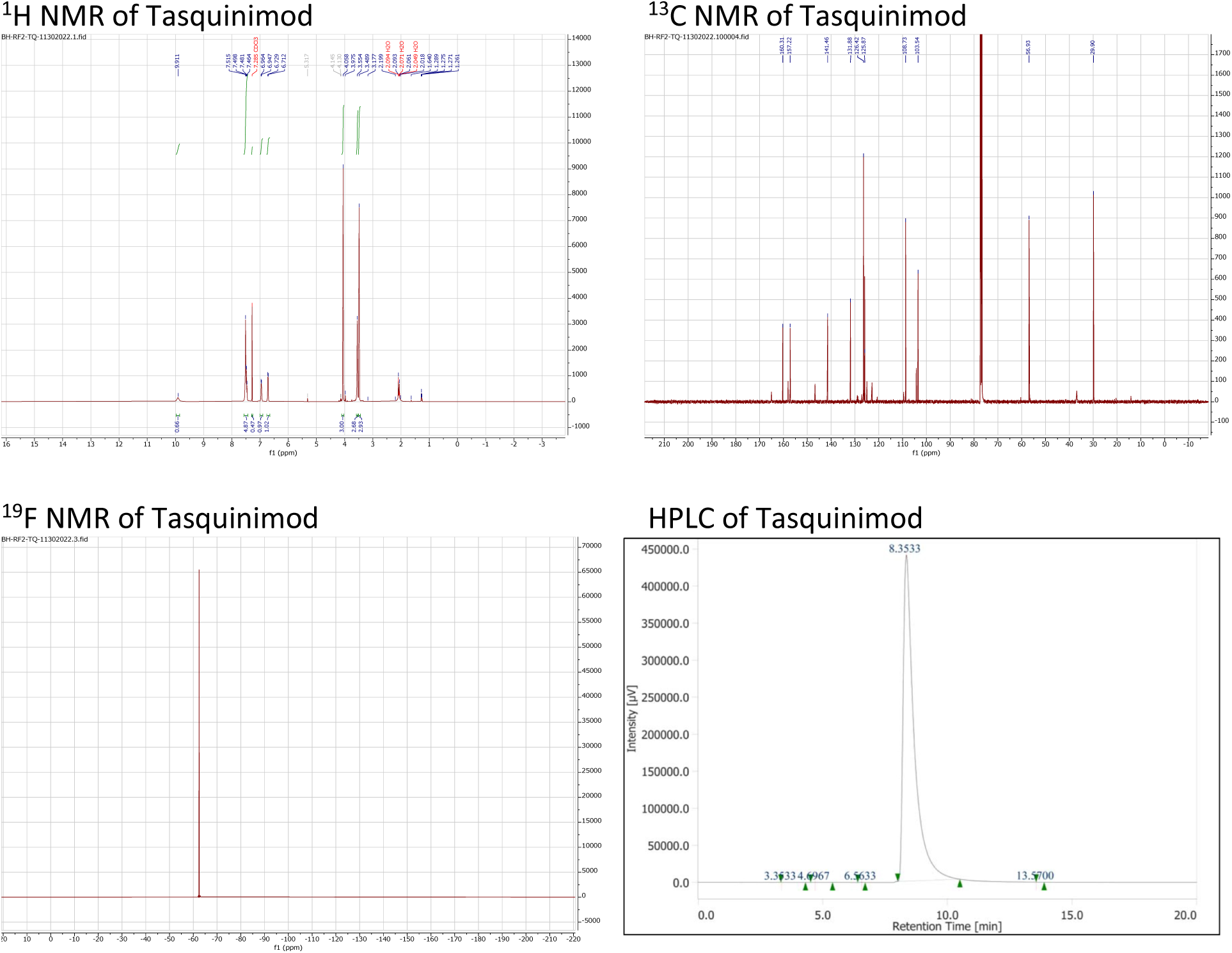

HPLC 99% (*t_R_* = 8.35 min, 5-95% CH_3_OH in water (0.1% TFA), 1mL/min, 20 min); 99% (*t_R_* = 4.92 min, 40% CH_3_OH in water (0.1% TFA), 1mL/min, 20 min). ^1^H NMR (500 MHz, CDCl_3_) δ 9.91 (s, 1H), 7.53 – 7.46 (m, 5H), 6.94 (d, *J* = 8.5 Hz, 1H), 6.70 (d, *J* = 8.2 Hz, 1H), 4.04 (s, 3H), 3.55 (s, 3H), 3.49 (s, 3H). ^19^F NMR (471 MHz, CDCl_3_) δ -62.42. ^13^C NMR (126 MHz, CDCl_3_) δ 165.1, 160.3, 158.1, 157.2, 146.8, 141.5, 131.9, 128.9 (q, ^2^*J*_CF_ = 31.5Hz), 126.4, 125.9 (q, ^3^*J*_CF_ = 3.8 Hz), 123.9 (q, ^1^*J*_CF_ = 272.1 Hz), 109.6, 108.7, 104.3, 103.5, 56.9, 36.9, 29.9. HPLC–MS (ESI+): *m/z* 407.1 (M+H)^+^. TOF-LCMS *m/z* calculated for C_20_H_18_F_3_N_2_O_4_^+^ (M+H)^+^ 407.1214, found 407.1222.

### Treatment of mice with Tasquinimod

Mx1-Cre^+/-^;Rosa26^LSL-MYC/LSL-MYC^ mice were treated with pIpC, and mice were then randomized to Tasquinimod *vs.* vehicle treatment. Tasquinimod treatment was initiated 5 days after the first dose of pIpC and Tasquinimod (30mg/kg/day) was administered via drinking water. Tasquinimod was dissolved in DMSO (final concentration 5%), then mixed with sucrose (final concentration 3%) and PEG300 (final concentration 2%) containing water. Vehicle group mice were treated with water containing DMSO (5%), sucrose (3%), and PEG300 (2%). Drinking water bottles were replaced every 3 days. Treatment was continued until endpoints.

### Synthesis of MYCi975

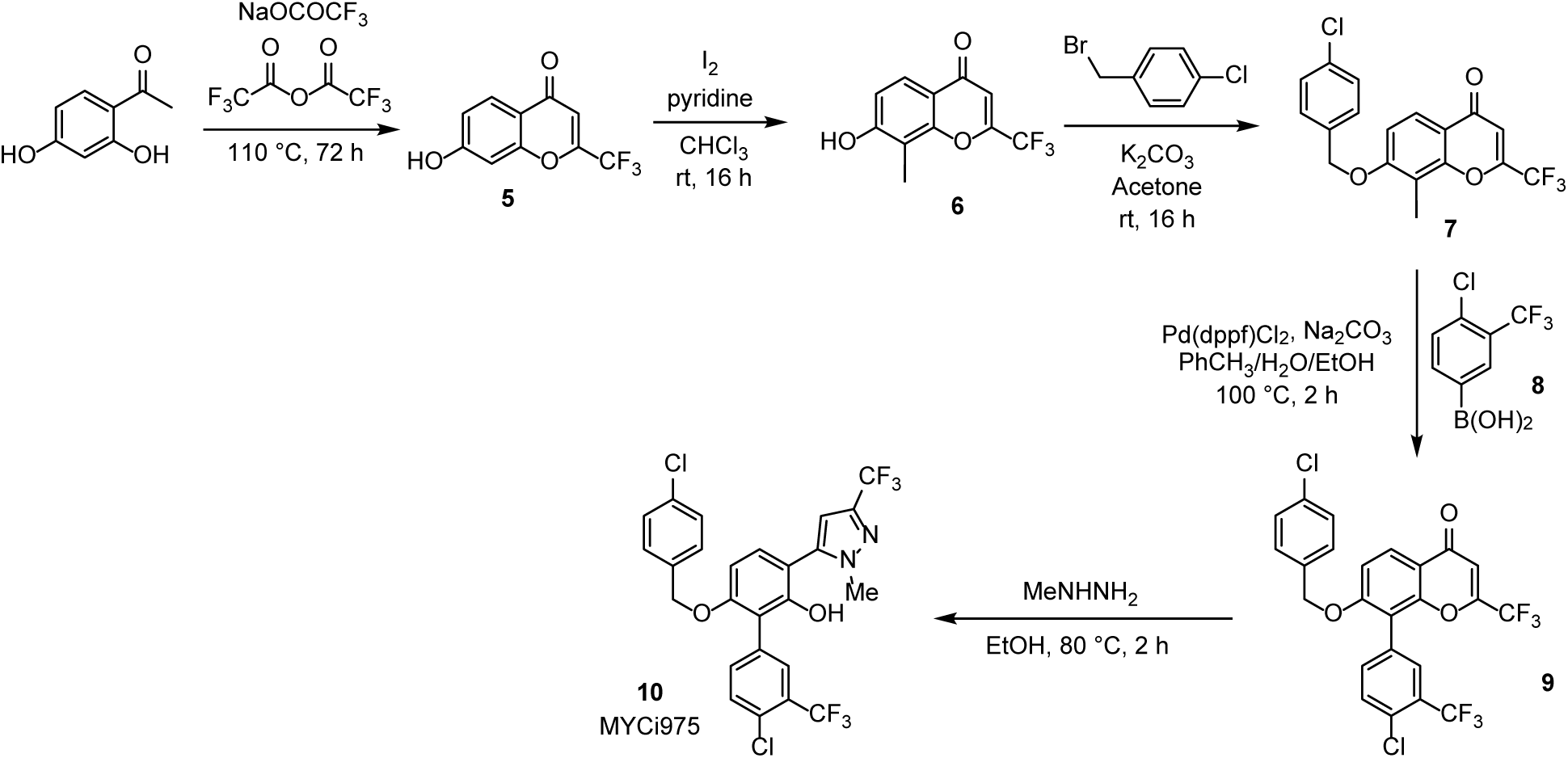

MYCi975 was prepared according to the reported procedure with minor modifications^69^.

Synthesis of compound **5**: To an oven-dry Schlenk flask (250 mL) was charged with (2,4- dihydroxyphenyl)ethan-1-one (5.00 g, 32.9 mmol, 1 equiv), sodium trifluoroacetate (9.84 g, 72,4 mmol, 2.2 equiv), and trifluoroacetic anhydride (18.5 mL, 131.6 mmol, 4 equiv). The flask was then tightly sealed with a polytetrafluoroethylene (Teflon) screw, and the reaction mixture was stirred at 110°C for 72 hr. The resulting mixture was allowed to cool down to approximately 70°C, and 200 mL of EtOAc was added in three batches to dilute the crude material. The solution was then transferred into a 1000 mL Erlenmeyer flask and neutralized with saturated aqueous K_2_CO_3_ solution until pH reached approximately 7. The organic layer was separated, and the water layer was washed with EtOAc (150 mL × 3). The combined organic layers were dried over anhydrous Na_2_SO_4_, filtered, and concentrated *in vacuo* to approximately 200 mL. The flask was kept open at room temperature for 2–3 days until no more solid was precipitated. Compound **5** (3.55g, 47%) was obtained by vacuum filtration.

Synthesis of compound **6**: To a suspension of 5 (4.00 g, 17.4 mmol, 1 equiv) in CHCl_3_ (110 mL) was added molecular iodine (17.6 g, 69.5 mmol, 4 equiv) and pyridine (5.62 mL, 69.5 mmol, 4 equiv). The mixture was allowed to stir at room temperature for 16 hr, and the reaction was quenched by saturated Na_2_S_2_O_3_ solution (120 mL). The organic layer was separated, and the aqueous layer was extracted with CH_2_Cl_2_ (80 mL × 3). The combined organic layers were washed with brine, dried over anhydrous Na_2_SO_4_, and concentrated *in vacuo* to give compound **6** (5.33 g, 86%).

Synthesis of compound **7**: To a suspension of 6 (2.00 g, 5.60 mmol) in acetone (20 mL) was added 4-chlorobenzyl bromide (1.50 g, 7.28 mmol, 1.3 equiv) and K_2_CO_3_ (1.55 g, 11.2 mmol, 2 equiv). The mixture was allowed to stir at room temperature for 16 hr. Upon completion, the mixture was filtered, and the filtrate was collected and concentrated *in vacuo* to give the dark-brown solids. The solids were then suspended in 75 mL of water, and crude material of 7 was obtained after vacuum filtration. No further purification was required.

Synthesis of compound **9**: To a round-bottom flask (50 mL) was added 7 (1.50 g, 3.12 mmol, 1 equiv), boronic acid 8 (770 mg, 3.43 mmol, 1.1 equiv), sodium carbonate (661 mg, 6.24 mmol, 2 equiv), and Pd(dppf)Cl_2_ (228 mg, 0.312 mmol, 10 mol %). Toluene (10 mL), water (4 mL), and ethanol (2 mL) were then introduced into the flask, and the mixture was bubbled with nitrogen gas for 10 minutes. The reaction was allowed to stir at 100 °C for 2 hr. Upon completion, the solution was diluted by 75 mL of EtOAc and washed with 50 mL of saturated NH_4_Cl solution. Crude material was obtained after drying over anhydrous Na_2_SO_4_, filtration, and concentration *in vacuo*. After SiO_2_ chromatography (Hexanes : EtOAc = 4 : 1) pure product 9 was obtained (1.12 g, 67%). Synthesis of compound **10** (MYCi975): To a suspension of 9 (1.50 g, 2.81 mmol, 1 equiv) in ethanol (11 mL) was introduced methylhydrazine (444 µL, 8.43 mmol, 3 equiv). The mixture was allowed to reflux (80°C) for 2 hr. Crude material was obtained by concentration *in vacuo*, and after SiO_2_ chromatography (Hexanes : EtOAc = 20 : 1 to 2 : 1) pure compound **10** was obtained (631 mg, 40%).

### Characterization of MYCi975

HPLC 93.3% (*t_R_* = 8.10 min), 85% CH_3_OH in water (0.1% TFA), 1mL/min, 20 min); ^1^H NMR (500 MHz, CDCl_3_) 7.79 (d, *J* = 2.0 Hz, 1H), 7.63 (d, *J* = 8.0 Hz, 1H), 7.53 (dd, *J* = 8.0, 2.0 Hz, 1H), 7.31 – 7.30 (m, 2H), 7.21 (d, *J* = 8.5 Hz, 1H), 7.16 – 7.14 (m, 2H), 6.72 (d, *J* = 8.5 Hz, 1H), 6.57 (s, 1H), 5.09 (s, 1H), 5.05 (s, 2H), 3.81 (s, 3H); ^19^F NMR (471 MHz, CDCl_3_) -62.02 (s, 3F), -62.60 (s, 3F); HPLC–MS (ESI+) *m/z* 559.8 and 558.7 (M-H)^+^.

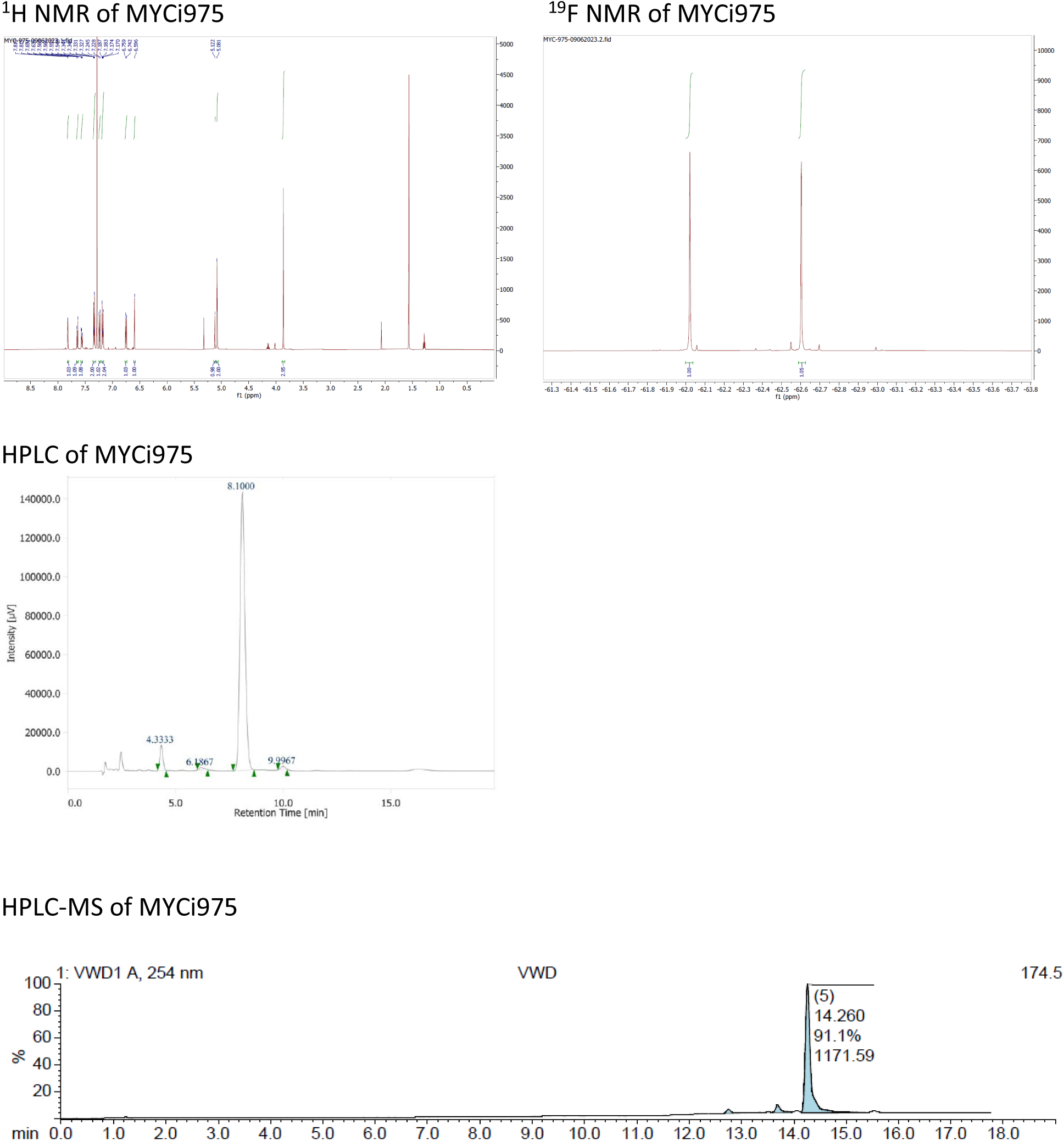

### Treatment of mice with MYCi975

CD45.2^+^ BM cells were harvested from Mx1-Cre^+/-^;Rosa26^LSL-MYC/LSL-MYC^ mice that was treated with pIpC 20 weeks prior. After RBC lysis, a total of 1 x 10^6^ cells were transplanted via tail vein into lethally irradiated CD45.1^+^ WT C57BL/6 recipient mice. Peripheral blood was collected at 4 weeks post-transplant, mononuclear cells were isolated following RBC lysis, and successful engraftment was confirmed by flow cytometry after staining cells with ZNIR viability dye, and anti-Ter-119, - CD45.1, and -CD45.2 antibodies. Mice were randomized to MYCi975 (100mg/kg/day, d1-5, 3 weeks on, 3 weeks off) *vs.* vehicle treatment. MYCi975 was initiated at 5 weeks post-transplant and drug was delivered via oral gavage. MYCi975 was dissolved in DMSO (final concentration 5%), mixed with corn oil, then vortexed multiple times until completely dissolved. Vehicle group was treated with corn oil. Treatment was continued until endpoints.

### Patient-derived xenograft studies

Primary BM cells from a TN-MF patient were collected under the TCC protocol. Somatic mutation in *JAK2, MPL,* or *CALR* gene was assessed by NGS. Trisomy 8 was assessed by both conventional karyotyping and FISH analyses. After isolating BM MNCs following SepMate^TM^ PBMC isolation protocol, cells were frozen in DMEM media supplemented with 20% FBS and 20% DMSO until used. After thawing BM MNCs were cultured in serum free SFEM II media (STEMCELL Technologies) supplemented with Pen-Strep (50 U/ml), human SCF (100 ng/ml), human TPO (100 ng/ml), and human FLT3L (100 ng/ml) for 16 hours. A total 1.6 x 10^5^ cells (in 30-µl of PBS) were transplanted via intra-tibial injection into sublethally irradiated (200 cGy) NSG-SGM3 mice. Engraftment of MF cells were confirmed by measuring human CD45^+^ cells in peripheral blood by flow cytometry at 9 weeks post-transplant. Mice were then randomized to vehicle, MYCi975 (100mg/kg, po, once a day, d1-5, 3 weeks on, 2 weeks off), or ruxolitinib (180mg/kg, po, once a day, d1-5, 3 weeks on, 2 weeks off) treatment.

### Statistical analysis

Under the assumption of independent variables, normal distribution, and equal variance of samples, statistical significance was assessed using unpaired two-tailed Student’s *t*-test for in vitro and ex vivo experiments. Error bars presented in the figures indicate the mean ± SEM. The statistical parameters are described in the individual figure legends. For survival analyses, LFS and OS were estimated using the Kaplan-Meier method and compared using a log-rank test. Statistical analyses were performed using SPSS v24.0 and GraphPad Prism 7. A *p*-value less than 0.05 was considered statistically significant.

### Data Sharing Statement

scRNA-Seq data from mouse and human BM cells have been deposited in the GEO under accession files GSE240963 and GSE242730, respectively.

## AUTHORS’ DISCLOSURE

D.J.M. has received funding from Merck Pharmaceuticals and PUMA Biotech for work unrelated to this project. A.T.K. received research funding from Bristol Myers Squibb, Novartis, Morphosys, GlaxoSmithKline, Janssesn. A.T.K. received honoraria from Abbvie, GlaxoSmithKline, MorphoSys, Incyte, Bristol Myers Squibb, CTI Biopharma, Kartos, and Karyopharm.

## AUTHOR CONTRIBUTIONS

Study conception and design: N.D.V., J.L.C., and S.Y.; performed experiments, collected, and assembled the data: N.D.V., X.Y., A.T.K., J.M., C-H.C., R.S., H.V.N., A.A.N., P.C.P., D.H.L., T.N.R., Q.M., R.S.K., and S.Y.; analyzed and interpreted the data: N.D.V., X.Y., C-H.C., O.C., Q.M., L.Z., D.J.M., S.H.K., J.L.C., and S.Y.; writing and/or revision of the manuscript: N.D.V., X.Y., H.L., D.L., S.H.K., J.L.C., and S.Y.; review of manuscript: all authors reviewed the manuscript; administrative, technical or material support: N.D.V., X.Y., S.S., H.V.N., E.E., S.N., R.B.F., H.L., S.C., D.J.M., D.L., S.H.K., J.L.C., and S.Y.; study supervision: N.D.V. and S.Y.

## ACKNOWLEDGEMENTS

The authors thank Jodi Kroeger, Dr. Neel Chaudhary, and the Moffitt Flow Cytometry Core; Sean Yoder, Dr. Chaomei Zhang, Dr. Lan Zhang, Tania Mesa, and the Moffitt Genomics Core for their help with scRNA-seq analyses; Dr. Eric Padron for providing Scl-CreERT mice; Dr. Peter Papenhausen and Dr. Liu Kenian for their assistance with conventional karyotyping and FISH studies; Jodi Balasi, Noel Clark, and the Moffitt Histology Core for their assistance with immunohistochemistry staining; Yamila Caraballo Perez, Janis De La Iglesia, and the Moffitt Clinical Pathology Lab for their help with reticulin, trichrome, and immunohistochemistry staining.

This work was supported in part by Deutsche Krebsilfe grant 109220 (D.J.M.), the program grant 1016701 from the Australian National Health and Medical Research Council (NHMRC) (S.C.), the Cortner-Couch Endowed Chair for Cancer Research from the University of South Florida School of Medicine (J.L.C.), NIH grant K08 CA237627 (S.Y.), an American Society of Hematology (ASH) Research Training Award for Fellows (RTAF) (S.Y.), an ASH Scholar Award for Fellow (S.Y.), an ASH Scholar Award for Fellow to Faculty (N.V.D.), a Career Development Award from the American Society of Clinical Oncology (ASCO) (S.Y.), a Miles for Moffitt grant from the H. Moffitt Cancer Center & Research Institute (S.Y. and Q.M.), a research grant from the Graduate Medical Education (GME) at the University of South Florida (S.Y. and P.C.P.), a research grant from the Clinical Science Division at the H. Lee Moffitt Cancer Center & Research Institute (S.Y.), and NCI Comprehensive Cancer Grant P30-CA076292 to the H. Lee Moffitt Cancer Center & Research Institute.

## NOTE

Supplemental data for this article are available at Cancer Discovery Online.

## Supplemental Information

### SUPPLEMENTAL FIGURE LEGENDS

**Figure S1.**
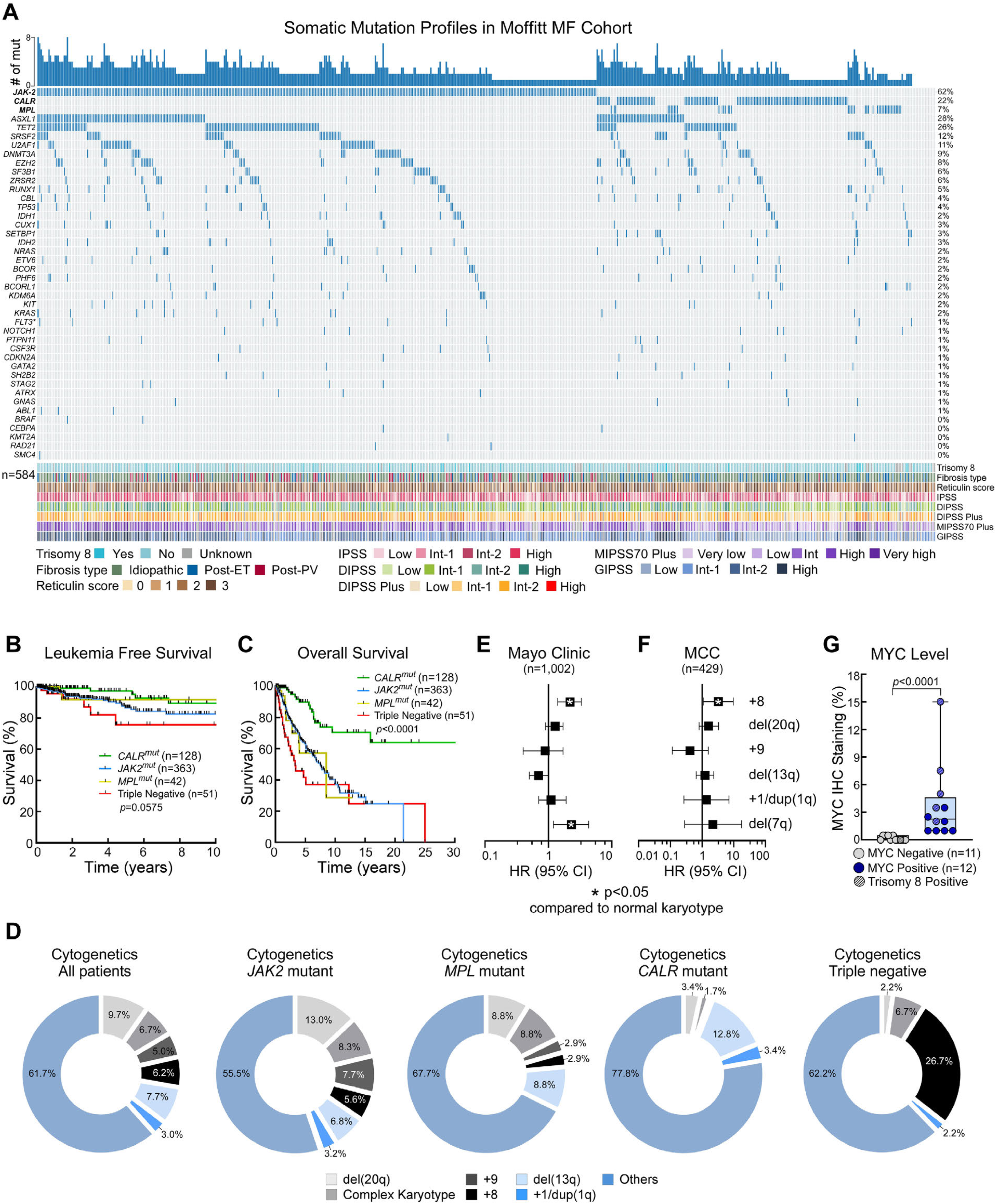
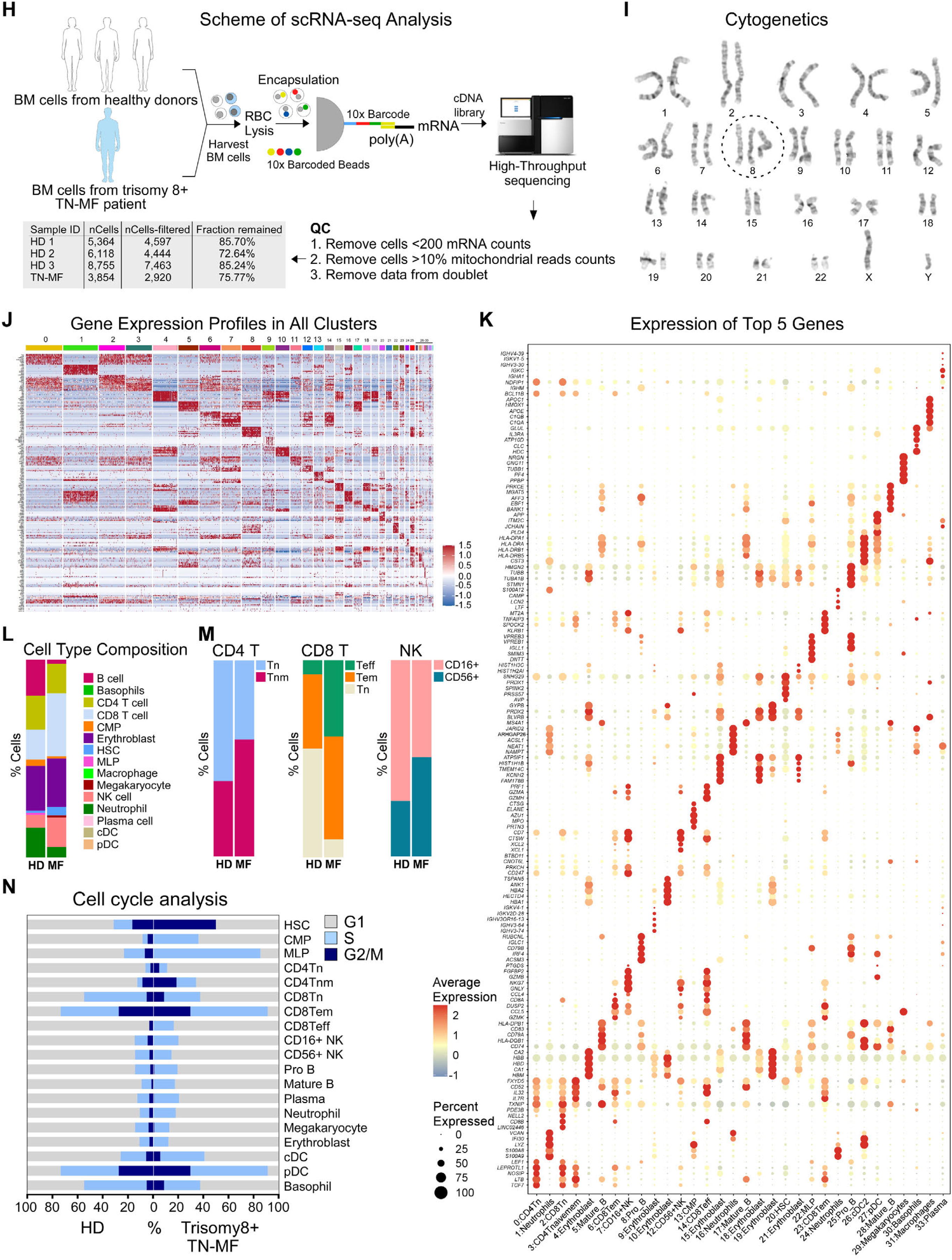
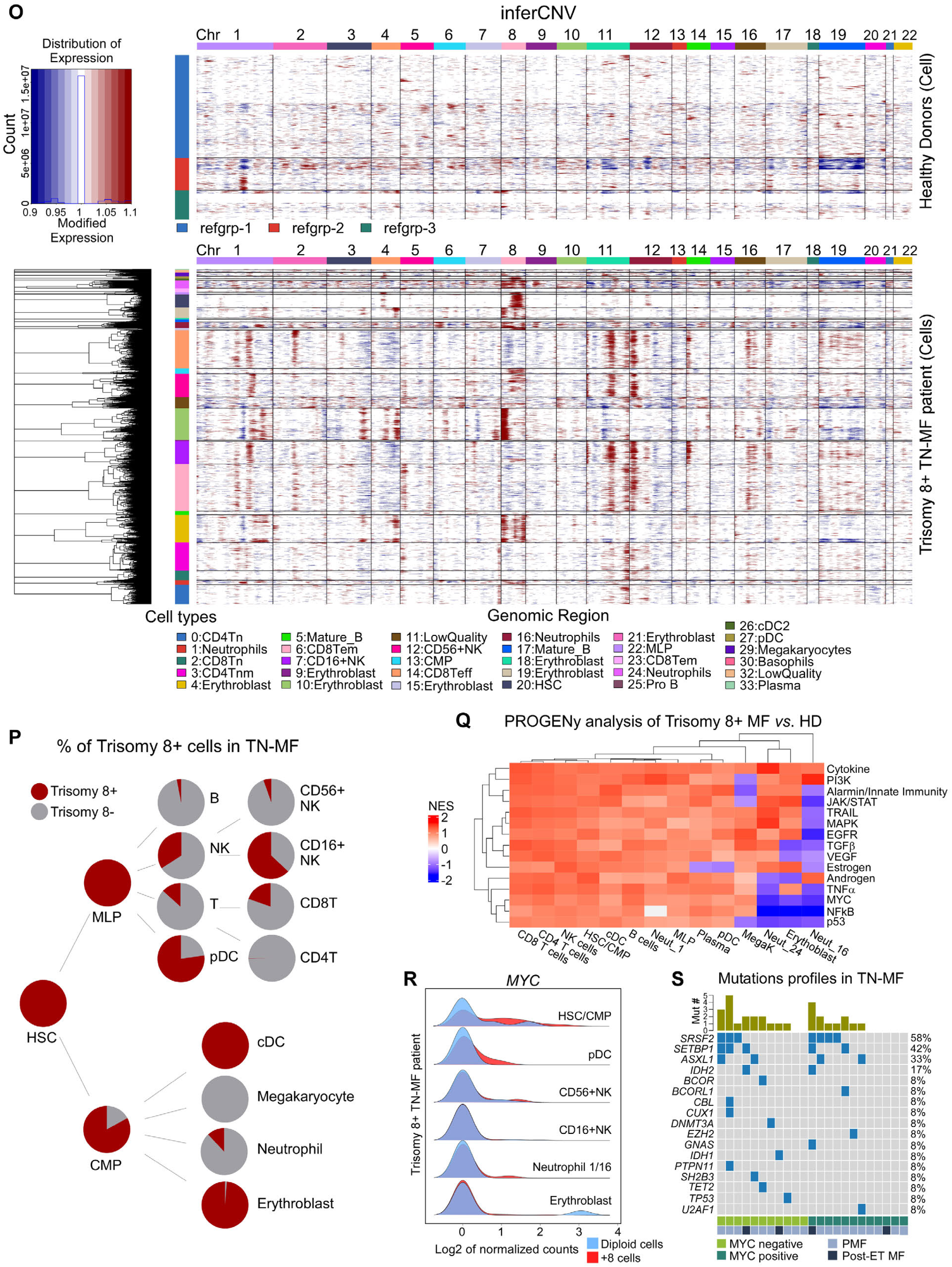
Landscape of somatic mutations and cytogenetics in Moffitt MF cohort. **A,** Somatic mutation profiles in Moffitt MF patients. Individual columns and rows represent each patient and somatic mutation, respectively. Positive mutations are highlighted in dark blue. Numbers of total mutations in individual patients and percentages of somatic mutations in the total number of patients are described in the top and right side of the plot, respectively. Trisomy 8 status, subtypes of MF, reticulin fibrosis score, and prognostic scores are described at the bottom of the plot. Demographic profiles and laboratory parameters are described in Table S1. **B-C,** Kaplan-Meier curves showing the LFS (B) and OS (C). **D,** Percentages of cytogenetic abnormalities (top 5 most frequent aberrations) and complex karyotype (defined as at least 3 concurrent chromosomal abnormalities) in individual molecular subtypes and all MF patients. **E-F,** Forest plots showing hazard ratio (HR) and 95% confidence intervals associated with individual cytogenetic abnormalities in the Mayo Clinic (E) and Moffitt MF (F) cohorts, respectively. Patients with more than one chromosomal abnormality were excluded from the analyses. Data in (E) were re-analyzed using the original published data^38^. **G,** Levels of MYC protein expression in BM cells of TN-MF patients. MYC positivity was defined as at least 1% of cells that express MYC protein. **H,** Schematic diagram describing scRNA-seq analyses of human BM cells. **I,** Conventional karyotyping of TN-MF patient BM cells used in scRNA-seq analysis. **J,** Heatmaps depicting differentially expressed genes between healthy donors (n = 3) vs. trisomy 8+ TN-MF patient in the indicated individual clusters. **K,** Cell type markers (top 5 genes) in the indicated individual clusters. **L-M,** Cell type composition of BM cells in healthy donors *vs.* trisomy 8+ TN-MF patient. **N,** Percentages of cells in G1, S, and G2/M phase of cell cycle in individual major cell types of healthy donors *vs.* trisomy 8+ TN-MF patient. **O,** inferCNV analysis based on scRNA-seq data. Top and bottom panels represent data from normal donors (used as reference) and trisomy 8+ TN-MF, respectively. Columns represent individual chromosomes and rows represent major cell types. **P,** Pie charts showing percentages of trisomy 8+ *vs.* diploid cells in individual major cell types in trisomy 8+ TN-MF patient BM.**Q,** PROGENy analysis comparing trisomy 8+ TN-MF *vs.* healthy donors BM cells. Individual columns and rows represent major cell types and pathways that are known to contribute to MPN pathogenesis. **R,** Ridgeline plots of *MYC* mRNA levels in diploid (blue) *vs.* trisomy 8+ (red) BM cells for each major cell type of the trisomy 8+ TN-MF patient. **S,** Somatic mutation profiles in MYC negative (n = 11) *vs.* positive (n = 12) TN-MF patients. Individual columns and rows represent each patient and somatic mutation, respectively. Positive and negative mutations are highlighted in dark blue and grey, respectively. Number of total mutations in individual patients and the percentages of somatic mutations in total patients are described in the top and right side, respectively. Subtypes of MF and MYC status are described at the bottom of the plot. Abbreviations: MF, myelofibrosis; Triple negative MF, TN-MF; LFS, leukemia free survival; OS, overall survival; MCC, Moffitt Cancer Center; IPSS, International Prognostic Scoring System; DIPSS, Dynamic IPSS; MIPSS70, Mutation-Enhanced IPSS; GIPSS, Genetically Inspired Prognostic Scoring System.

**Figure S2.**
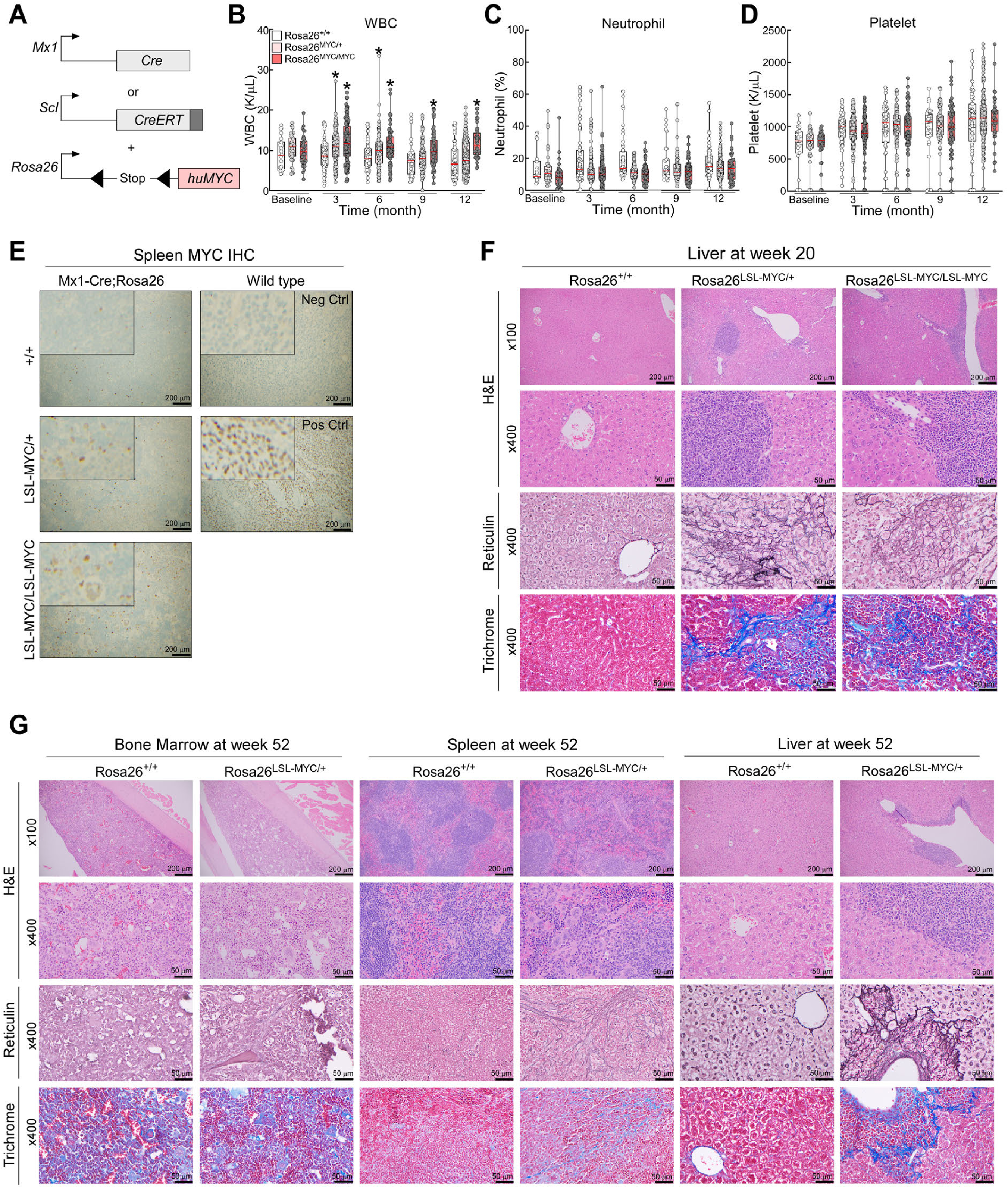
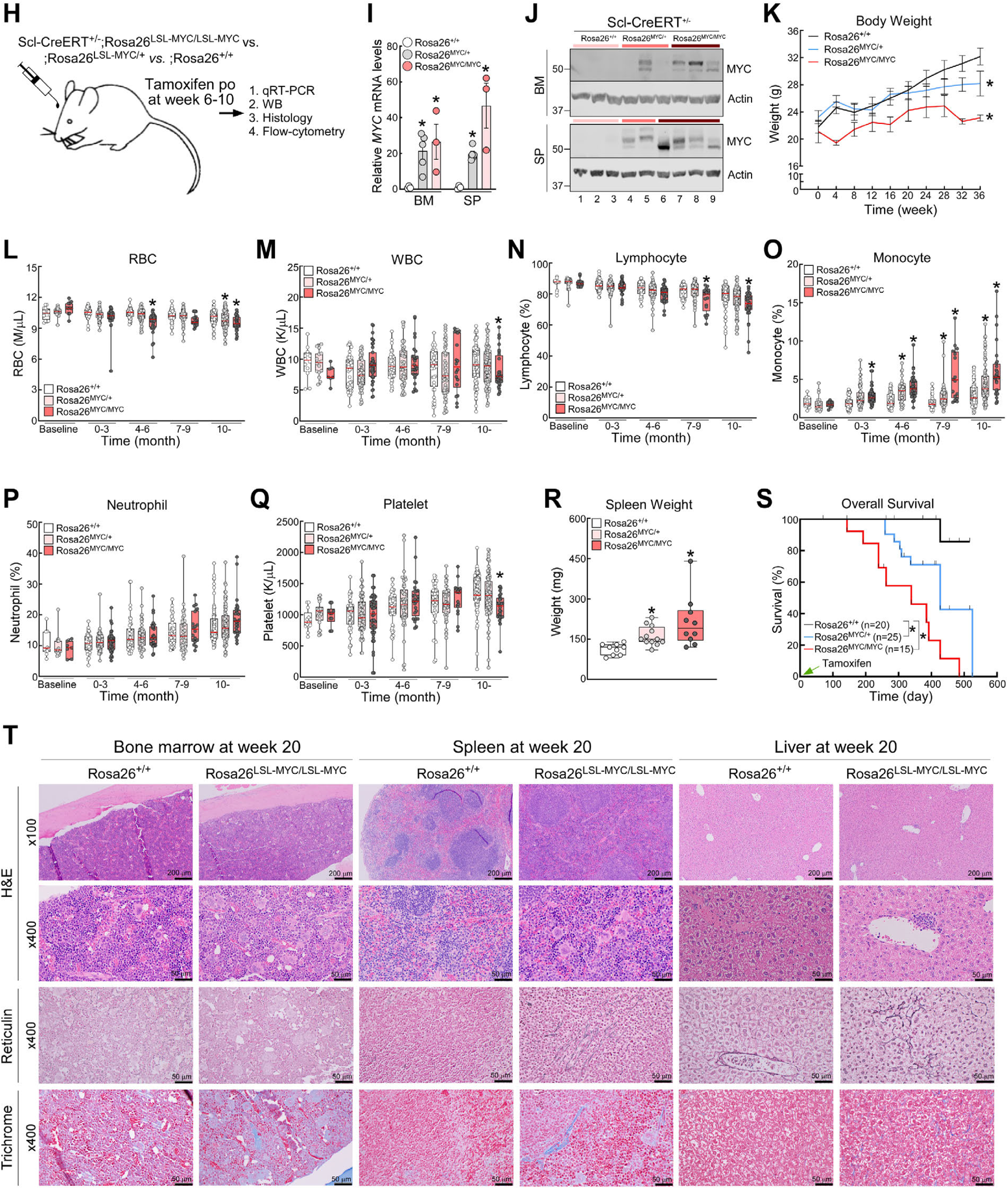
Effects of enforced MYC expression in HSCs in vivo. **A,** Schematic of the *Mx1- Cre, Scl-CreERT*, and *Rosa26-LSL-MYC* alleles. **B-D,** PB CBC analyses of white blood cells (WBC) (B), neutrophils (C), and platelets (D). Baseline CBC was performed 1 week prior to pIpC treatment. **E,** MYC IHC staining of spleen tissue. **F,** H&E, reticulin, and trichrome stained images of liver tissues at week 20 following pIpC injection. **G,** H&E, reticulin, and trichrome stained images of BM, spleen, and liver at week 52 post-pIpC treatment. Demographic profiles and laboratory parameters are described in Table S4. **H,** Schematic of in vivo studies using Scl-CreERT2^+/-^;Rosa26^LSL-MYC/LSL-MYC^ mice. **I-J,** MYC mRNA (I) and protein (J) levels in BM and spleen of Scl-CreERT^+/-^;Rosa26^+/+^, Scl-CreERT^+/-^;Rosa26^LSL-MYC/+^, and Scl-CreERT^+/-^;Rosa26^LSL-MYC/LSL-MYC^ mice, respectively. **K,** Comparison of body weight between the indicated experimental groups. **L-Q,** PB CBC analyses of RBC (L), WBC (M), lymphocytes (N), monocytes (O), neutrophils (P), and platelets (Q). Baseline CBC was performed 1 week prior to tamoxifen treatment. **R,** Spleen weight at endpoints. Scl-CreERT^+/-^;Rosa26^+/+^ mice were sacrificed as control when Scl-CreERT^+/-^;Rosa26^LSL-MYC/+^ and Scl-CreERT^+/-^;Rosa26^LSL-MYC/LSL-MYC^ mice were at their endpoints. Endpoints criteria is described in “Animal studies” section. **S,** Kaplan-Meier curves of OS, calculated from the date of tamoxifen treatment. **T,** H&E, reticulin, and trichrome stained images of BM, spleen, and liver at week 20 following tamoxifen treatment. Demographic profiles and laboratory parameters are described in Table S6. Error bars in (I) and (K) indicate mean ±SEM of at least 3 independent mice. Box plots in panels (B-D) and (L-Q) represent data from 38-54 and 15-25 mice, respectively, in each experimental group. * In panels (B, I, K-O, and Q-S), *P*<0.05 compared with control group.

**Figure S3.**
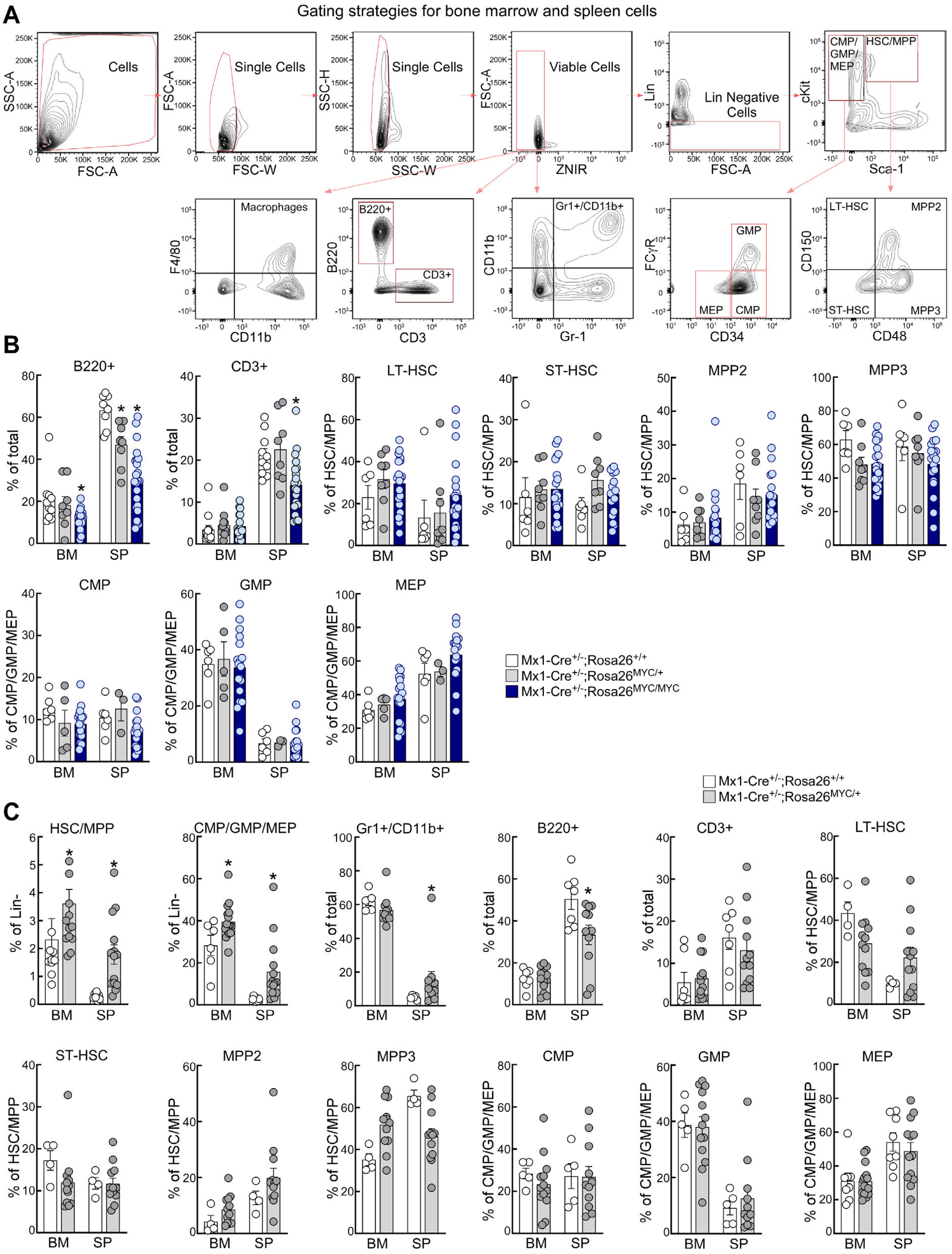
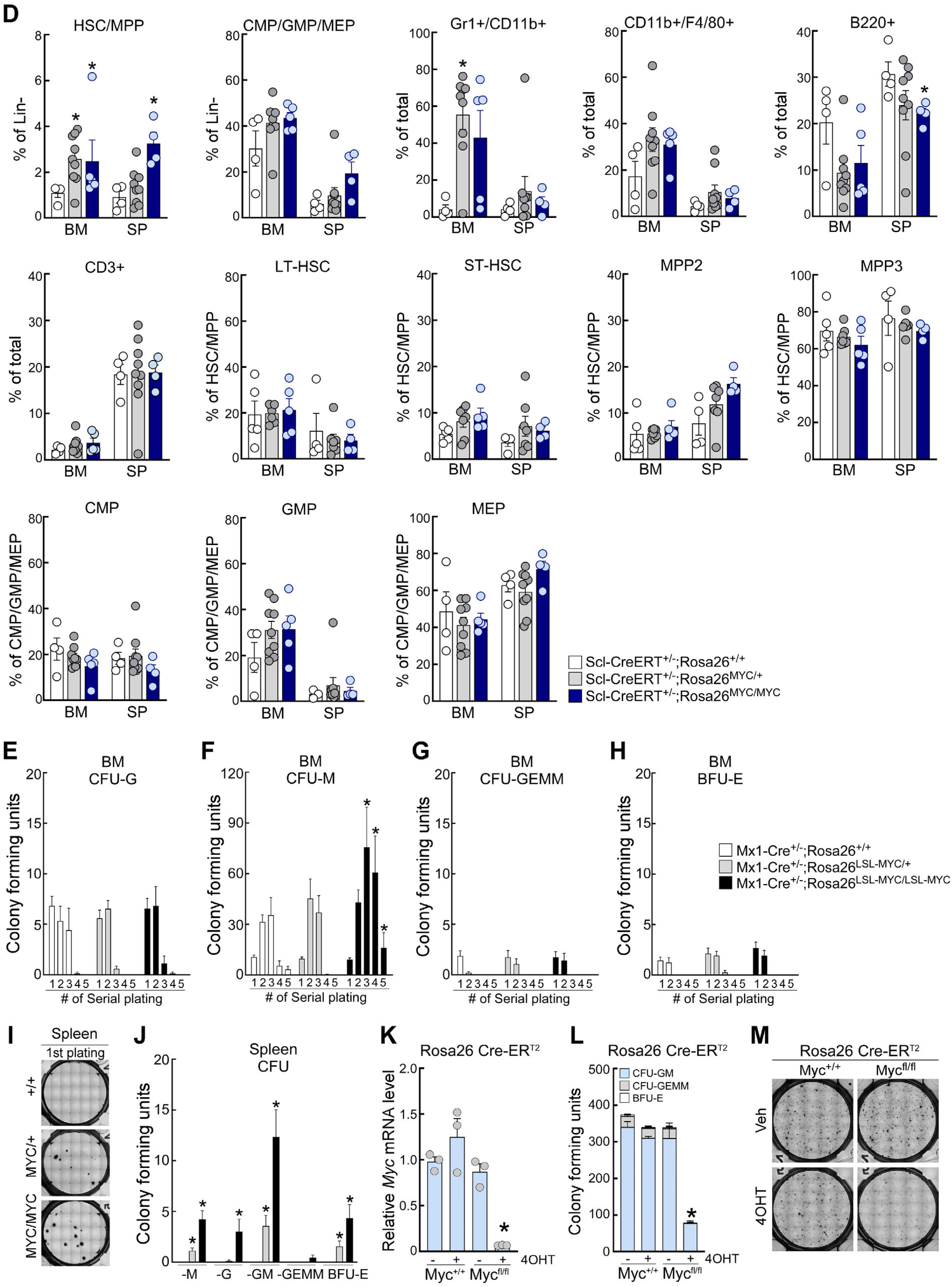
MYC-induced changes in hematopoietic sub-populations and colony forming potential. **A,** Gating strategies used to quantify hematopoietic subpopulations. **B,** Percentages of B220^+^ B-cells, CD3^+^ T-cells, LT-HSC, ST-HSC, MPP2, MPP3, CMPs, GMPs, and MEPs in BM or spleen of the indicated groups before 45 weeks from pIpC treatment. **C,** Percentages of individual hematopoietic subpopulations in Mx1-Cre^+/-^;Rosa26^+/+^ and Mx1-Cre^+/-^;Rosa26^LSL-MYC/+^ mice, respectively, after 45 weeks from pIpC treatment. **D,** Comparison of individual hematopoietic cell types in BM and spleen of Scl-CreERT^+/-^;Rosa26^+/+^, Scl-CreERT^+/-^;Rosa26^LSL-MYC/+^ and Scl-CreERT^+/-^;Rosa26^LSL-MYC/LSL-MYC^ mice. **E-H,** Serial colony forming assays using primary BM cells from Mx1- Cre^+/-^;Rosa26^+/+^, Mx1-Cre^+/-^;Rosa26^LSL-MYC/+^ and Mx1-Cre^+/-^;Rosa26^LSL-MYC/LSL-MYC^ mice at 20 weeks post-pIpC. A total 1x10^4^ cells were plated from each experimental group, cultured for 10 days, and CFU-G (E), CFU-M (F), CFU-GEMM (G), and BFU-E (H) were then counted manually. **I-J,** Serial colony forming assays using the indicated primary spleen cells. **K-M,** Colony forming assays using primary BM cells harvested from 7-week old Rosa26-Cre-ERT2^+/-^;*Myc^+/+^* and Rosa26-Cre-ERT2^+/-^;*Myc*^fl/fl^ mice. Cells were cultured ex vivo with vehicle or 4-OHT (1µM) for 10 days, and then assessed for colony forming units. Knockout of *Myc* was confirmed by qRT-PCR assays (K). Error bars in (B-H and J-L) indicate mean ±SEM of at least 3 independent mice. * in (B-D, F, and J-L), *P*<0.05 compared with control group.

**Figure S4.**
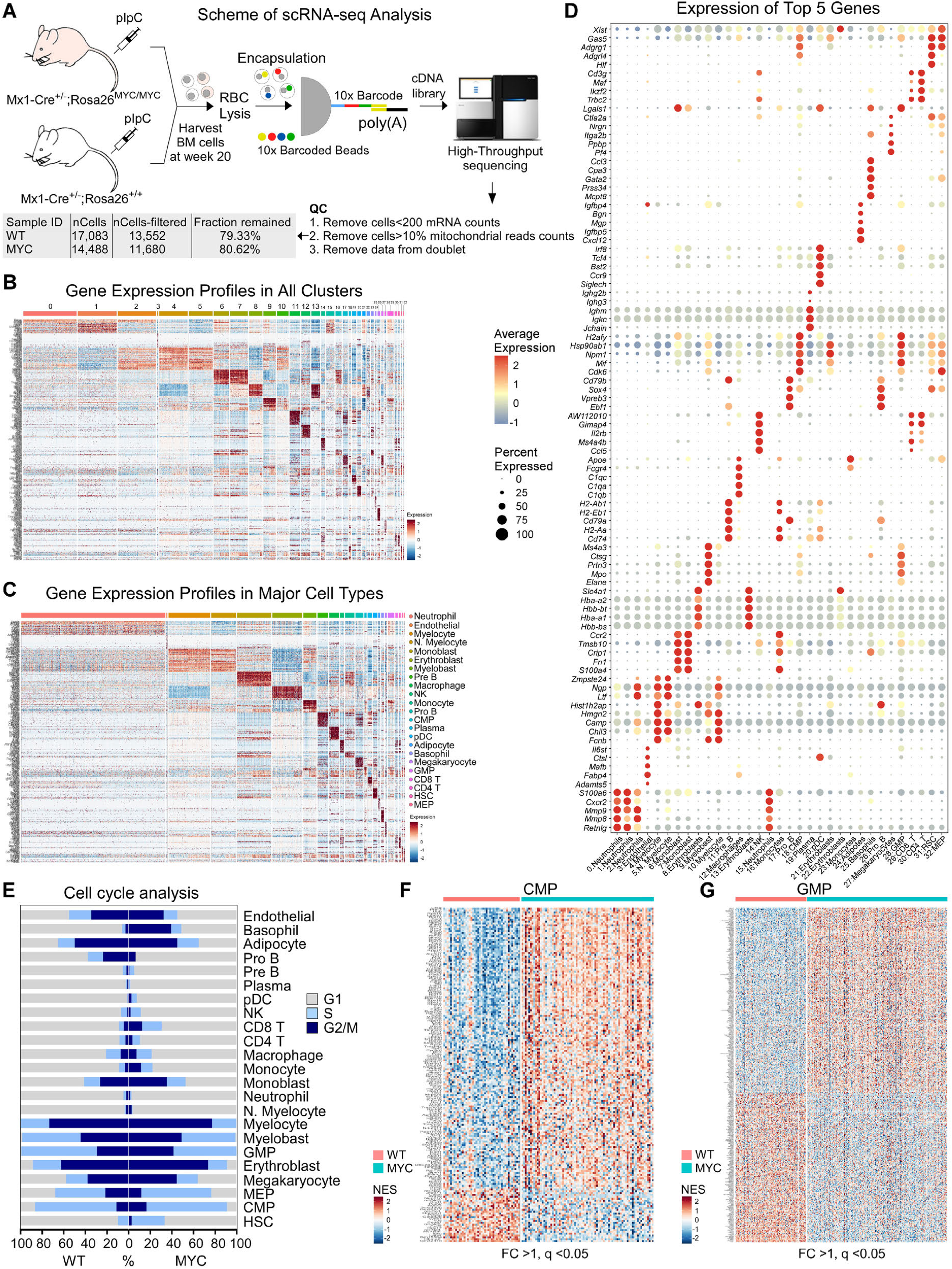
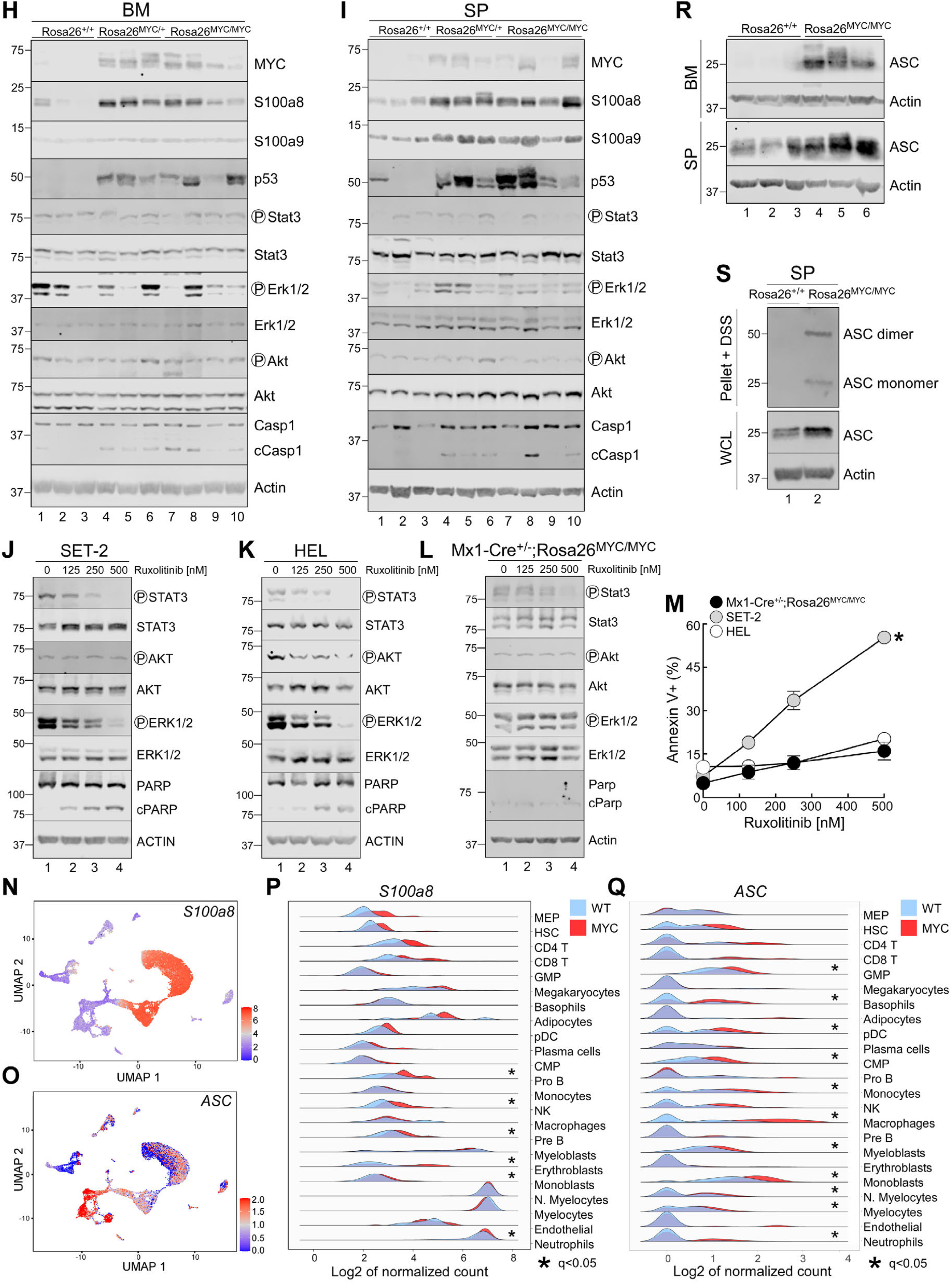
JAK/STAT, PI3K/AKT, MEK/ERK, and alarmin pathways in MYC-driven MF. **A,** Schematic of scRNA-seq analyses of mouse BM cells. **B-C,** Heatmaps depicting differentially expressed genes between Mx1-Cre^+/-^;Rosa26^+/+^ control *vs.* Mx1-Cre^+/-^;Rosa26^LSL-MYC/LSL-MYC^ mouse in individual clusters (B) and major cell types (C). **D,** Cell type markers (top 5 genes) in individual clusters. **E,** Percentages of cells in G1, S, and G2/M phase of cell cycle in each major cell type of Mx1-Cre^+/-^;Rosa26^LSL-MYC/LSL-MYC^ (MYC) *vs.* control (WT) mice. **F-G,** Heatmaps showing differentially expressed genes (log2 fold change>0.25, q<0.05) between Mx1-Cre^+/-^;Rosa26^+/+^ *vs.* Mx1-Cre^+/-^;Rosa26^LSL-MYC/LSL-MYC^ mouse in the CMPs (F) and GMPs (G) clusters. **H-I,** Immunoblotting of BM (H) and spleen cells (I) harvested from Mx1-Cre^+/-^;Rosa26^+/+^, Mx1-Cre^+/-^;Rosa26^LSL-MYC/+^ and Mx1- Cre^+/-^;Rosa26^LSL-MYC/LSL-MYC^ mice. **J-M,** SET-2, HEL, and primary BM cells harvested from Mx1-Cre^+/-^;Rosa26^LSL-MYC/LSL-MYC^ mice were incubated with vehicle *vs.* ruxolitinib as indicated for 48 hr, and then assessed for changes in phospho-STAT3, -ERK1/2, -AKT, and cleaved PARP protein levels (J-L) and apoptosis (M, % Annexin V^+^ cells). **N-Q,** *S100a8* and *ASC* mRNAs levels in single cell are projected onto the UMAP plots (N, O, respectively) and changes in these genes in the individual major cell types in MYC (red) *vs.* control (blue) BM cells are presented in the ridgeline plots (P, Q). **R,** Immunoblotting of BM (top panel) and spleen cells (bottom panel) harvested from Mx1- Cre^+/-^;Rosa26^+/+^ and Mx1-Cre^+/-^;Rosa26^LSL-MYC/LSL-MYC^ mice. **S,** ASC speck cross-linking using spleen cells harvested from Mx1-Cre^+/-^;Rosa26^+/+^ and Mx1-Cre^+/-^;Rosa26^LSL-MYC/LSL-MYC^ mice. See method for the details. Actin blots in panels (H-L) were from the same samples run on different gels. Error bars in (M) indicate mean ±SEM of 3 independent assays. * In (M, P, and Q), *P*<0.05 compared with control group.

**Figure S5.**
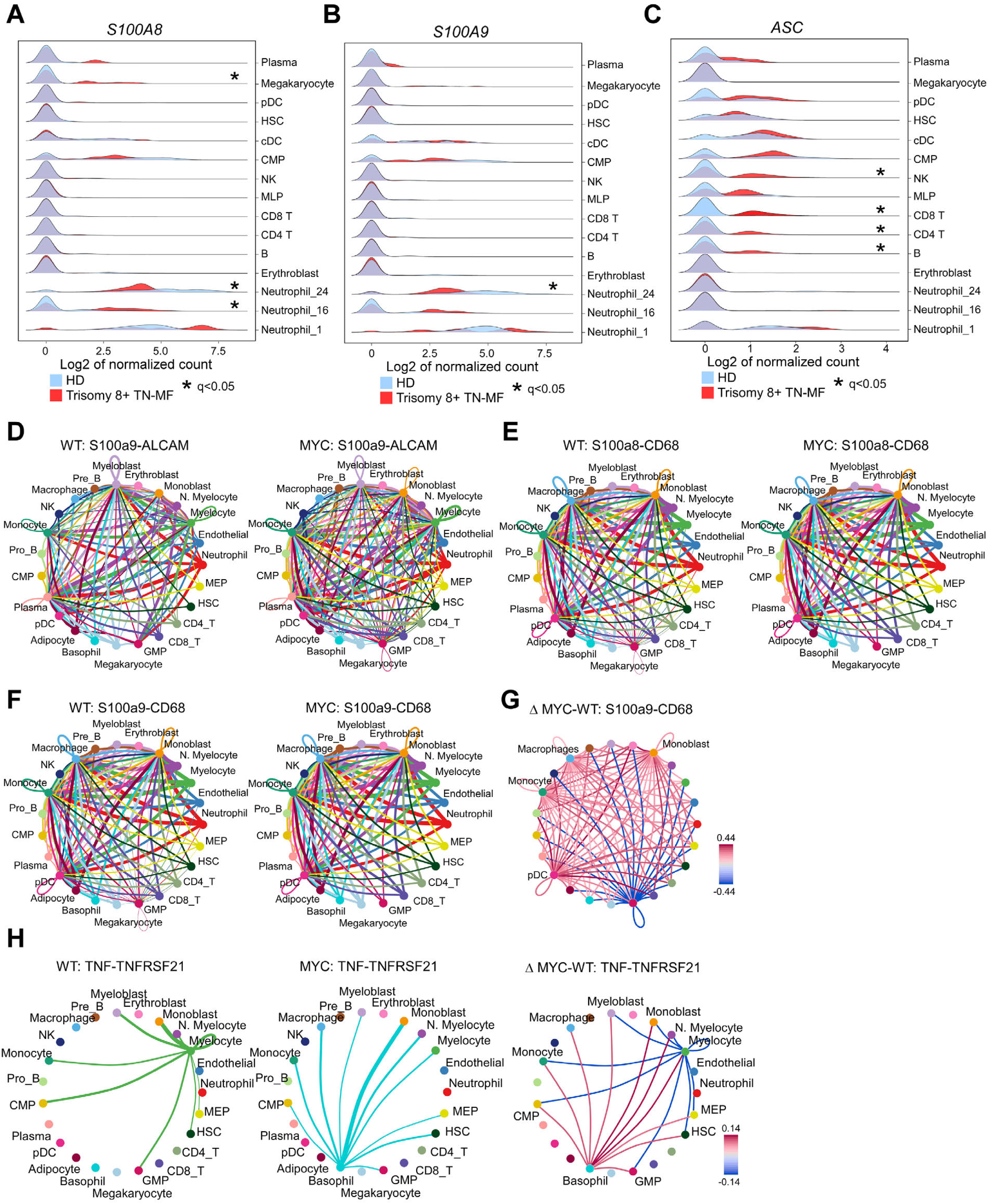
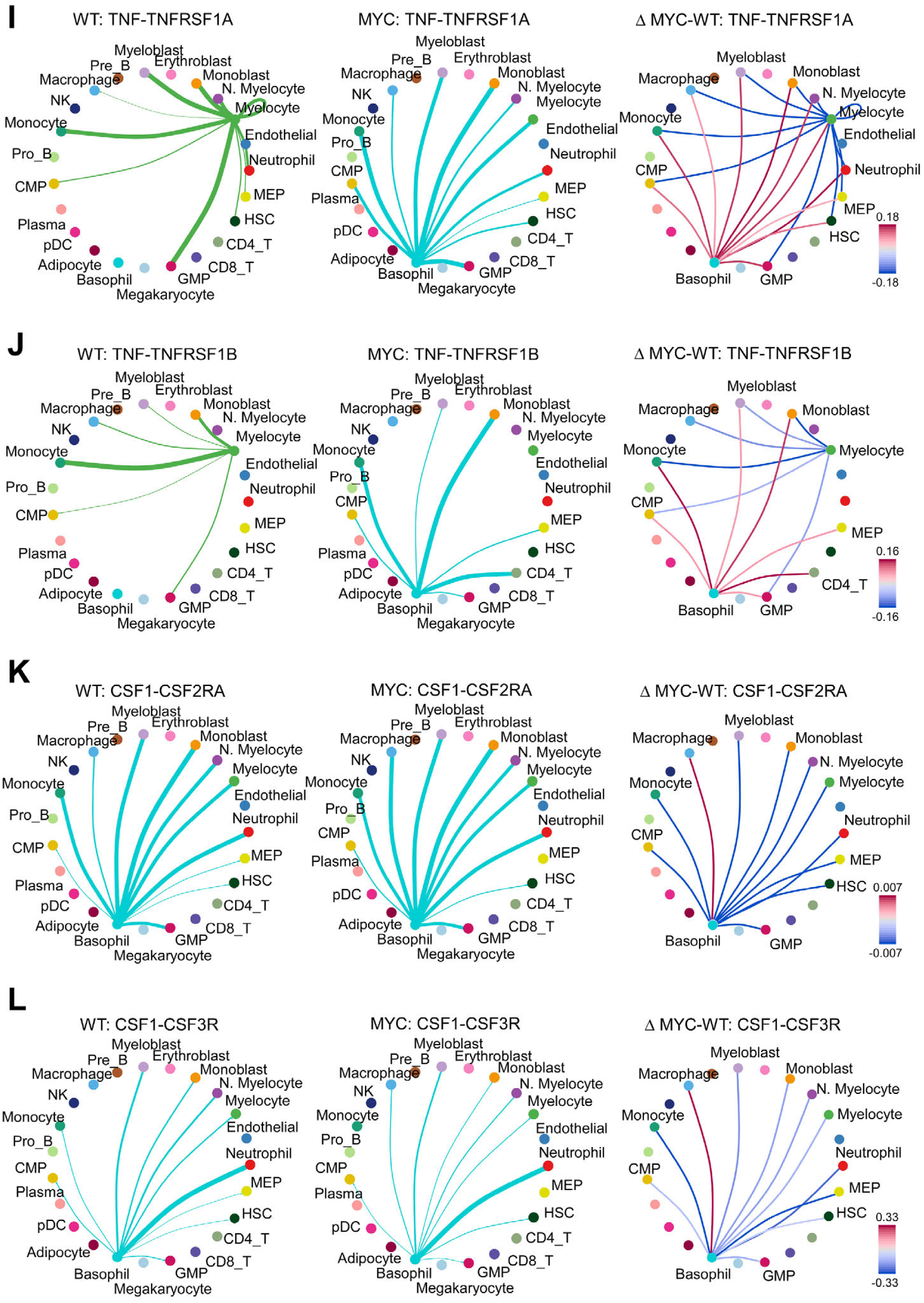
Changes in signaling in BM of trisomy 8+ TN-MF patient and MYC MF mice. **A-C,** Ridgeline plots comparing *S100A8/A9* and *ASC* mRNA levels in each major BM cell type of trisomy 8+ TN-MF patient *vs.* normal healthy donors. **D-G,** Network plots of interactions of ligands (S100a8 or S100a9) and receptors (ALCAM or CD68). Ligand signals originating from individual major cell types are presented in different colors. Thicker lines in (D-F) indicate a more frequent interaction between cell types. Differences in intensity of interactions are presented in (G). Higher and lower signal activity in MYC homozygous cells (compared to WT) are presented in red and blue, respectively. **H-L,** Network plots of interactions of ligands (TNF-α or CSF-1) and receptors (TNFRSF21, TNFRSF1A, TNFRSF1B, CSF2RA, or CSF3R) in mouse BM cells. Differences in intensity of interactions are presented in the third plots of the individual panels. Higher and lower signal activity in MYC cells (compared to WT) are presented in red and blue, respectively. * in panels (A-C), *P*<0.05 compared with control group.

**Figure S6.**
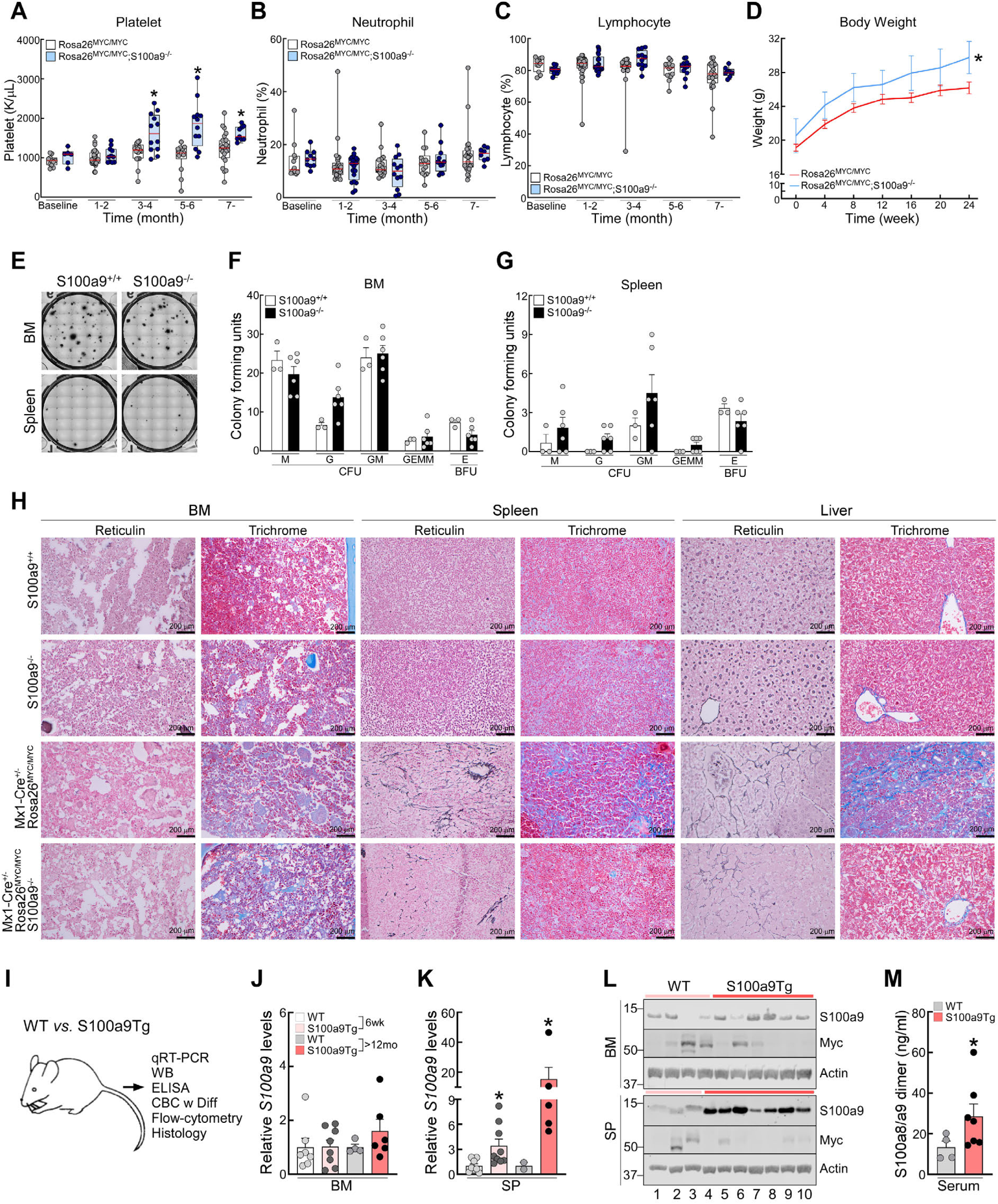
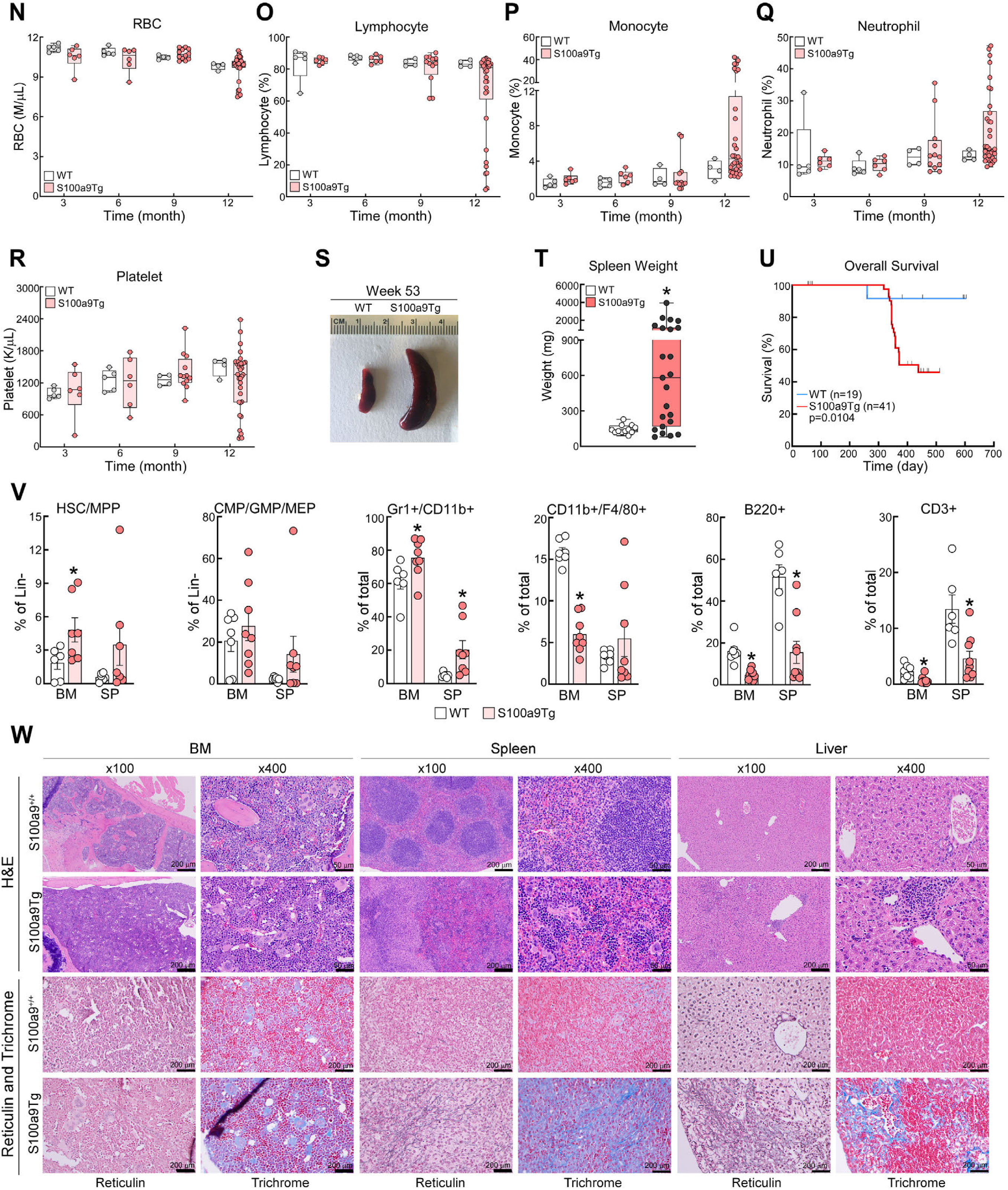
Effects of silencing in MYC-driven MF or overexpression of S100a9. **A-C,** PB CBC analyses in Mx-Cre^+/-^;Rosa26^LSL-MYC/LSL-MYC^ and Mx-Cre^+/-^;Rosa26^LSL-MYC/LSL-MYC^;*S100A9*^-/-^ mice following pIpC treatment. Baseline CBC was performed 1 week prior to pIpC. **D,** Comparison of body weight of Mx-Cre^+/-^;Rosa26^LSL-MYC/LSL-MYC^ and Mx-Cre^+/-^;Rosa26^LSL-MYC/LSL-MYC^;*S100A9*^-/-^ mice following treatment with pIpC. **E-G,** Comparison of colony forming units between WT *vs. S100a9*^-/-^ mice. **H,** Reticulin and trichrome stained images of BM, spleen, and liver in the indicated mice. Demographic profiles and laboratory parameters are described in Table S11. **I,** Schematic of in vivo S100a9Tg mouse studies. **J-K,** Levels of S100a9 mRNA in BM (J) and spleen (K) cells of S100a9Tg *vs*. control mice at indicated time points. **L,** Levels of S100a9 protein in BM (top panels) and spleen (bottom panels) of S100a9Tg mice at age >12 months. Age-matched control mice were sacrificed at the same time for paired analysis. **M,** Serum levels of S100a8/a9 heterodimers in S100a9Tg *vs*. control mice at age >12 months. **N-R,** PB CBC analyses of RBC (N), lymphocytes (O), monocytes (P), neutrophils (Q), and platelets (R) in S100a9Tg *vs*. control mice at indicated time points. **S-T,** Spleen weight of S100a9Tg mice at age >12 months. Age-matched control mice were sacrificed as control. **U,** Kaplan-Meier curves of OS, calculated from date of birth. **V,** Percentages of HSCs and MPPs, myeloid progenitors, Gr1^+^/CD11b^+^ mature myeloid cells, macrophages, B-cells, and T-cells in each cohort. **W,** H&E, reticulin, and trichrome stained images of BM, spleen, and liver of S100a9Tg *vs.* control mice at age >12 months. Clinical parameters are described in Table S12. Actin blots in (L) were from the same samples run on different gels. Box plots in (A-C, N-R, and T) represent data from at least 5 mice in each group. Error bars in (D, F, G, J, K, M, and V) indicate mean ±SEM of at least 5 independent assays. * in (A, D, K, M and V), *P*<0.05 compared with control group.

**Figure S7.**
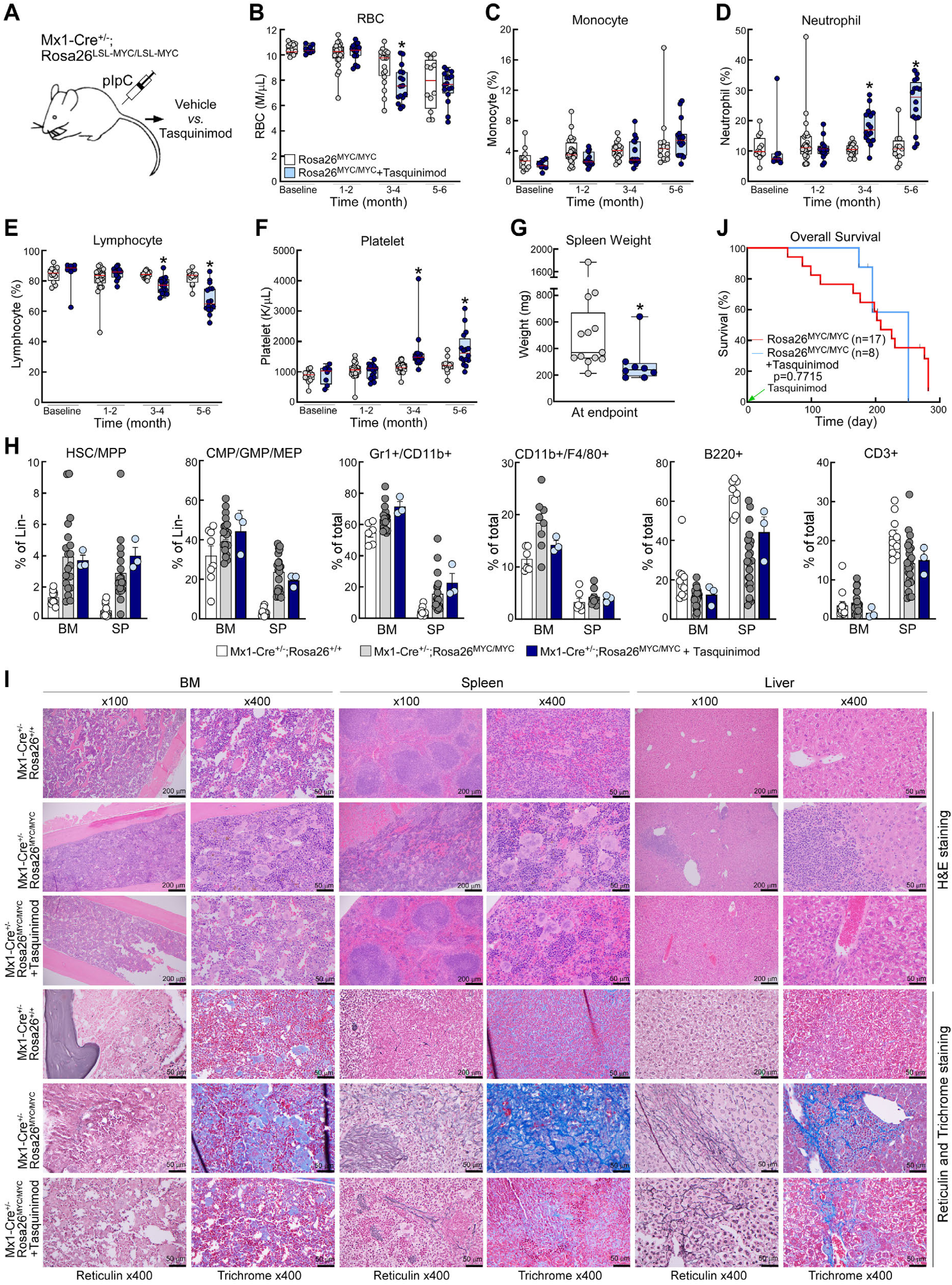
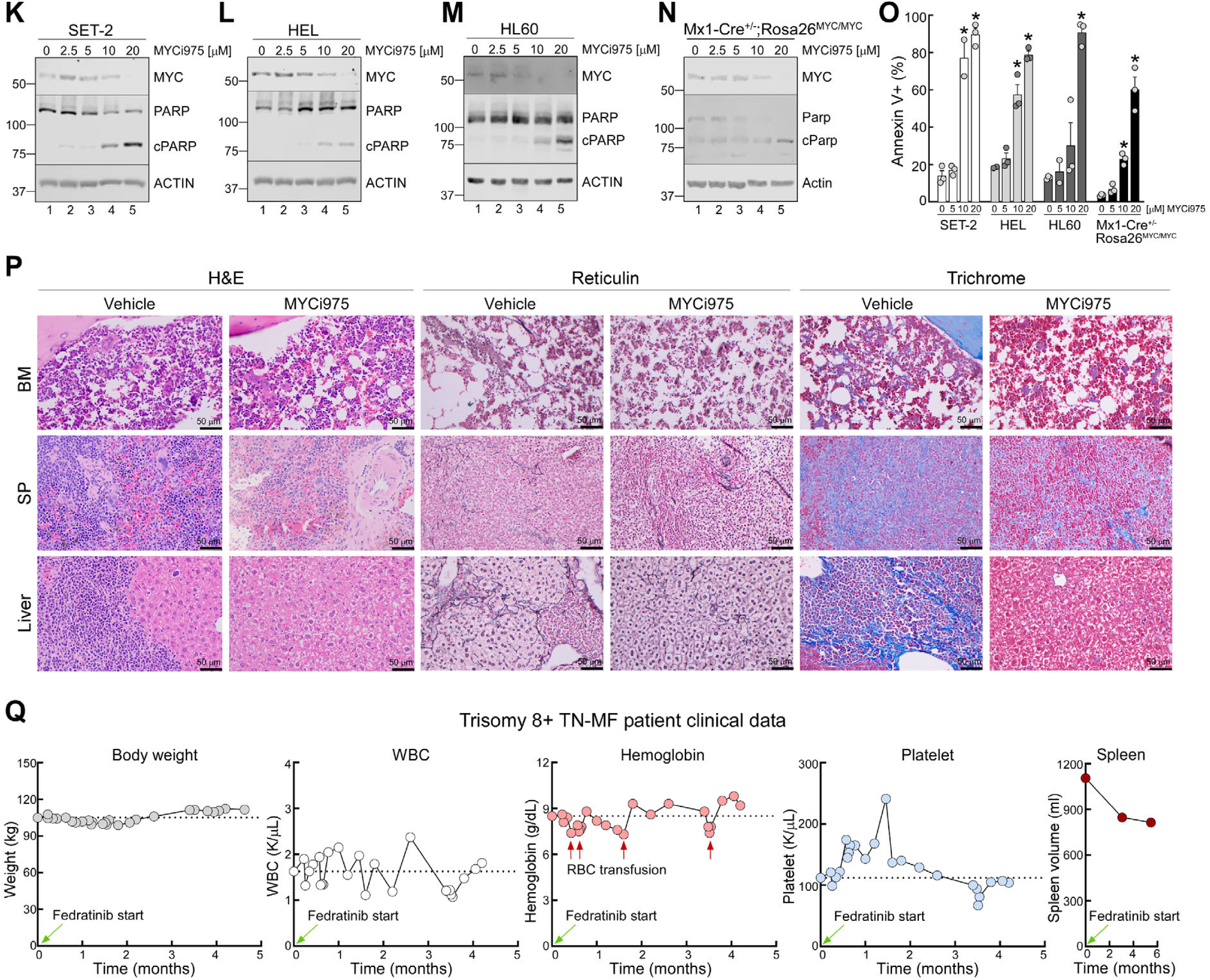
Effects of inhibition of S100a9 or MYC in MYC-driven MF. **A,** Schematic of in vivo Tasquinimod efficacy studies. Mice were randomized to vehicle *vs.* Tasquinimod treatment following pIpC treatment. **B-F,** PB CBC analyses of RBC (B), monocytes (C), neutrophils (D), lymphocytes (E), and platelets (F) in Tasquinimod *vs.* vehicle treated cohorts. Baseline CBC was performed 1 week prior to drug treatment. **G,** Spleen weight at endpoints. Tasquinimod treated mice were sacrificed at ∼32 weeks post-pIpC treatment, which is equivalent to the median OS of vehicle treated Mx1-Cre^+/-^;Rosa26^LSL-MYC/LSL-MYC^ mice. **H,** Percentages of HSCs and MPPs, myeloid progenitors, Gr1^+^/CD11b^+^ mature myeloid cells, macrophages, B-cells, and T-cells in each cohort. **I,** H&E, reticulin, and trichrome stained images of BM, spleen, and liver at week 32 following pIpC injection in each group. **J,** Kaplan-Meier curves of OS. OS was calculated from the first date of Tasquinimod treatment. Demographic profiles and laboratory parameters are described in Table S13. **K-O,** Changes in MYC and cleaved PARP levels (K-N) by immunoblotting and apoptosis (O, % Annexin V^+^ cells) were assessed following 48 hr incubation with MYCi975 at the indicated concentrations. **P,** H&E, reticulin, and trichome stained images of BM, spleen, and liver at MYCi975 study endpoints. **Q,** Serial clinical data in trisomy 8+ TN-MF patient before and during fedratinib treatment. Body weight, WBC, hemoglobin, platelets counts and spleen volume were serially assessed as indicated. Experiments in (H) were performed simultaneously with experiments in Figure 3A-C and Figure S3B. Actin blots in (K-N) were from the same samples run on different gels. Box plots in panels (B-G) represent data from at least 8 mice in each group. Error bars in (H and O) indicate mean ±SEM of at least 3 independent assays. * in (B, D-G and O), *P*<0.05 compared with control group.

### SUPPLEMENTAL TABLES

**Table S1.** Demographics of Moffitt Total Cancer Care MF patients.

| Parameters | *JAK2* mutant patients (n = 363) | *MPL* mutant patients (n = 42) | *CALR* mutant patients (n = 128) | TN patients (n = 51) | All patients (n = 584) |
| --- | --- | --- | --- | --- | --- |
| **Median age at diagnosis, years (range)** | 67 (19-94) | 69 (47-86) | 61 (31-87) | 70 (19-86) | 67 (19-94) |
| **Sex, n (%)** |  |  |  |  |  |
| Male | 209 (58) | 23 (55) | 71 (55) | 29 (57) | 332 (57) |
| Female | 154 (42) | 19 (45) | 57 (45) | 22 (43) | 252 (43) |
| **Type of MF, n (%)** |  |  |  |  |  |
| PMF | 227 (63) | 30 (71) | 79 (62) | 43 (84) | 378 (65) |
| Post-ET | 55 (15) | 12 (29) | 48 (38) | 8 (16) | 122 (21) |
| Post-PV | 81 (22) | 0 (0) | 2 (2) | 0 (0) | 83 (14) |
| **Labs at Diagnosis (range)** |  |  |  |  |  |
| WBC, K/μL | 10.3 (1.4-184.4) | 8.1 (1.9-49.8) | 7.5 (1.5-87.5) | 8.5 (2.0-85.9) | 9.1 (1.4-184.4) |
| Hemoglobin, g/dL | 11.2 (4.9-17.4) | 10.1 (6.4-13.7) | 10.7 (5.6-16.3) | 10.1 (3.4-15.6) | 10.9 (3.4-17.4) |
| Platelet, K/μL | 263.5 (8-2195) | 315 (37-1307) | 381 (18-1459) | 156.5 (8-1238) | 287 (8-2195) |
| Monocyte, K/μL | 0.6 (0.0-31.2) | 0.6 (0.1-9.5) | 0.5 (0.0-3.9) | 0.6 (0.1-5.9) | 0.6 (0.1-5.9) |
| ANC, K/μL | 7.4 (0.4-165.8) | 4.7 (1.1-30.1) | 5.2 (0.6-56.9) | 6.3 (0.2-53.5) | 6.2 (0.2-14.1) |
| ALC, K/μL | 1.5 (0-14.1) | 1.5 (0.5-4.9) | 1.5 (0.4-10.2) | 1.5 (0.4-5.0) | 1.5 (0.0-14.1) |
| **BM assessment** |  |  |  |  |  |
| Blast, % (range) | 0 (0-15) | 0 (0-8) | 0 (0-5) | 0 (0-4) | 0 (0-15) |
| Reticulin fibrosis, grade (range) | 2 (1-3) | 2 (1-3) | 2 (1-3) | 2 (1-3) | 2 (1-3) |
| **Cytogenetics, n (%) \*** | n = 339 | n = 34 | n = 117 | n = 45 | n = 535 |
| +1/dup(1q) | 11 (3) | 0 (0) | 4 (3) | 1 (2) | 16 (3) |
| Del(17p) | 1 (0.4) | 0 (0) | 0 (0) | 1 (2) | 2 (0.4) |
| Del(13q) | 23 (7) | 3 (9) | 15 (13) | 0 (0) | 41 (8) |
| +9 | 26 (8) | 1 (3) | 0 (0) | 0 (0) | 27 (5) |
| Del(20q) | 44 (13) | 3 (9) | 4 (3) | 1 (2) | 52 (10) |
| +8 | 19 (6) | 1 (3) | 1 (1) | 12 (27) | 33 (6) |
| Complex karyotype ** | 28 (8) | 3 (9) | 2 (2) | 3 (7) | 36 (7) |
| **Spleen size, cm (range)** | 16.5 (10-36) | 15 (8-29) | 15.5 (7.6-30) | 14.8 (9.9-23.1) | 16 (7.6-36) |
| **DIPSS, n (%)** | n = 360 | n = 42 | n = 128 | n = 51 | n = 581 |
| Low | 26 (7) | 3 (7) | 24 (19) | 8 (16) | 61 (10) |
| Intermediate-1 | 158 (44) | 15 (36) | 56 (44) | 18 (35) | 247 (43) |
| Intermediate-2 | 130 (36) | 17 (40) | 40 (31) | 16 (31) | 203 (35) |
| High | 46 (13) | 7 (17) | 8 (6) | 9 (18) | 70 (12) |

**Abbreviations:** Myelofibrosis (MF), triple negative (TN), primary MF (PMF), essential thrombocythemia (ET), polycythemia vera (PV), white blood cells (WBC), absolute neutrophil counts (ANC), absolute lymphocyte counts (ALC), bone marrow (BM), Dynamic International Prognostic Scoring System (DIPSS).
\*Cytogenetics were assessed by conventional karyotyping and/or Fluorescence In Situ Hybridization (FISH). Six most common cytogenetic abnormalities are presented.
\*\*Complex karyotype was defined as 3 or more cytogenetic abnormalities.

**Table S2.** Demographic Profile of trisomy 8+ TN-MF patient and healthy donors used in scRNA-seq analysis and PDX studies.

| ID | Age (yr) | Gender | Diagnosis | Conventional Karyotyping | FISH | Somatic mutations | Previous treatments |
| --- | --- | --- | --- | --- | --- | --- | --- |
| PDX | 40 | Male | MF | 47,XY,+8[20] | Trisomy 8+ in 75% | *ASXL1* p.G646Wfs*12, VAF 36.6%<br>*U2AF1* p.S34F, VAF 48.4% | None |
| HD1 | 22 | Female | HD | NA | NA | NA | None |
| HD2 | 20 | Male | HD | NA | NA | NA | None |
| HD3 | 19 | Female | HD | NA | NA | NA | None |

**Abbreviations:** Fluorescence in situ hybridization (FISH), variant allele frequency (VAF), healthy donor (HD), single cell RNA sequencing (scRNA-seq), patient-derived xenograft (PDX), not assessed (NA). See Figure S1I and Figure 7H for the conventional karyotyping and FISH results, respectively, of trisomy 8+ TN-MF patient.

**Table S3.**
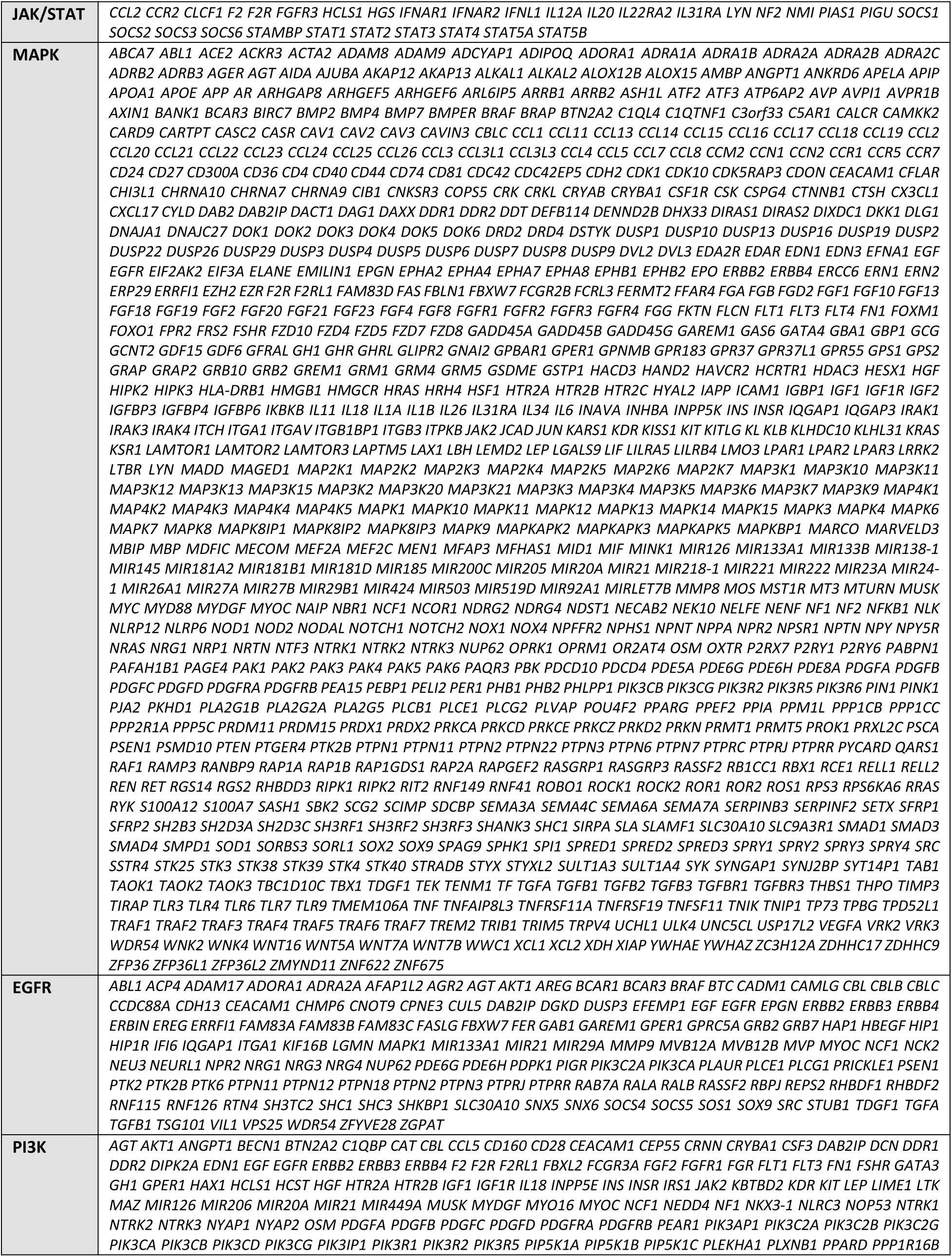
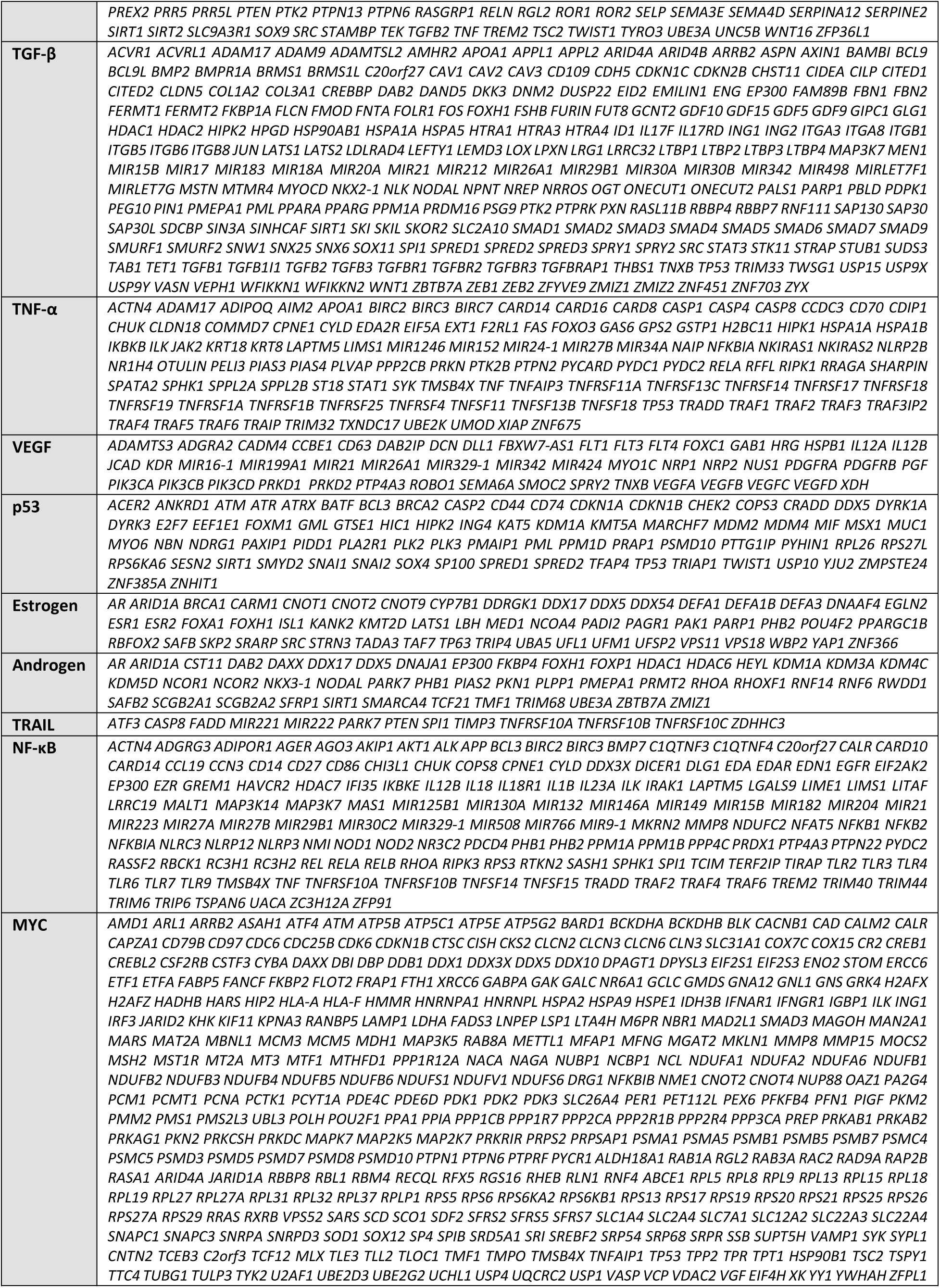
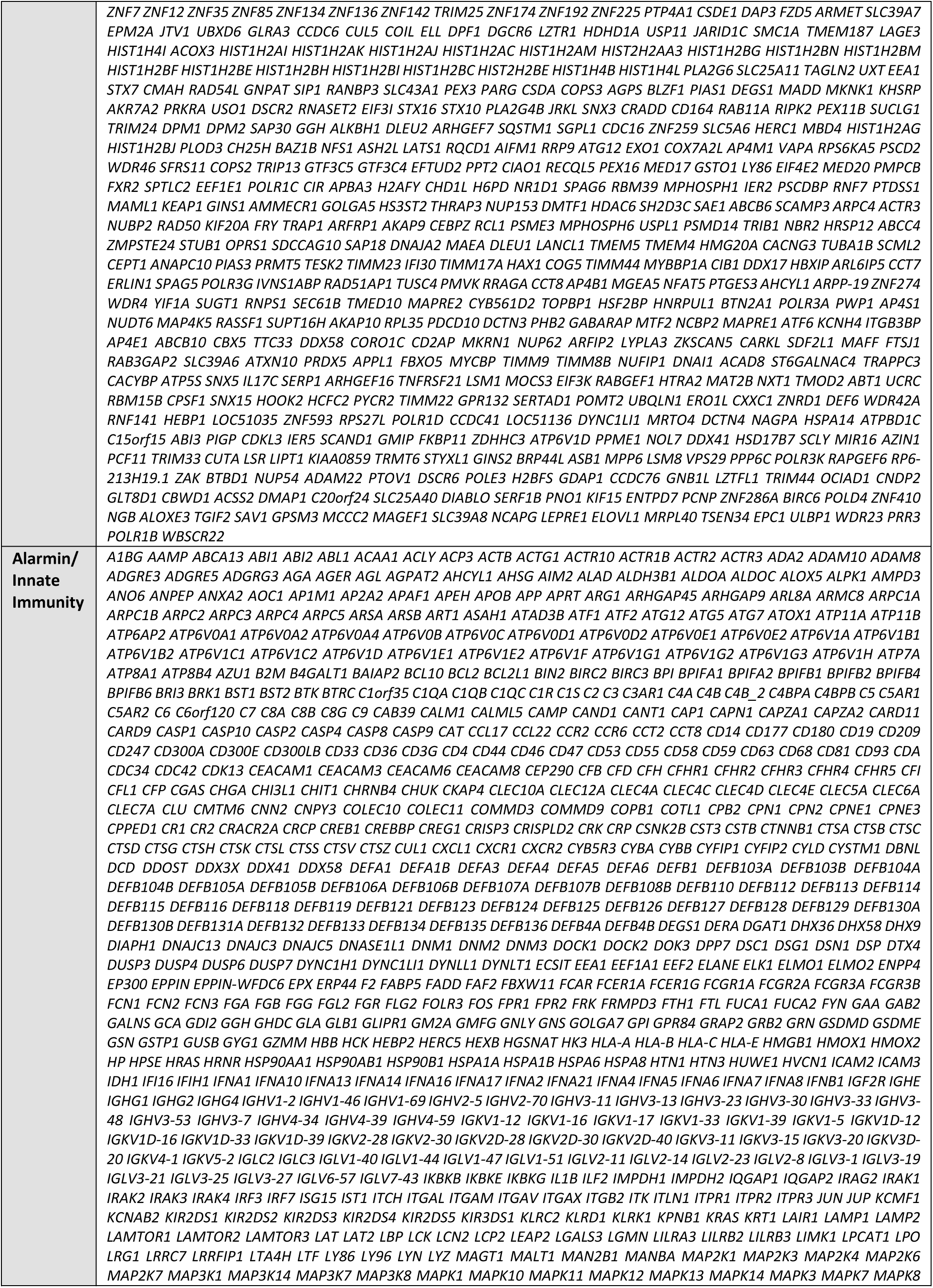
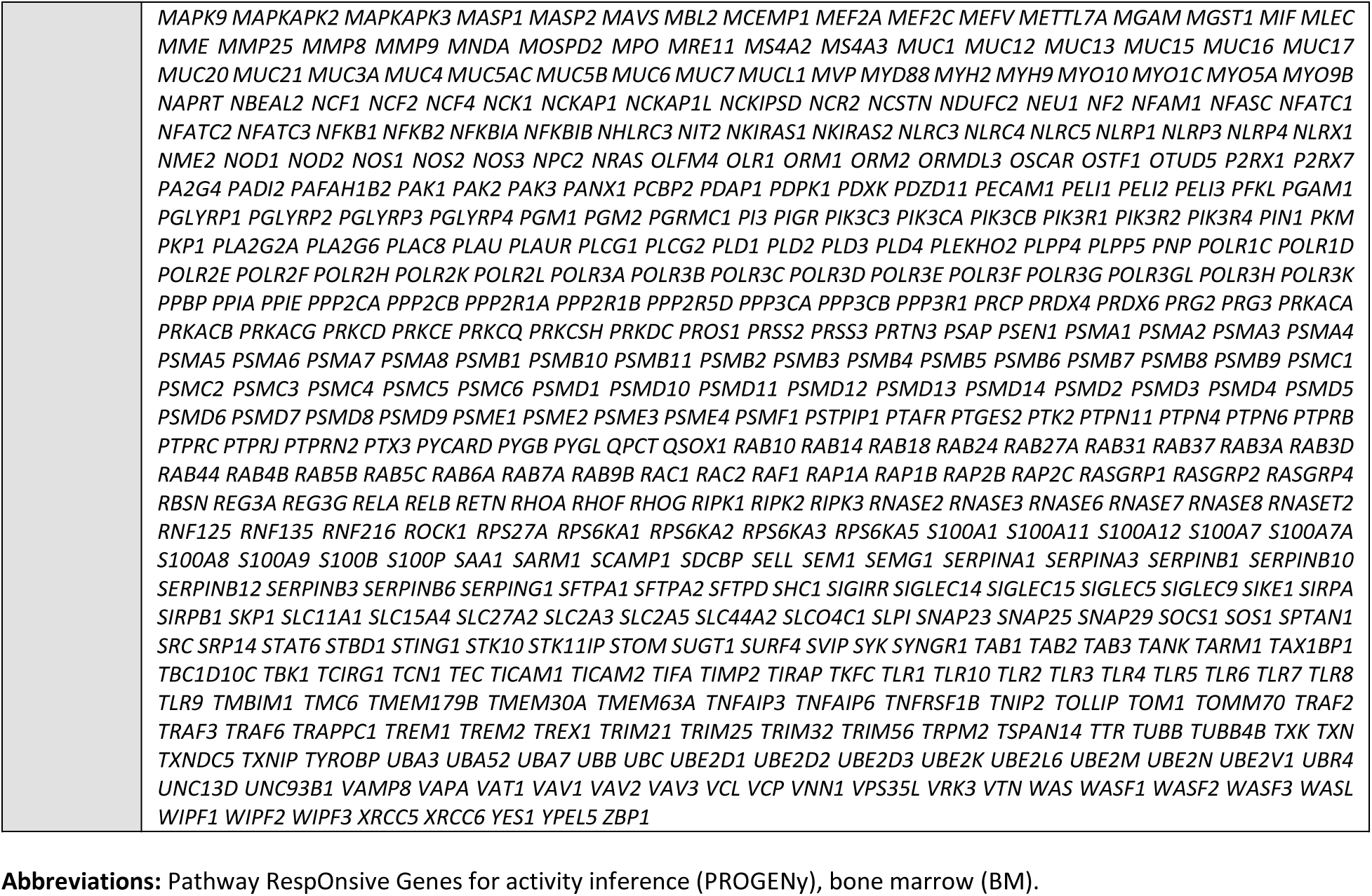
List of genes used for PROGENy analysis of human BM cells.

**Table S4.** Demographics of TN-MF patients.

| Parameters | MYC negative patients<br>(n = 11) | MYC positive patients<br>(n = 12) | All patients (n = 23) |
| --- | --- | --- | --- |
| **Median age at diagnosis, years (range)** | 62 (19-86) | 65 (53-77) | 63 (19-86) |
| **Sex, n (%)** |  |  |  |
| Male | 7 (64) | 5 (42) | 12 (52) |
| Female | 4 (36) | 7 (58) | 11 (48) |
| **Type of MF, n (%)** |  |  |  |
| PMF | 9 (82) | 10 (83) | 19 (83) |
| Post-ET | 2 (18) | 2 (17) | 4 (17) |
| Post-PV | 0 (0) | 0 (0) | 0 (0) |
| **Labs at Diagnosis (range)** |  |  |  |
| WBC, K/μL | 8.5 (2.3-82.8) | 6.0 (2.2-51.7) | 6.6 (2.2-82.8) |
| Hemoglobin, g/dL | 10.9 (7.7-15.6) | 9.6 (3.4-12.8) | 10.5 (3.4-15.6) |
| Platelet, K/μL | 172 (48-967) | 114 (8-378) | 151 (8-967) |
| Monocyte, K/μL | 0.3 (0.1-5.4) | 0.4 (0.1-1.2) | 0.3 (0.1-5.4) |
| **BM assessment** |  |  |  |
| Blast, % (range) | 0 (0-4) | 0 (0-3) | 0 (0-4) |
| Reticulin fibrosis, grade (mean) | 2 | 2.4 | 2.2 |
| **Cytogenetics, n (%) \*** |  |  |  |
| +8 | 1 (9) | 5 (42) | 6 (26) |
| +21 | 1 (9) | 0 (0) | 1 (4) |
| +1/dup(1q) | 0 (0) | 1 (8) | 1 (4) |
| Tetraploidy | 0 (0) | 1 (8) | 1 (4) |
| Complex karyotype ** | 1 (9) | 0 (0) | 1 (4) |
| **DIPSS, n (%)** |  |  |  |
| Low | 4 (36) | 0 (0) | 4 (17) |
| Intermediate-1 | 4 (36) | 6 (50) | 10 (43) |
| Intermediate-2 | 2 (18) | 5 (42) | 7 (30) |
| High | 1 (9) | 1 (8) | 2 (9) |

**Abbreviations:** Myelofibrosis (MF), triple negative MF (TN-MF), primary MF (PMF), essential thrombocythemia (ET), polycythemia vera (PV), white blood cells (WBC), absolute neutrophil counts (ANC), absolute lymphocyte counts (ALC), bone marrow (BM), Dynamic International Prognostic Scoring System (DIPSS).
\*Cytogenetics were assessed by conventional karyotyping and/or Fluorescence In Situ Hybridization (FISH). \*\*Complex karyotype was defined as 3 or more cytogenetic abnormalities.

**Table S5.** Clinical parameters of Mx1-Cre^+/-^;Rosa26^LSL-MYC/LSL-MYC^ studies.

| Parameters | Rosa26<sup>+/+</sup> | Rosa26<sup>LSL-MYC/+</sup> | Rosa26<sup>LSL-MYC/LSL-MYC</sup> |
| --- | --- | --- | --- |
| Number of mice | 38 | 44 | 54 |
| Age at plpC treatment, weeks | 6-11 | 6-11 | 6-11 |
| Female, n (%) | 17 (45) | 38 (86) | 24 (44) |
| Median baseline body weight, mg (range) | 19.0 (13.0-26.9) | 18.7 (14.4-27.2) | 19.1 (15.4-26.6) |
| **CBC at baseline (range)\*** |  |  |  |
| WBC, K/μL | 8.72 (4.15-16.3) | 11.13 (4.16-16.85) | 9.67 (2.86-19.27) |
| RBC, M/μL | 10.15 (8.92-10.91) | 10.02 (8.68-11.67) | 10.24 (8.40-11.57) |
| Platelet, K/μL | 781.5 (15.4-1175) | 790 (14.4-1207) | 785 (15-1011) |
| Neutrophil, % | 9.45 (4.1-35.9) | 9.9 (2.5-49.4) | 9.2 (3.5-45.2) |
| Monocyte, % | 1.05 (0.6-2.4) | 1.2 (0.4-3.6) | 1.85 (0.2-3.9) |
| Lymphocyte, % | 87.85 (61.3-93.8) | 87 (47.5-95.3) | 87.9 (51.6-94.2) |
| **CBC at week 25-36 (range)** |  |  |  |
| WBC, K/μL | 7.28 (2.12-18.9) | 7.62 (0.13-18.7) | 9.74 (1.82-20.32) |
| RBC, M/μL | 10.97 (9.05-12.63) | 10.29 (7.9-12.14) | 9.27 (0.79-11.29) |
| Platelet, K/μL | 1076.5 (17.1-1591) | 1014 (14.8-1603) | 994 (1-2013) |
| Neutrophil, % | 12.1 (6.2-50.5) | 11.7 (2.3-53.8) | 11.45 (1.6-33.1) |
| Monocyte, % | 2.4 (0.6-8.3) | 2.9 (0-6.1) | 5.05 (1.6-15.7) |
| Lymphocyte, % | 78.5 (46.6-90.2) | 78.6 (42.8-88.1) | 76.9 (9.2-89.1) |
| **Median spleen weight, mg (range)\*\*** |  |  |  |
| At week 15-44 | 100 (80-130) | 160 (90-940) | 325 (130-720) |
| At week 45-62 | 110 (30-140) | 205 (130-780) | 1300 (1300-1300) |
| **Median OS, days\*\*\*** | NR | 385 | 258 |

**Abbreviations:** Polyinosinic-polycytidylic acid (plpC), complete blood counts (CBC), white blood cells (WBC), red blood cells (RBC), overall survival (OS), and not reached (NR).
\*Baseline CBC values were determined 1 week prior to plpC injection.
\*\*Spleen weight was measured at the time of death or at pre-defined endpoints. See Method.
\*\*\*Overall survival was calculated from the first date of plpC injection.

**Table S6.**
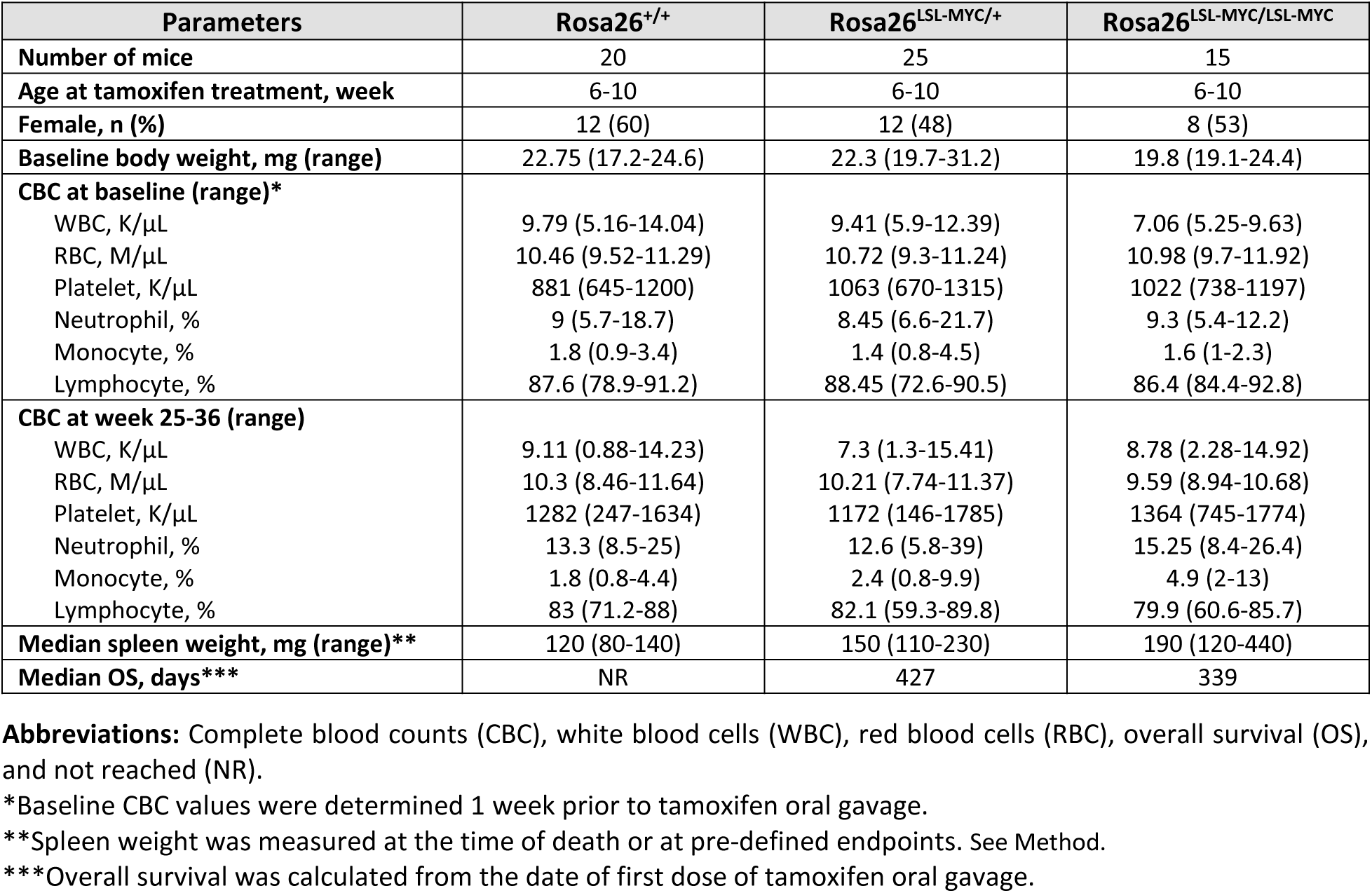
Clinical parameters of Scl-CreERT^+/-^;Rosa26^LSL-MYC/LSL-MYC^ *in vivo* studies.

**Table S7.**
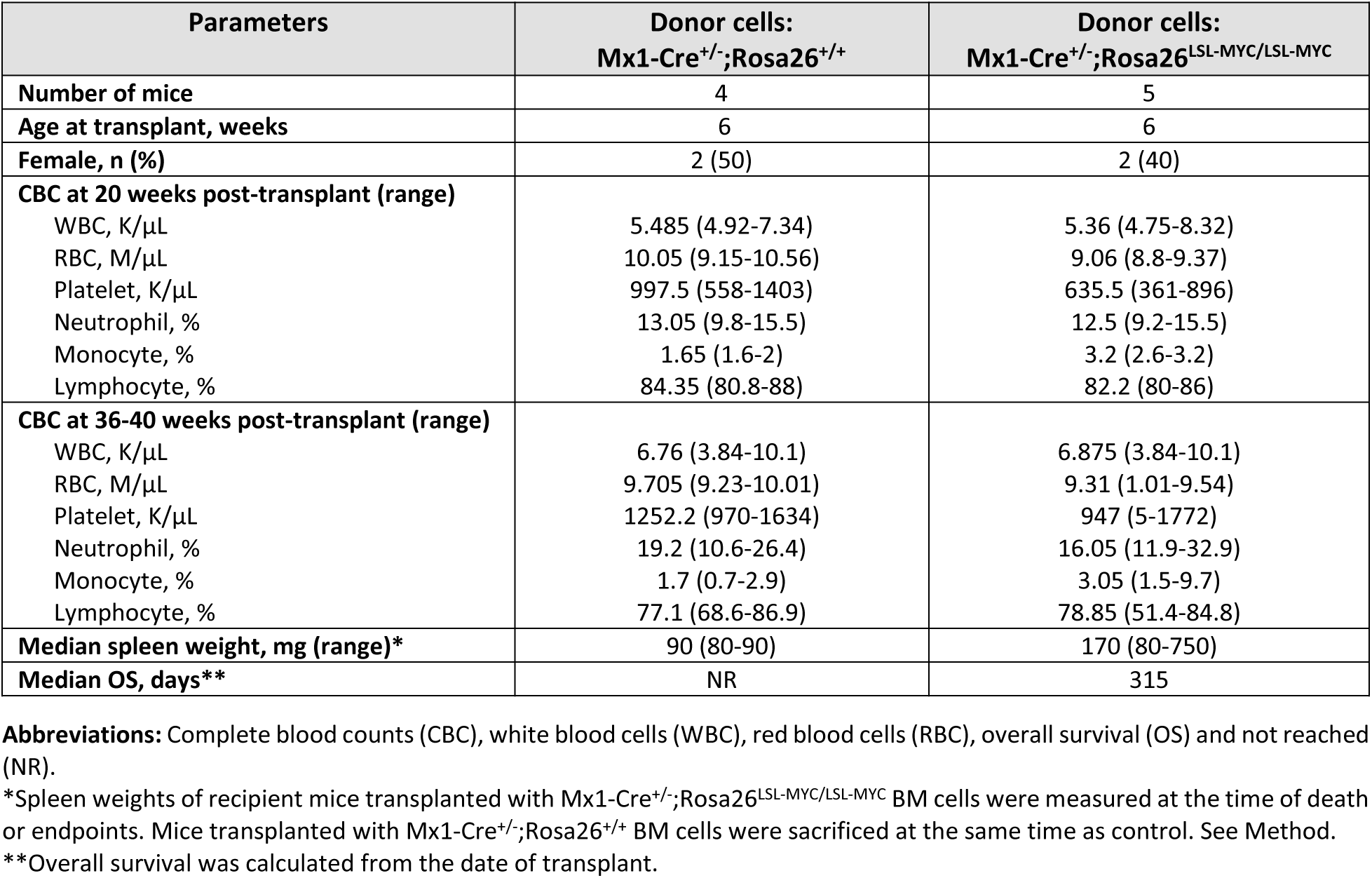
Clinical parameters in competitive transplant studies.

**Table S8.** Comparison of gene expression profiles of Mx1-Cre^+/-^;Rosa26^LSL-MYC/LSL-MYC^ *vs.* Mx1-Cre^+/-^;Rosa26^+/+^ mouse. See “Table S8” file.

**Table S9.**
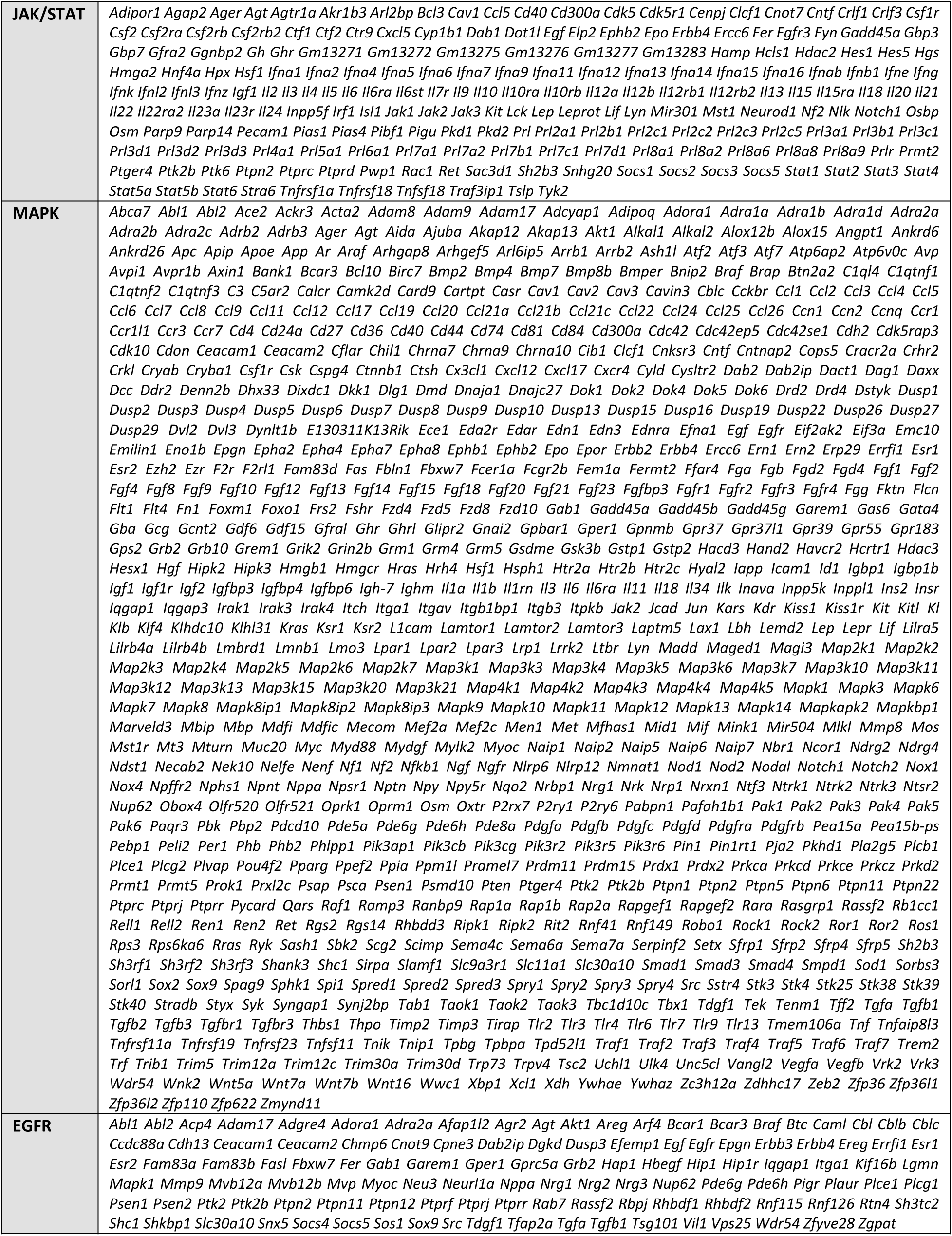
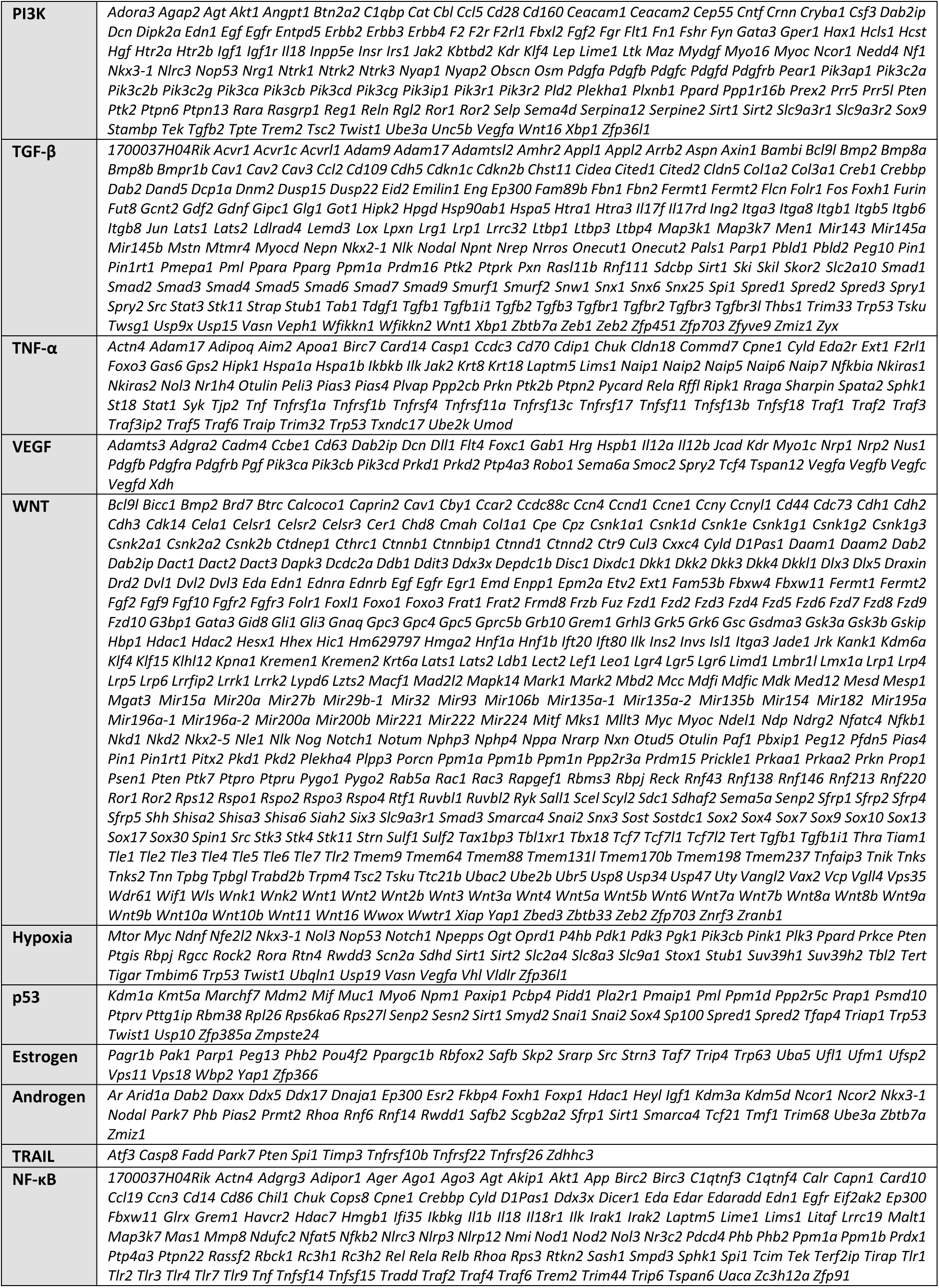
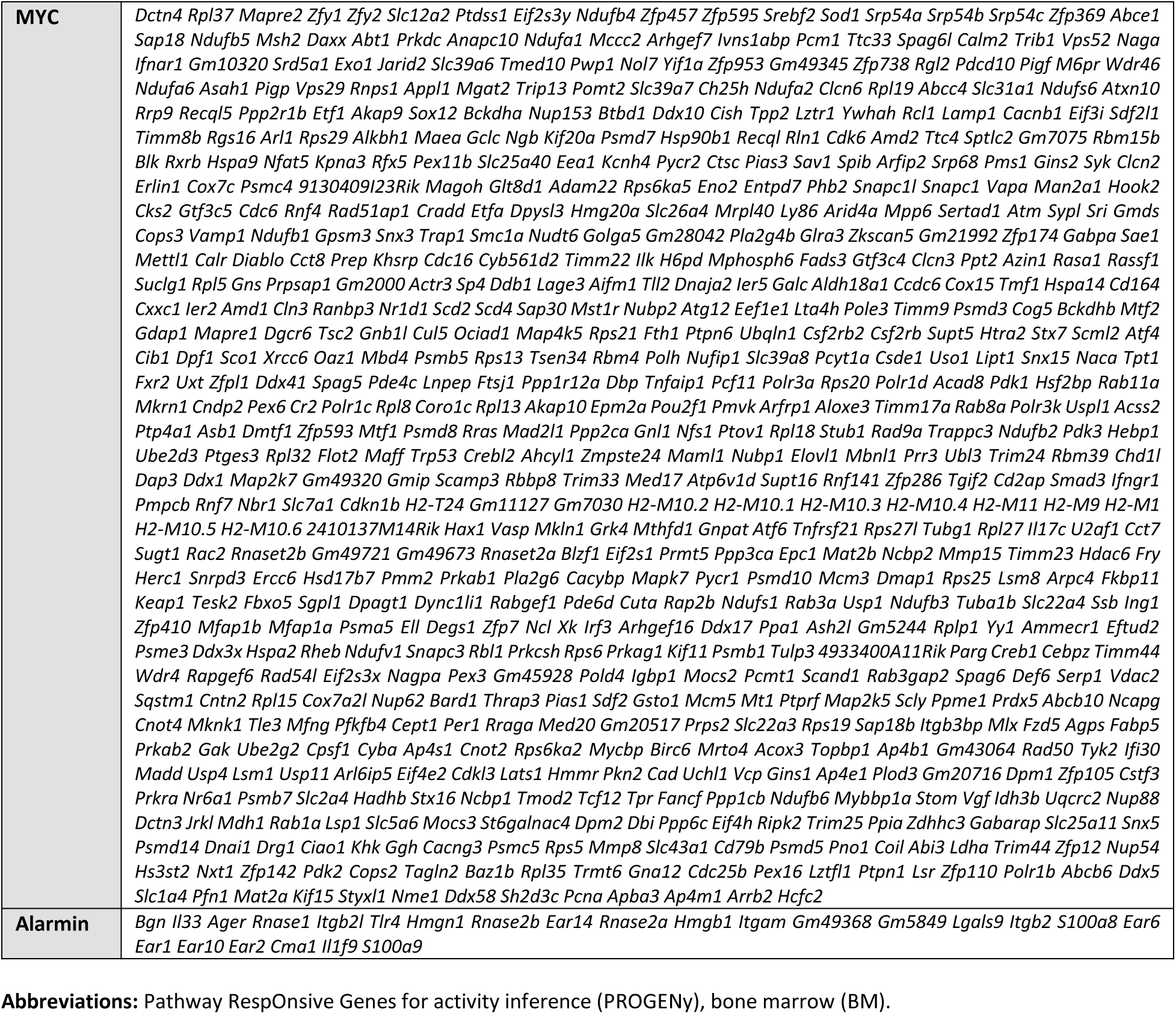
List of genes used for PROGENy analysis of mouse BM cells.

**Table S10.** Cell-cell interaction analyses. See “Table S10” file.

**Table S11.**
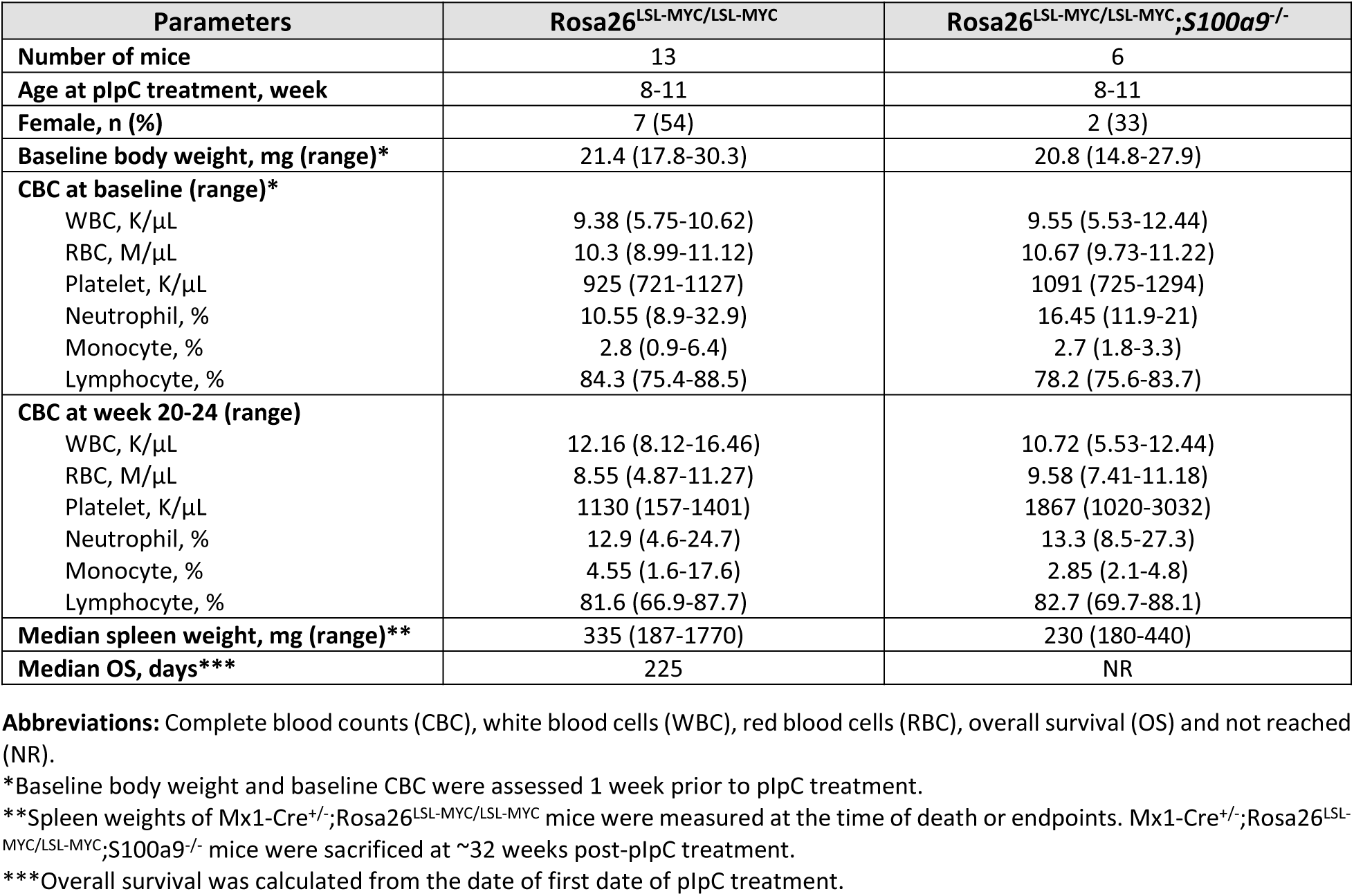
Clinical parameters of Mx1-Cre^+/-^;Rosa26^LSL-MYC/LSL-MYC^;*S100a9*^-/-^ studies.

**Table S12.** Clinical parameters of S100a9 transgenic mouse studies.

| Parameters | Wild type control | S100a9Tg |
| --- | --- | --- |
| **Number of mice** | 20 | 56 |
| **Female, n (%)** | 10 (50) | 26 (46) |
| **CBC at month 3 (range)** | N=5 | N=6 |
| WBC, K/μL | 8.46 (6.82-11.46) | 10.1 (6.42-14.57) |
| RBC, M/μL | 11.25 (10.84-11.55) | 10.63 (8.8-11.36) |
| Platelet, K/μL | 963 (901-1149) | 1074.5 (216-1546) |
| Neutrophil, % | 9.4 (7.1-32.6) | 11.2 (8.6-14.4) |
| Monocyte, % | 1.3 (1.0-2.3) | 1.85 (1.4-3.1) |
| Lymphocyte, % | 87.2 (64.8-90.8) | 85.5 (82.2-87.5) |
| **CBC at month 12 (range)** | N=4 | N=34 |
| WBC, K/μL | 10.36 (6.41-11.04) | 11.9 (4.47-164.59) |
| RBC, M/μL | 9.89 (9.45-10.0) | 10.00 (7.5-10.99) |
| Platelet, K/μL | 1571.5 (1254-1625) | 1357.5 (161-2383) |
| Neutrophil, % | 12.45 (11.5-14.8) | 15 (9.4-47.2) |
| Monocyte, % | 3.1 (1.7-4.3) | 4.25 (2.2-42.2) |
| Lymphocyte, % | 83.05 (81.5-85.7) | 80.8 (4.6-86.7) |
| **Median spleen weight, mg (range)\*** | 135 (90-230) | 580 (80-3950) |
| **Median OS, days\*\*\*** | NR | 371 |

**Abbreviations:** S100a9 transgenic mouse (S100a9Tg), complete blood counts (CBC), white blood cells (WBC), red blood cells (RBC), and overall survival (OS).
\*Spleen weights of S100a9Tg mice were measured at the time of death or at pre-defined endpoints. Age-matched control mice were sacrificed at the same time.
\*\*Overall survival was calculated from the date of birth

**Table S13.** Clinical parameters of Tasquinimod efficacy studies in MYC-driven MF.

| Parameters | Regular water | Tasquinimod water |
| --- | --- | --- |
| Number of mice | 13 | 8 |
| Age at plpC treatment, week | 8-11 | 8-11 |
| Female, n (%) | 7 (54) | 5 (62.5) |
| Baseline body weight, mg (range)* | 21.4 (17.8-30.3) | 20.9 (16.3-24) |
| **CBC at baseline (range)\*** |  |  |
| WBC, K/μL | 9.38 (5.75-10.62) | 9.81 (6.46-13.45) |
| RBC, M/μL | 10.3 (8.99-11.12) | 10.41 (10.02-10.91) |
| Platelet, K/μL | 925 (721-1127) | 1030.5 (420-1260) |
| Neutrophil, % | 10.55 (8.9-32.9) | 7.4 (6.2-33.9) |
| Monocyte, % | 2.8 (0.9-6.4) | 2.1 (1.1-3) |
| Lymphocyte, % | 84.3 (75.4-88.5) | 88.05 (62.6-90.4) |
| **CBC at week 20 (range)** |  |  |
| WBC, K/μL | 12.16 (8.12-16.46) | 11.07 (8.93-20.08) |
| RBC, M/μL | 8.55 (4.87-11.27) | 7.68 (4.69-9.02) |
| Platelet, K/μL | 1130 (157-1401) | 1615 (998-3084) |
| Neutrophil, % | 12.9 (4.6-24.7) | 27.7 (11.2-36.5) |
| Monocyte, % | 4.55 (1.6-17.6) | 5.4 (2.3-10.6) |
| Lymphocyte, % | 81.6 (66.9-87.7) | 64.85 (52.4-86.2) |
| Median spleen weight, mg (range)** | 370 (212-1770) | 240 (180-640) |
| Median OS, days*** | 208 | 251 |

**Abbreviations:** Complete blood counts (CBC), white blood cells (WBC), red blood cells (RBC), overall survival (OS) and not reached (NR).
\*Baseline body weight and baseline CBC were assessed 1 week prior to plpC treatment.
\*\*Spleen weights of control mice (vehicle treatment) were measured at the time of death or endpoints. Mice treated with Tasquinimod were sacrificed at ~32 weeks post-plpC treatment.
\*\*\*Overall survival was calculated from the date of first date of Tasquinimod treatment.

**Table S14.** Clinical parameters of MYCi975 efficacy studies in MYC-driven MF.

| Parameters | Vehicle Treatment | MYCi975 Treatment |
| --- | --- | --- |
| Number of mice | 6 | 6 |
| Age at transplant, week | 12-15 | 12-15 |
| Female, n (%) | 6 (100) | 6 (100) |
| Baseline body weight, mg (range) | 21.2 (17.2-23.8) | 20.35 (19.5-23.4) |
| **CBC at baseline (range)\*** |  |  |
| WBC, K/μL | 6.952 (5.83-8.49) | 8.35 (6.27-10.33) |
| RBC, M/μL | 10.965 (10.5-11.34) | 10.945 (10.6-11.2) |
| Platelet, K/μL | 1041 (338-1210) | 851 (479-1062) |
| Neutrophil, % | 5.15 (3.7-8) | 6.8 (4.3-7.7) |
| Monocyte, % | 1.95 (0.9-2.6) | 1.5 (1.3-2.9) |
| Lymphocyte, % | 90.4 (86.9-94.4) | 89.4 (88-92.9) |
| **CBC at 32-36 weeks post-transplant (range)** |  |  |
| WBC, K/μL | 7.345 (3.18-12.08) | 9.6 (7.2-13.86) |
| RBC, M/μL | 9.84 (8.66-10.12) | 9.535 (8.77-10.08) |
| Platelet, K/μL | 817 (440-884) | 1153.5 (424-1801) |
| Neutrophil, % | 9.85 (5-16.3) | 10.2 (7.3-13.8) |
| Monocyte, % | 6.25 (3.6-8.6) | 2.45 (1.5-3) |
| Lymphocyte, % | 81.1 (76.1-89) | 84.25 (82.6-88.5) |
| **Median spleen weight, mg (range)\*\*** | 90 (80-90) | 170 (80-750) |
| **Median OS, days\*\*\*** | 252 | NR |

**Abbreviations:** Complete blood counts (CBC), white blood cells (WBC), red blood cells (RBC), overall survival (OS) and not reached (NR).
\*Baseline CBC values were determined 1 week prior to transplant.
\*\*Spleen weights of vehicle treatment group were measured at the time of death or endpoints. MYCi975 treated mice were sacrificed at the same time for paired analysis.
\*\*\*Overall survival was calculated from the date of transplant.

**Table S15:**
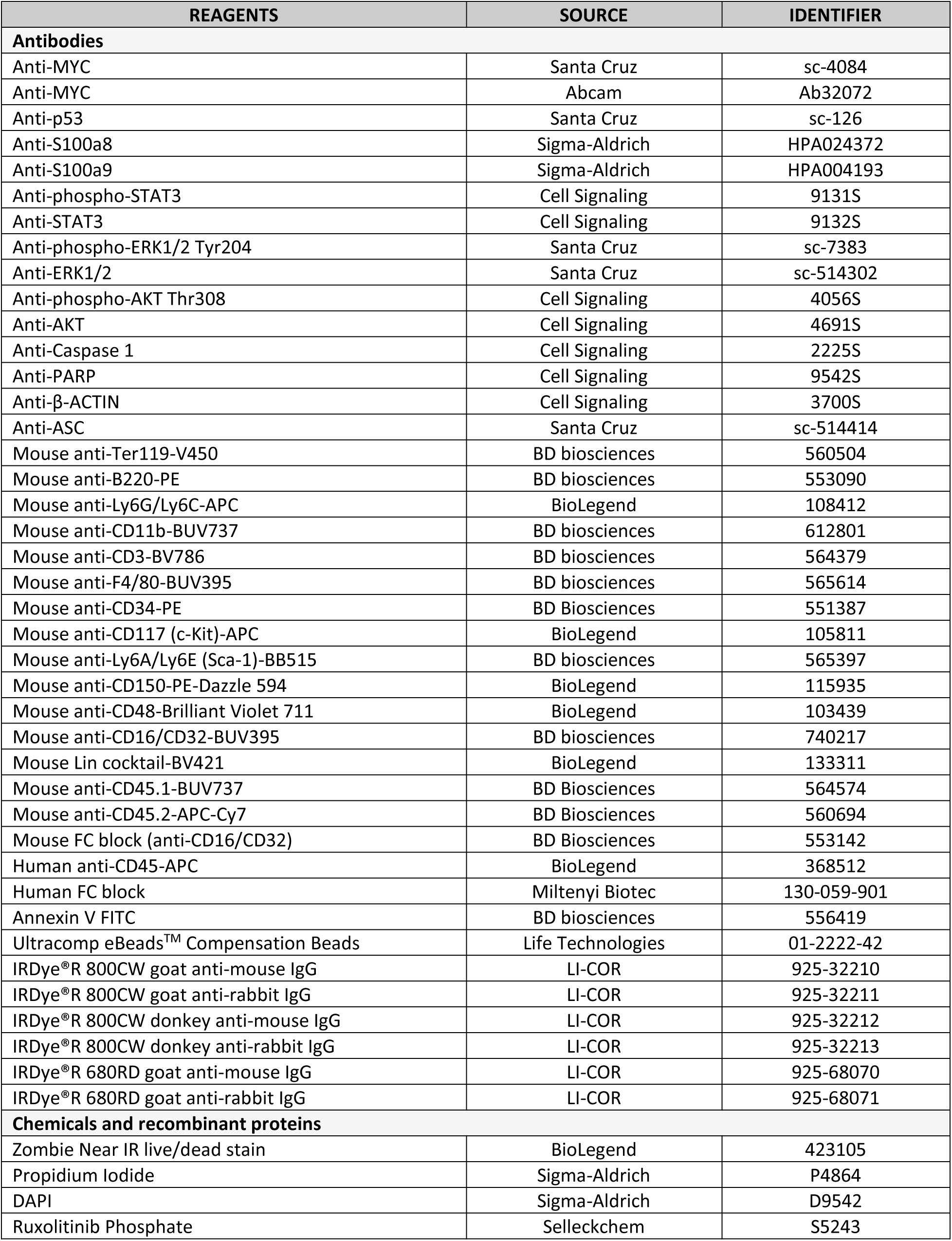
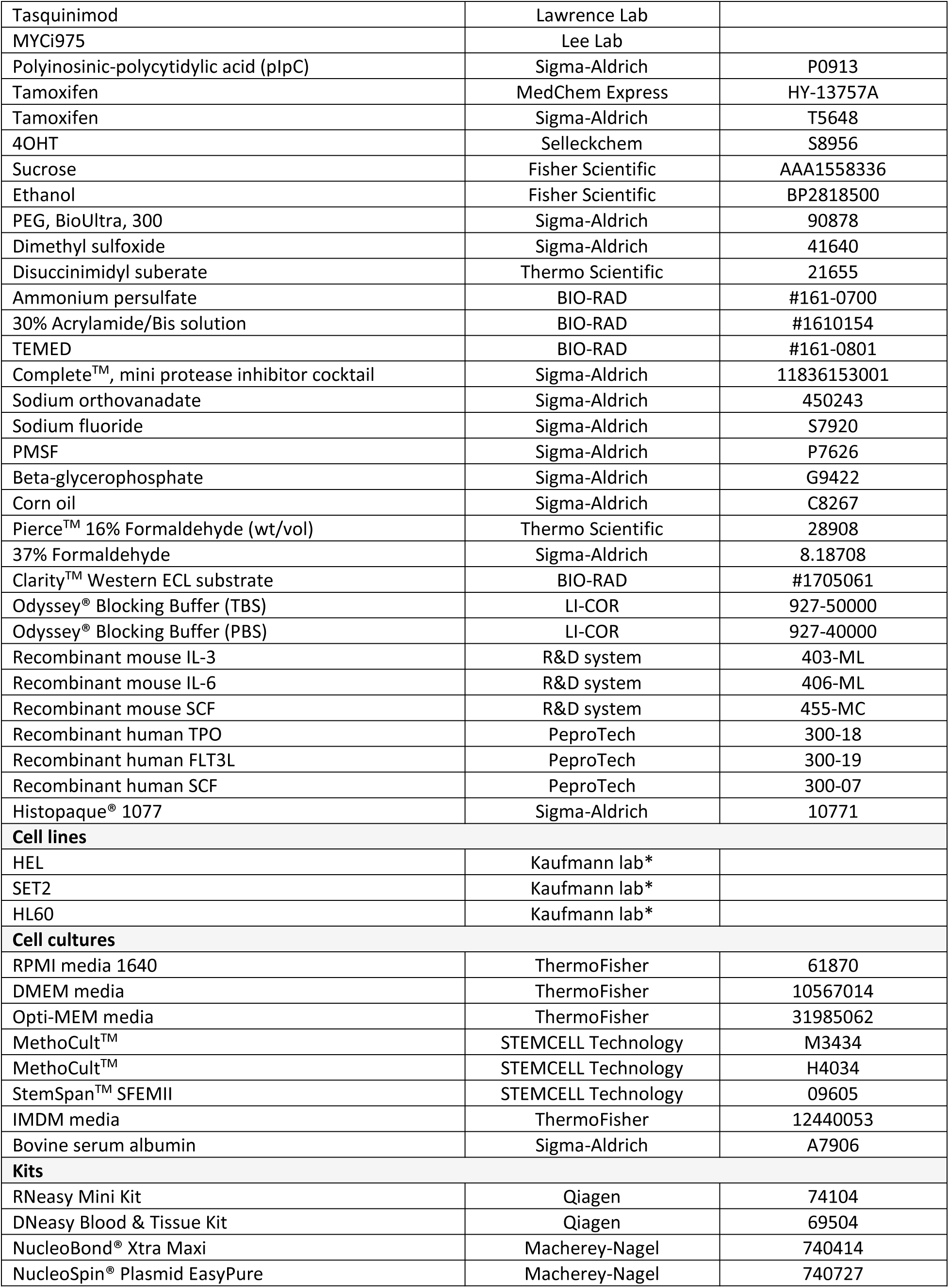
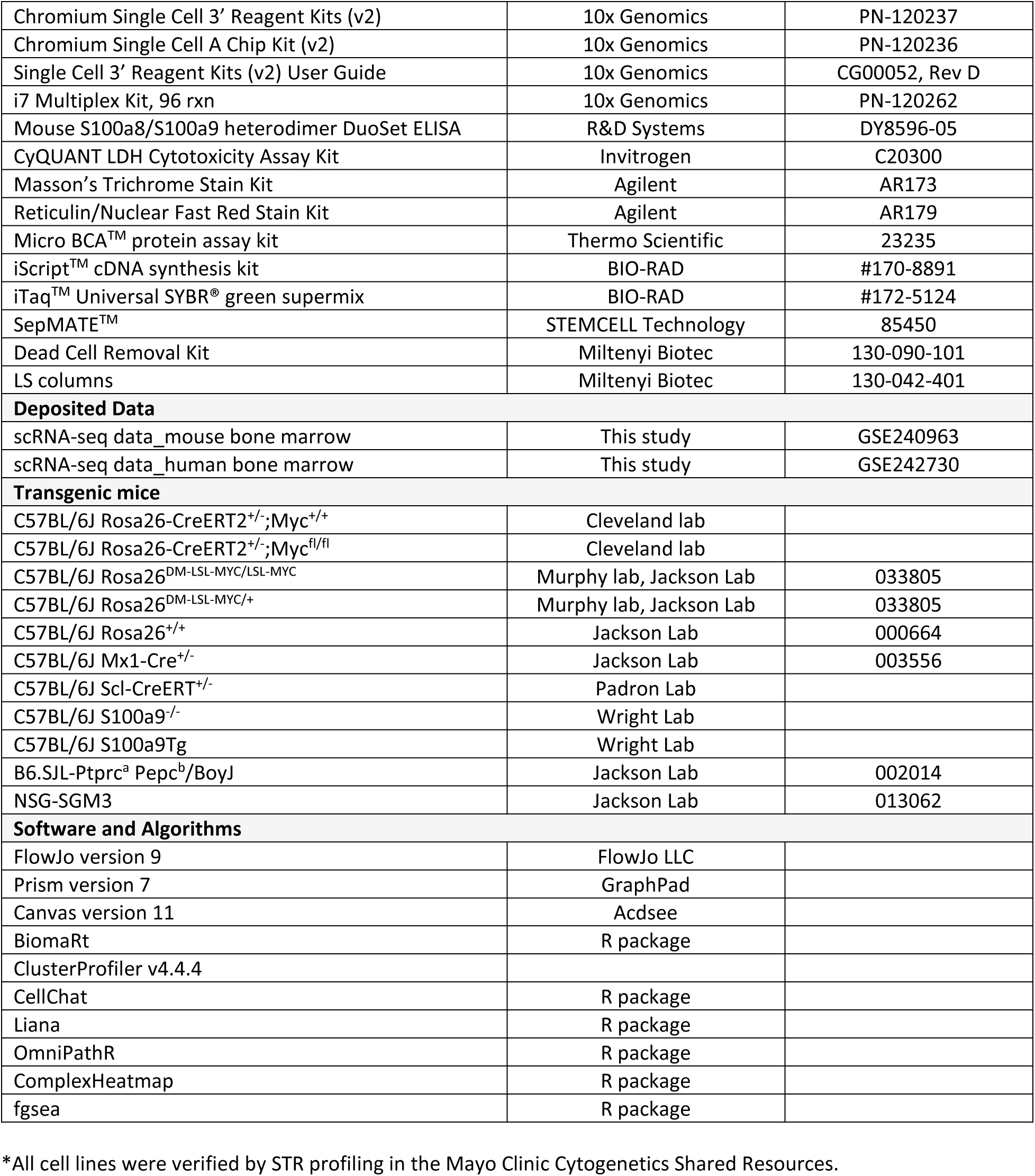
Key Resources Table.

**Table S16:**
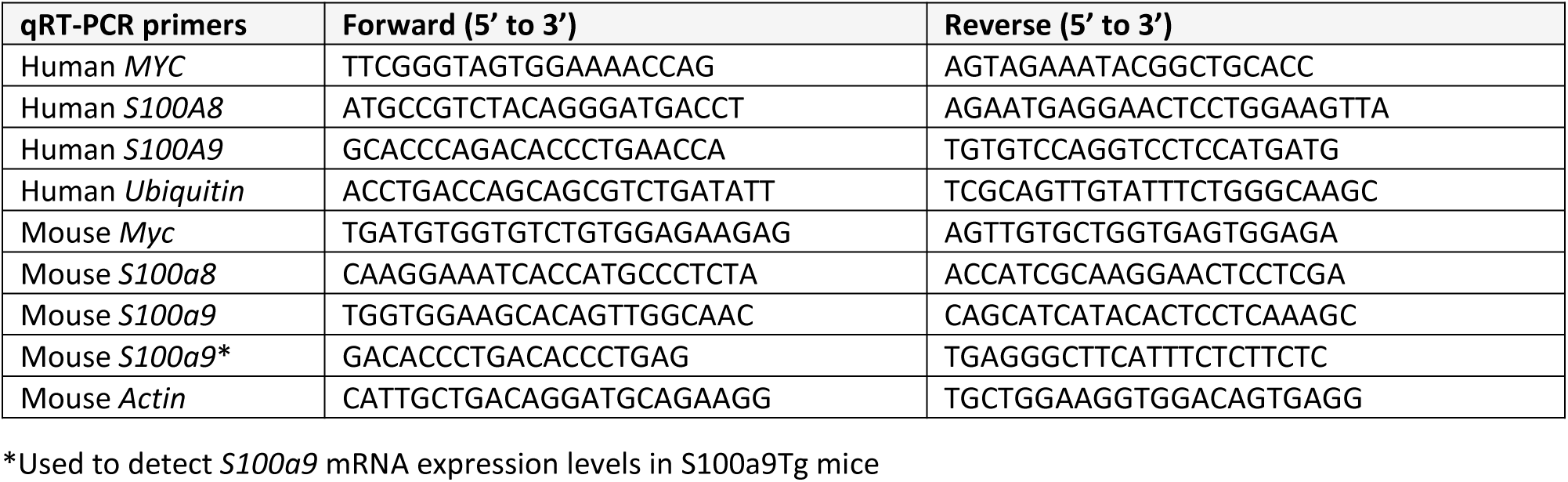
Sequence Information.

## REFERENCES

1. Verstovsek, S., Yu, J., Scherber, R.M., Verma, S., Dieyi, C., Chen, C.-C., and Parasuraman, S. (2022). Changes in the incidence and overall survival of patients with myeloproliferative neoplasms between 2002 and 2016 in the United States. Leukemia & Lymphoma 63, 694–702. 10.1080/10428194.2021.1992756.

2. Spivak, J.L. (2017). Myeloproliferative Neoplasms. N Engl J Med 376, 2168–2181. 10.1056/NEJMra1406186.

3. Tefferi, A. (2021). Primary myelofibrosis: 2021 update on diagnosis, risk-stratification and management. Am J Hematol 96, 145–162. 10.1002/ajh.26050.

4. Klampfl, T., Gisslinger, H., Harutyunyan, A.S., Nivarthi, H., Rumi, E., Milosevic, J.D., Them, N.C.C., Berg, T., Gisslinger, B., Pietra, D., et al. (2013). Somatic Mutations of Calreticulin in Myeloproliferative Neoplasms. N Engl J Med 369, 2379–2390. 10.1056/NEJMoa1311347.

5. Nangalia, J., Massie, C.E., Baxter, E.J., Nice, F.L., Gundem, G., Wedge, D.C., Avezov, E., Li, J., Kollmann, K., Kent, D.G., et al. (2013). Somatic CALR Mutations in Myeloproliferative Neoplasms with Nonmutated JAK2. N Engl J Med 369, 2391–2405. 10.1056/NEJMoa1312542.

6. Cabagnols, X., Favale, F., Pasquier, F., Messaoudi, K., Defour, J.P., Ianotto, J.C., Marzac, C., Le Couédic, J.P., Droin, N., Chachoua, I., et al. (2016). Presence of atypical thrombopoietin receptor (MPL) mutations in triple-negative essential thrombocythemia patients. Blood 127, 333–342. 10.1182/blood-2015-07-661983 %J Blood.

7. Milosevic Feenstra, J.D., Nivarthi, H., Gisslinger, H., Leroy, E., Rumi, E., Chachoua, I., Bagienski, K., Kubesova, B., Pietra, D., Gisslinger, B., et al. (2016). Whole-exome sequencing identifies novel MPL and JAK2 mutations in triple-negative myeloproliferative neoplasms. Blood 127, 325–332. 10.1182/blood-2015-07-661835 %J Blood.

8. Pikman, Y., Lee, B.H., Mercher, T., McDowell, E., Ebert, B.L., Gozo, M., Cuker, A., Wernig, G., Moore, S., Galinsky, I., et al. (2006). MPLW515L Is a Novel Somatic Activating Mutation in Myelofibrosis with Myeloid Metaplasia. PLOS Medicine 3, e270. 10.1371/journal.pmed.0030270.

9. Grinfeld, J., Nangalia, J., Baxter, E.J., Wedge, D.C., Angelopoulos, N., Cantrill, R., Godfrey, A.L., Papaemmanuil, E., Gundem, G., MacLean, C., et al. (2018). Classification and Personalized Prognosis in Myeloproliferative Neoplasms. N Engl J Med 379, 1416–1430. 10.1056/NEJMoa1716614.

10. Kleppe, M., Kwak, M., Koppikar, P., Riester, M., Keller, M., Bastian, L., Hricik, T., Bhagwat, N., McKenney, A.S., Papalexi, E., et al. (2015). JAK–STAT Pathway Activation in Malignant and Nonmalignant Cells Contributes to MPN Pathogenesis and Therapeutic Response. Cancer Discov 5, 316–331. 10.1158/2159-8290.CD-14-0736 %J Cancer Discovery.

11. Tefferi, A., Vaidya, R., Caramazza, D., Finke, C., Lasho, T., and Pardanani, A. (2011). Circulating Interleukin (IL)-8, IL-2R, IL-12, and IL-15 Levels Are Independently Prognostic in Primary Myelofibrosis: A Comprehensive Cytokine Profiling Study. J Clin Oncol 29, 1356–1363. 10.1200/jco.2010.32.9490.

12. Verstovsek, S., Mesa, R.A., Gotlib, J., Levy, R.S., Gupta, V., DiPersio, J.F., Catalano, J.V., Deininger, M., Miller, C., Silver, R.T., et al. (2012). A Double-Blind, Placebo-Controlled Trial of Ruxolitinib for Myelofibrosis. N Engl J Med 366, 799–807. 10.1056/NEJMoa1110557.

13. Harrison, C., Kiladjian, J.-J., Al-Ali, H.K., Gisslinger, H., Waltzman, R., Stalbovskaya, V., McQuitty, M., Hunter, D.S., Levy, R., Knoops, L., et al. (2012). JAK Inhibition with Ruxolitinib versus Best Available Therapy for Myelofibrosis. N Engl J Med 366, 787–798. 10.1056/NEJMoa1110556.

14. Pardanani, A., Harrison, C., Cortes, J.E., Cervantes, F., Mesa, R.A., Milligan, D., Masszi, T., Mishchenko, E., Jourdan, E., Vannucchi, A.M., et al. (2015). Safety and Efficacy of Fedratinib in Patients With Primary or Secondary Myelofibrosis: A Randomized Clinical Trial. JAMA Oncology 1, 643–651. 10.1001/jamaoncol.2015.1590 %J JAMA Oncology.

15. Mascarenhas, J., Hoffman, R., Talpaz, M., Gerds, A.T., Stein, B., Gupta, V., Szoke, A., Drummond, M., Pristupa, A., Granston, T., et al. (2018). Pacritinib vs Best Available Therapy, Including Ruxolitinib, in Patients With Myelofibrosis: A Randomized Clinical Trial. JAMA Oncology 4, 652–659. 10.1001/jamaoncol.2017.5818 %J JAMA Oncology.

16. Aguirre, L.E.E., Jain, A.G., Ball, S., Al Ali, N., Tinsley-Vance, S.M., Sallman, D.A., Sweet, K., Lancet, J.E., Padron, E., Kuykendall, A.T., and Komrokji, R.S. (2021). Triple-Negative Myelofibrosis: Disease Features, Response to Treatment and Outcomes. Blood 138, 1494–1494. 10.1182/blood-2021-151978 %J Blood.

17. Tefferi, A., Lasho, T.L., Finke, C.M., Knudson, R.A., Ketterling, R., Hanson, C.H., Maffioli, M., Caramazza, D., Passamonti, F., and Pardanani, A. (2014). CALR vs JAK2 vs MPL-mutated or triple-negative myelofibrosis: clinical, cytogenetic and molecular comparisons. Leukemia 28, 1472–1477. 10.1038/leu.2014.3.

18. Ross, D.M., Babon, J.J., Tvorogov, D., and Thomas, D. (2021). Persistence of myelofibrosis treated with ruxolitinib: biology and clinical implications. Haematologica 106, 1244–1253. 10.3324/haematol.2020.262691.

19. Vachhani, P., Verstovsek, S., and Bose, P. (2022). Disease Modification in Myelofibrosis: An Elusive Goal? J Clin Oncol 10;40(11):1147–1154 JCO.21.02246. 10.1200/jco.21.02246.

20. Dang, C.V. (1999). c-Myc target genes involved in cell growth, apoptosis, and metabolism. Mol Cell Biol 19, 1–11.

21. Dang, C.V. (2012). MYC on the path to cancer. Cell 149, 22–35. 10.1016/j.cell.2012.03.003.

22. Dang, C.V., O’Donnell, K.A., Zeller, K.I., Nguyen, T., Osthus, R.C., and Li, F. (2006). The c-Myc target gene network. Semin Cancer Biol 16, 253–264. 10.1016/j.semcancer.2006.07.014.

23. Adams, J.M., Harris, A.W., Pinkert, C.A., Corcoran, L.M., Alexander, W.S., Cory, S., Palmiter, R.D., and Brinster, R.L. (1985). The c-myc oncogene driven by immunoglobulin enhancers induces lymphoid malignancy in transgenic mice. Nature 318, 533–538. 10.1038/318533a0.

24. Taub, R., Kirsch, I., Morton, C., Lenoir, G., Swan, D., Tronick, S., Aaronson, S., and Leder, P. (1982). Translocation of the c-myc gene into the immunoglobulin heavy chain locus in human Burkitt lymphoma and murine plasmacytoma cells. Proc Natl Acad Sci 79, 7837–7841. doi:10.1073/pnas.79.24.7837.

25. Smith, D.P., Bath, M.L., Harris, A.W., and Cory, S. (2005). T-cell lymphomas mask slower developing B-lymphoid and myeloid tumours in transgenic mice with broad haemopoietic expression of MYC. Oncogene 24, 3544–3553. 10.1038/sj.onc.1208399.

26. Yun, S., Vincelette, N.D., Yu, X., Watson, G.W., Fernandez, M.R., Yang, C., Hitosugi, T., Cheng, C.-H., Freischel, A.R., Zhang, L., et al. (2021). TFEB Links MYC Signaling to Epigenetic Control of Myeloid Differentiation and Acute Myeloid Leukemia. Blood Cancer Discov 2, 162–185. 10.1158/2643-3230.BCD-20-0029 %J Blood Cancer Discovery.

27. Gajzer, D., Logothetis, C.N., Sallman, D.A., Calon, G., Babu, A., Chan, O., Vincelette, N.D., Volpe, V.O., Al Ali, N.H., Basra, P., et al. (2021). MYC overexpression is associated with an early disease progression from MDS to AML. Leuk Res 111, 106733. 10.1016/j.leukres.2021.106733.

28. Ohanian, M., Rozovski, U., Kanagal-Shamanna, R., Abruzzo, L.V., Loghavi, S., Kadia, T., Futreal, A., Bhalla, K., Zuo, Z., Huh, Y.O., et al. (2018). MYC protein expression is an important prognostic factor in acute myeloid leukemia. Leukemia & Lymphoma 60, 37–48. 10.1080/10428194.2018.1464158.

29. Sakamoto, K., Katayama, R., Asaka, R., Sakata, S., Baba, S., Nakasone, H., Koike, S., Tsuyama, N., Dobashi, A., Sasaki, M., et al. (2018). Recurrent 8q24 rearrangement in blastic plasmacytoid dendritic cell neoplasm: association with immunoblastoid cytomorphology, MYC expression, and drug response. Leukemia 32, 2590–2603. 10.1038/s41375-018-0154-5.

30. Sloand, E.M., Pfannes, L., Chen, G., Shah, S., Solomou, E.E., Barrett, J., and Young, N.S. (2006). CD34 cells from patients with trisomy 8 myelodysplastic syndrome (MDS) express early apoptotic markers but avoid programmed cell death by up-regulation of antiapoptotic proteins. Blood 109, 2399–2405. 10.1182/blood-2006-01-030643 %J Blood.

31. Reavie, L., Buckley, Shannon M., Loizou, E., Takeishi, S., Aranda-Orgilles, B., Ndiaye-Lobry, D., Abdel-Wahab, O., Ibrahim, S., Nakayama, Keiichi I., and Aifantis, I. (2013). Regulation of c-Myc Ubiquitination Controls Chronic Myelogenous Leukemia Initiation and Progression. Cancer Cell 23, 362–375. 10.1016/j.ccr.2013.01.025.

32. Luo, H., Li, Q., O’Neal, J., Kreisel, F., Le Beau, M.M., and Tomasson, M.H. (2005). c-Myc rapidly induces acute myeloid leukemia in mice without evidence of lymphoma-associated antiapoptotic mutations. Blood 106, 2452–2461. 10.1182/blood-2005-02-0734.

33. Bulaeva, E., Pellacani, D., Nakamichi, N., Hammond, C.A., Beer, P.A., Lorzadeh, A., Moksa, M., Carles, A., Bilenky, M., Lefort, S., et al. (2020). MYC-induced human acute myeloid leukemia requires a continuing IL-3/GM-CSF costimulus. Blood 136, 2764–2773. 10.1182/blood.2020006374 %J Blood.

34. Yun, S., Sharma, R., Chan, O., Vincelette, N.D., Sallman, D.A., Sweet, K., Padron, E., Komrokji, R., Lancet, J.E., Abraham, I., et al. (2019). Prognostic significance of MYC oncoprotein expression on survival outcome in patients with acute myeloid leukemia with myelodysplasia related changes (AML-MRC). Leuk Res 84, 106194. 10.1016/j.leukres.2019.106194.

35. Haase, D. (2008). Cytogenetic features in myelodysplastic syndromes. Annals of Hematology 87, 515–526. 10.1007/s00277-008-0483-y.

36. Wassie, E., Finke, C., Gangat, N., Lasho, T.L., Pardanani, A., Hanson, C.A., Ketterling, R.P., and Tefferi, A. (2015). A compendium of cytogenetic abnormalities in myelofibrosis: molecular and phenotypic correlates in 826 patients. Br J Haematol 169, 71–76. 10.1111/bjh.13260.

37. Vaidya, R., Caramazza, D., Begna, K.H., Gangat, N., Van Dyke, D.L., Hanson, C.A., Pardanani, A., and Tefferi, A. (2011). Monosomal karyotype in primary myelofibrosis is detrimental to both overall and leukemia-free survival. Blood 117, 5612–5615. 10.1182/blood-2010-11-320002 %J Blood.

38. Tefferi, A., Nicolosi, M., Mudireddy, M., Lasho, T.L., Gangat, N., Begna, K.H., Hanson, C.A., Ketterling, R.P., and Pardanani, A. (2018). Revised cytogenetic risk stratification in primary myelofibrosis: analysis based on 1002 informative patients. Leukemia 32, 1189–1199. 10.1038/s41375-018-0018-z.

39. Papaemmanuil, E., Gerstung, M., Bullinger, L., Gaidzik, V.I., Paschka, P., Roberts, N.D., Potter, N.E., Heuser, M., Thol, F., Bolli, N., et al. (2016). Genomic Classification and Prognosis in Acute Myeloid Leukemia. N Engl J Med 374, 2209–2221. doi:10.1056/NEJMoa1516192.

40. Kleppe, M., Koche, R., Zou, L., van Galen, P., Hill, C.E., Dong, L., De Groote, S., Papalexi, E., Hanasoge Somasundara, A.V., Cordner, K., et al. (2018). Dual Targeting of Oncogenic Activation and Inflammatory Signaling Increases Therapeutic Efficacy in Myeloproliferative Neoplasms. Cancer Cell 33, 29–43.e27. 10.1016/j.ccell.2017.11.009.

41. Huang, S.-M.A., Wang, A., Greco, R., Li, Z., Barberis, C., Tabart, M., Patel, V., Schio, L., Hurley, R., Cheng, H., et al. (2014). Combination of PIM and JAK2 inhibitors synergistically suppresses cell proliferation and overcomes drug resistance of myeloproliferative neoplasms. Oncotarget 30;5(10):3362–74.

42. Kim, S.-K., Knight, D.A., Jones, L.R., Vervoort, S., Ng, A.P., Seymour, J.F., Bradner, J.E., Waibel, M., Kats, L., and Johnstone, R.W. (2018). JAK2 is dispensable for maintenance of JAK2 mutant B-cell acute lymphoblastic leukemias. Genes Dev 32, 849–864. 10.1101/gad.307504.117.

43. Wang, S., Song, R., Wang, Z., Jing, Z., Wang, S., and Ma, J. (2018). S100A8/A9 in Inflammation. Front Immunol 9, 1298. 10.3389/fimmu.2018.01298.

44. Gleitz, H.F.E., Benabid, A., and Schneider, R.K. (2021). Still a burning question: the interplay between inflammation and fibrosis in myeloproliferative neoplasms. Curr Opin Hematol 28, 364–371. 10.1097/moh.0000000000000669.

45. Leimkühler, N.B., Gleitz, H.F.E., Ronghui, L., Snoeren, I.A.M., Fuchs, S.N.R., Nagai, J.S., Banjanin, B., Lam, K.H., Vogl, T., Kuppe, C., et al. (2021). Heterogeneous bone-marrow stromal progenitors drive myelofibrosis via a druggable alarmin axis. Cell Stem Cell 28, 637–652.e638. 10.1016/j.stem.2020.11.004.

46. Zambetti, Noemi A., Ping, Z., Chen, S., Kenswil, Keane J.G., Mylona, Maria A., Sanders, Mathijs A., Hoogenboezem, Remco M., Bindels, Eric M.J., Adisty, Maria N., Van Strien, Paulina M.H., et al. (2016). Mesenchymal Inflammation Drives Genotoxic Stress in Hematopoietic Stem Cells and Predicts Disease Evolution in Human Pre-leukemia. Cell Stem Cell 19, 613–627. 10.1016/j.stem.2016.08.021.

47. Bresnick, A.R., Weber, D.J., and Zimmer, D.B. (2015). S100 proteins in cancer. Nature Reviews Cancer 15, 96–109. 10.1038/nrc3893.

48. Vogl, T., Tenbrock, K., Ludwig, S., Leukert, N., Ehrhardt, C., van Zoelen, M.A.D., Nacken, W., Foell, D., van der Poll, T., Sorg, C., and Roth, J. (2007). Mrp8 and Mrp14 are endogenous activators of Toll-like receptor 4, promoting lethal, endotoxin-induced shock. Nature Medicine 13, 1042–1049. 10.1038/nm1638.

49. Yun, S., Geyer, S.M., Komrokji, R.S., Al Ali, N.H., Song, J., Hussaini, M., Sweet, K.L., Lancet, J.E., List, A.F., Padron, E., and Sallman, D.A. (2020). Prognostic significance of serial molecular annotation in myelodysplastic syndromes (MDS) and secondary acute myeloid leukemia (sAML). Leukemia. 35, 1145–1155 10.1038/s41375-020-0997-4.

50. Kruspig, B., Monteverde, T., Neidler, S., Hock, A., Kerr, E., Nixon, C., Clark, W., Hedley, A., Laing, S., Coffelt, S.B., et al. (2018). The ERBB network facilitates KRAS-driven lung tumorigenesis. Sci Transl Med 10, eaao2565. 10.1126/scitranslmed.aao2565 %J Science Translational Medicine.

51. Harris, A.W., Pinkert, C.A., Crawford, M., Langdon, W.Y., Brinster, R.L., and Adams, J.M. (1988). The E mu-myc transgenic mouse. A model for high-incidence spontaneous lymphoma and leukemia of early B cells. Journal of Experimental Medicine 167, 353–371. 10.1084/jem.167.2.353 %J Journal of Experimental Medicine.

52. Fernandez, M.R., Schaub, F.X., Yang, C., Li, W., Yun, S., Schaub, S.K., Dorsey, F.C., Liu, M., Steeves, M.A., Ballabio, A., et al. (2022). Disrupting the MYC-TFEB Circuit Impairs Amino Acid Homeostasis and Provokes Metabolic Anergy. Cancer Research 82, 1234–1250. 10.1158/0008-5472.CAN-21-1168 %J Cancer Research.

53. Smith, D.P., Bath, M.L., Metcalf, D., Harris, A.W., and Cory, S. (2006). MYC levels govern hematopoietic tumor type and latency in transgenic mice. Blood 108, 653–661. 10.1182/blood-2006-01-0172 %J Blood.

54. Ohshima, K., Kikuchi, M., and Takeshita, M. (1995). A megakaryocyte analysis of the bone marrow in patients with myelodysplastic syndrome, myeloproliferative disorder and allied disorders. J Pathol 177, 181–189. 10.1002/path.1711770212.

55. Rabellino, E.M., Levene, R.B., Nachman, R.L., and Leung, L.L.K. (1984). Human Megakaryocytes III. Characterization in Myeloproliferative Disorders. Blood 63, 615–622. 10.1182/blood.V63.3.615.615.

56. Arber, D.A., Orazi, A., Hasserjian, R., Thiele, J., Borowitz, M.J., Le Beau, M.M., Bloomfield, C.D., Cazzola, M., and Vardiman, J.W. (2016). The 2016 revision to the World Health Organization classification of myeloid neoplasms and acute leukemia. Blood 127, 2391–2405. 10.1182/blood-2016-03-643544.

57. Guglielmelli, P., Pacilli, A., Rotunno, G., Rumi, E., Rosti, V., Delaini, F., Maffioli, M., Fanelli, T., Pancrazzi, A., Pietra, D., et al. (2017). Presentation and outcome of patients with 2016 WHO diagnosis of prefibrotic and overt primary myelofibrosis. Blood 129, 3227–3236. 10.1182/blood-2017-01-761999 %J Blood.

58. Khoury, J.D., Solary, E., Abla, O., Akkari, Y., Alaggio, R., Apperley, J.F., Bejar, R., Berti, E., Busque, L., Chan, J.K.C., et al. (2022). The 5th edition of the World Health Organization Classification of Haematolymphoid Tumours: Myeloid and Histiocytic/Dendritic Neoplasms. Leukemia 36, 1703–1719. 10.1038/s41375-022-01613-1.

59. Baumeister, J., Maié, T., Chatain, N., Gan, L., Weinbergerova, B., de Toledo, M.A.S., Eschweiler, J., Maurer, A., Mayer, J., Kubesova, B., et al. (2021). Early and late stage MPN patients show distinct gene expression profiles in CD34+ cells. Annals of Hematology 100, 2943–2956. 10.1007/s00277-021-04615-8.

60. Čokić, V.P., Mossuz, P., Han, J., Socoro, N., Beleslin-Čokić, B.B., Mitrović, O., Subotički, T., Diklić, M., Leković, D., Gotić, M., et al. (2015). Microarray and Proteomic Analyses of Myeloproliferative Neoplasms with a Highlight on the mTOR Signaling Pathway. PLOS ONE 10, e0135463. 10.1371/journal.pone.0135463.

61. Bergsbaken, T., Fink, S.L., and Cookson, B.T. (2009). Pyroptosis: host cell death and inflammation. Nature Reviews Microbiology 7, 99–109. 10.1038/nrmicro2070.

62. von Bauer, R., Oikonomou, D., Sulaj, A., Mohammed, S., Hotz-Wagenblatt, A., Gröne, H.J., Arnold, B., Falk, C., Luethje, D., Erhardt, A., et al. (2013). CD166/ALCAM mediates proinflammatory effects of S100B in delayed type hypersensitivity. J Immunol 191, 369–377. 10.4049/jimmunol.1201864.

63. Wakahashi, K., Minagawa, K., Kawano, Y., Kawano, H., Suzuki, T., Ishii, S., Sada, A., Asada, N., Sato, M., Kato, S., et al. (2019). Vitamin D receptor–mediated skewed differentiation of macrophages initiates myelofibrosis and subsequent osteosclerosis. Blood 133, 1619–1629. 10.1182/blood-2018-09-876615 %J Blood.

64. Sunahori, K., Yamamura, M., Yamana, J., Takasugi, K., Kawashima, M., Yamamoto, H., Chazin, W.J., Nakatani, Y., Yui, S., and Makino, H. (2006). The S100A8/A9 heterodimer amplifies proinflammatory cytokine production by macrophages via activation of nuclear factor kappa B and p38 mitogen-activated protein kinase in rheumatoid arthritis. Arthritis Research & Therapy 8, R69. 10.1186/ar1939.

65. Okada, K., Arai, S., Itoh, H., Adachi, S., Hayashida, M., Nakase, H., and Ikemoto, M. (2016). CD68 on rat macrophages binds tightly to S100A8 and S100A9 and helps to regulate the cells’ immune functions. Journal of leukocyte biology 100, 1093–1104. 10.1189/jlb.2A0415-170RRR.

66. Cheng, P., Corzo, C.A., Luetteke, N., Yu, B., Nagaraj, S., Bui, M.M., Ortiz, M., Nacken, W., Sorg, C., Vogl, T., et al. (2008). Inhibition of dendritic cell differentiation and accumulation of myeloid-derived suppressor cells in cancer is regulated by S100A9 protein. Journal of Experimental Medicine 205, 2235–2249. 10.1084/jem.20080132 %J Journal of Experimental Medicine.

67. Björk, P., Björk, A., Vogl, T., Stenström, M., Liberg, D., Olsson, A., Roth, J., Ivars, F., and Leanderson, T. (2009). Identification of Human S100A9 as a Novel Target for Treatment of Autoimmune Disease via Binding to Quinoline-3-Carboxamides. PLOS Biology 7, e1000097. 10.1371/journal.pbio.1000097.

68. Sternberg, C., Armstrong, A., Pili, R., Ng, S., Huddart, R., Agarwal, N., Khvorostenko, D., Lyulko, O., Brize, A., Vogelzang, N., et al. (2016). Randomized, Double-Blind, Placebo-Controlled Phase III Study of Tasquinimod in Men With Metastatic Castration-Resistant Prostate Cancer. J Clin Oncol 34, 2636–2643. 10.1200/jco.2016.66.9697.

69. Han, H., Jain, A.D., Truica, M.I., Izquierdo-Ferrer, J., Anker, J.F., Lysy, B., Sagar, V., Luan, Y., Chalmers, Z.R., Unno, K., et al. (2019). Small-Molecule MYC Inhibitors Suppress Tumor Growth and Enhance Immunotherapy. Cancer Cell 36, 483–497.e415. 10.1016/j.ccell.2019.10.001.

70. Bolouri, H., Farrar, J.E., Triche Jr, T., Ries, R.E., Lim, E.L., Alonzo, T.A., Ma, Y., Moore, R., Mungall, A.J., Marra, M.A., et al. (2017). The molecular landscape of pediatric acute myeloid leukemia reveals recurrent structural alterations and age-specific mutational interactions. Nature Medicine 24, 103. 10.1038/nm.4439 https://www.nature.com/articles/nm.4439#supplementary-information.

71. Tyner, J.W., Tognon, C.E., Bottomly, D., Wilmot, B., Kurtz, S.E., Savage, S.L., Long, N., Schultz, A.R., Traer, E., Abel, M., et al. (2018). Functional genomic landscape of acute myeloid leukaemia. Nature 562, 526–531. 10.1038/s41586-018-0623-z.

72. Bejar, R., Stevenson, K., Abdel-Wahab, O., Galili, N., Nilsson, B., Garcia-Manero, G., Kantarjian, H., Raza, A., Levine, R.L., Neuberg, D., and Ebert, B.L. (2011). Clinical Effect of Point Mutations in Myelodysplastic Syndromes. New England Journal of Medicine 364, 2496–2506. 10.1056/NEJMoa1013343.

73. Haferlach, T., Nagata, Y., Grossmann, V., Okuno, Y., Bacher, U., Nagae, G., Schnittger, S., Sanada, M., Kon, A., Alpermann, T., et al. (2014). Landscape of genetic lesions in 944 patients with myelodysplastic syndromes. Leukemia 28, 241–247. 10.1038/leu.2013.336.

74. Yoshizato, T., Nannya, Y., Atsuta, Y., Shiozawa, Y., Iijima-Yamashita, Y., Yoshida, K., Shiraishi, Y., Suzuki, H., Nagata, Y., Sato, Y., et al. (2017). Genetic abnormalities in myelodysplasia and secondary acute myeloid leukemia: impact on outcome of stem cell transplantation. Blood 129, 2347–2358. 10.1182/blood-2016-12-754796.

75. Nagata, Y., Makishima, H., Kerr, C.M., Przychodzen, B.P., Aly, M., Goyal, A., Awada, H., Asad, M.F., Kuzmanovic, T., Suzuki, H., et al. (2019). Invariant patterns of clonal succession determine specific clinical features of myelodysplastic syndromes. Nature Communications 10, 5386. 10.1038/s41467-019-13001-y.

76. Papaemmanuil, E., Gerstung, M., Malcovati, L., Tauro, S., Gundem, G., Van Loo, P., Yoon, C.J., Ellis, P., Wedge, D.C., Pellagatti, A., et al. (2013). Clinical and biological implications of driver mutations in myelodysplastic syndromes. Blood 122, 3616–3627. 10.1182/blood-2013-08-518886.

77. Al-Ghamdi, Y.A., Lake, J., Bagg, A., Thakral, B., Wang, S.A., Bueso-Ramos, C., Masarova, L., Verstovsek, S., Rogers, H.J., Hsi, E.D., et al. (2023). Triple-Negative Primary Myelofibrosis: A Bone Marrow Pathology Group Study. Mod Pathol 36. 10.1016/j.modpat.2022.100016.

78. Hérault, L., Poplineau, M., Mazuel, A., Platet, N., Remy, É., and Duprez, E. (2021). Single-cell RNA-seq reveals a concomitant delay in differentiation and cell cycle of aged hematopoietic stem cells. BMC Biology 19, 19. 10.1186/s12915-021-00955-z.

79. Wilson, A., Murphy, M.J., Oskarsson, T., Kaloulis, K., Bettess, M.D., Oser, G.M., Pasche, A.-C., Knabenhans, C., MacDonald, H.R., and Trumpp, A. (2004). c-Myc controls the balance between hematopoietic stem cell self-renewal and differentiation. Genes Dev 18, 2747–2763. 10.1101/gad.313104.

80. Guo, Y., Niu, C., Breslin, P., Tang, M., Zhang, S., Wei, W., Kini, A.R., Paner, G.P., Alkan, S., Morris, S.W., et al. (2009). c-Myc–mediated control of cell fate in megakaryocyte-erythrocyte progenitors. Blood 114, 2097–2106. 10.1182/blood-2009-01-197947 %J Blood.

81. Felsher, D.W., and Bishop, J.M. (1999). Reversible Tumorigenesis by MYC in Hematopoietic Lineages. Molecular Cell 4, 199–207. 10.1016/S1097-2765(00)80367-6.

82. Bhatia, M., Bonnet, D., Kapp, U., Wang, J.C.Y., Murdoch, B., and Dick, J.E. (1997). Quantitative Analysis Reveals Expansion of Human Hematopoietic Repopulating Cells After Short-term Ex Vivo Culture. Journal of Experimental Medicine 186, 619–624. 10.1084/jem.186.4.619 %J Journal of Experimental Medicine.

83. Conneally, E., Cashman, J., Petzer, A., and Eaves, C. (1997). Expansion in vitro of transplantable human cord blood stem cells demonstrated using a quantitative assay of their lympho-myeloid repopulating activity in nonobese diabetic scid/scid mice. Proc Natl Acad Sci 94, 9836–9841. doi:10.1073/pnas.94.18.9836.

84. Gammaitoni, L., Bruno, S., Sanavio, F., Gunetti, M., Kollet, O., Cavalloni, G., Falda, M., Fagioli, F., Lapidot, T., Aglietta, M., and Piacibello, W. (2003). Ex vivo expansion of human adult stem cells capable of primary and secondary hemopoietic reconstitution. Experimental Hematology 31, 261–270. 10.1016/S0301-472X(02)01077-9.

85. Ueda, T., Tsuji, K., Yoshino, H., Ebihara, Y., Yagasaki, H., Hisakawa, H., Mitsui, T., Manabe, A., Tanaka, R., Kobayashi, K., et al. (2000). Expansion of human NOD/SCID-repopulating cells by stem cell factor, Flk2/Flt3 ligand, thrombopoietin, IL-6, and soluble IL-6 receptor. The Journal of Clinical Investigation 105, 1013–1021. 10.1172/JCI8583.

86. Marx-Blümel, L., Marx, C., Sonnemann, J., Weise, F., Hampl, J., Frey, J., Rothenburger, L., Cirri, E., Rahnis, N., Koch, P., et al. (2021). Molecular characterization of hematopoietic stem cells after in vitro amplification on biomimetic 3D PDMS cell culture scaffolds. Scientific Reports 11, 21163. 10.1038/s41598-021-00619-6.

87. Italiani, P., and Boraschi, D. (2014). From Monocytes to M1/M2 Macrophages: Phenotypical vs. Functional Differentiation. Front Immunol 5. 514 10.3389/fimmu.2014.00514.

88. Orecchioni, M., Ghosheh, Y., Pramod, A.B., and Ley, K. (2019). Macrophage Polarization: Different Gene Signatures in M1(LPS+) vs. Classically and M2(LPS–) vs. Alternatively Activated Macrophages. Front Immunol 10. 1084 10.3389/fimmu.2019.01084.

89. Molitor, D.C.A., Boor, P., Buness, A., Schneider, R.K., Teichmann, L.L., Körber, R.-M., Horvath, G.L., Koschmieder, S., and Gütgemann, I. (2021). Macrophage frequency in the bone marrow correlates with morphologic subtype of myeloproliferative neoplasm. Annals of Hematology 100, 97–104. 10.1007/s00277-020-04304-y.

90. Funakoshi-Tago, M., Sumi, K., Kasahara, T., and Tago, K. (2013). Critical Roles of Myc-ODC Axis in the Cellular Transformation Induced by Myeloproliferative Neoplasm-Associated JAK2 V617F Mutant. PLOS ONE 8, e52844. 10.1371/journal.pone.0052844.

91. Wernig, G., Gonneville, J.R., Crowley, B.J., Rodrigues, M.S., Reddy, M.M., Hudon, H.E., Walz, C., Reiter, A., Podar, K., Royer, Y., et al. (2008). The Jak2V617F oncogene associated with myeloproliferative diseases requires a functional FERM domain for transformation and for expression of the Myc and Pim proto-oncogenes. Blood 111, 3751–3759. 10.1182/blood-2007-07-102186 %J Blood.

92. Decker, M., Martinez-Morentin, L., Wang, G., Lee, Y., Liu, Q., Leslie, J., and Ding, L. (2017). Leptin-receptor-expressing bone marrow stromal cells are myofibroblasts in primary myelofibrosis. Nature Cell Biology 19, 677–688. 10.1038/ncb3530.

93. Schneider, R.K., Mullally, A., Dugourd, A., Peisker, F., Hoogenboezem, R., Van Strien, P.M.H., Bindels, E.M., Heckl, D., Büsche, G., Fleck, D., et al. (2017). Gli1+ Mesenchymal Stromal Cells Are a Key Driver of Bone Marrow Fibrosis and an Important Cellular Therapeutic Target. Cell Stem Cell 20, 785–800.e788. 10.1016/j.stem.2017.03.008.

94. Kramann, R., and Schneider, R.K. (2018). The identification of fibrosis-driving myofibroblast precursors reveals new therapeutic avenues in myelofibrosis. Blood 131, 2111–2119. 10.1182/blood-2018-02-834820 %J Blood.

95. Mullally, A., Lane, S.W., Ball, B., Megerdichian, C., Okabe, R., Al-Shahrour, F., Paktinat, M., Haydu, J.E., Housman, E., Lord, A.M., et al. (2010). Physiological Jak2V617F Expression Causes a Lethal Myeloproliferative Neoplasm with Differential Effects on Hematopoietic Stem and Progenitor Cells. Cancer Cell 17, 584–596. 10.1016/j.ccr.2010.05.015.

96. Akada, H., Yan, D., Zou, H., Fiering, S., Hutchison, R.E., and Mohi, M.G. (2010). Conditional expression of heterozygous or homozygous Jak2V617F from its endogenous promoter induces a polycythemia vera–like disease. Blood 115, 3589–3597. 10.1182/blood-2009-04-215848 %J Blood.

97. Li, J., Spensberger, D., Ahn, J.S., Anand, S., Beer, P.A., Ghevaert, C., Chen, E., Forrai, A., Scott, L.M., Ferreira, R., et al. (2010). JAK2 V617F impairs hematopoietic stem cell function in a conditional knock-in mouse model of JAK2 V617F–positive essential thrombocythemia. Blood 116, 1528–1538. 10.1182/blood-2009-12-259747 %J Blood.

98. Marty, C., Lacout, C., Martin, A., Hasan, S., Jacquot, S., Birling, M.-C., Vainchenker, W., and Villeval, J.-L. (2010). Myeloproliferative neoplasm induced by constitutive expression of JAK2V617F in knock-in mice. Blood 116, 783–787. 10.1182/blood-2009-12-257063 %J Blood.

99. Chapeau, E.A., Mandon, E., Gill, J., Romanet, V., Ebel, N., Powajbo, V., Andraos-Rey, R., Qian, Z., Kininis, M., Zumstein-Mecker, S., et al. (2019). A conditional inducible JAK2V617F transgenic mouse model reveals myeloproliferative disease that is reversible upon switching off transgene expression. PLOS ONE 14, e0221635. 10.1371/journal.pone.0221635.

100. Li, J., Kent, D.G., Chen, E., and Green, A.R. (2011). Mouse models of myeloproliferative neoplasms: JAK of all grades. Disease Models & Mechanisms 4, 311–317. 10.1242/dmm.006817 %J Disease Models & Mechanisms.

101. Jacquelin, S., Kramer, F., Mullally, A., and Lane, S.W. (2020). Murine Models of Myelofibrosis. Cancer (Basel)12, 2381.

102. Niroula, A., Sekar, A., Murakami, M.A., Trinder, M., Agrawal, M., Wong, W.J., Bick, A.G., Uddin, M.M., Gibson, C.J., Griffin, G.K., et al. (2021). Distinction of lymphoid and myeloid clonal hematopoiesis. Nature Medicine 27, 1921–1927. 10.1038/s41591-021-01521-4.

103. Saiki, R., Momozawa, Y., Nannya, Y., Nakagawa, M.M., Ochi, Y., Yoshizato, T., Terao, C., Kuroda, Y., Shiraishi, Y., Chiba, K., et al. (2021). Combined landscape of single-nucleotide variants and copy number alterations in clonal hematopoiesis. Nature Medicine 27, 1239–1249. 10.1038/s41591-021-01411-9.

104. Loh, P.-R., Genovese, G., and McCarroll, S.A. (2020). Monogenic and polygenic inheritance become instruments for clonal selection. Nature 584, 136–141. 10.1038/s41586-020-2430-6.

105. Loh, P.-R., Genovese, G., Handsaker, R.E., Finucane, H.K., Reshef, Y.A., Palamara, P.F., Birmann, B.M., Talkowski, M.E., Bakhoum, S.F., McCarroll, S.A., and Price, A.L. (2018). Insights into clonal haematopoiesis from 8,342 mosaic chromosomal alterations. Nature 559, 350–355. 10.1038/s41586-018-0321-x.

106. Hormaechea-Agulla, D., Matatall, K.A., Le, D.T., Kain, B., Long, X., Kus, P., Jaksik, R., Challen, G.A., Kimmel, M., and King, K.Y. (2021). Chronic infection drives Dnmt3a-loss-of-function clonal hematopoiesis via IFN&#x3b3; signaling. Cell Stem Cell 28, 1428–1442.e1426. 10.1016/j.stem.2021.03.002.

107. Liao, M., Chen, R., Yang, Y., He, H., Xu, L., Jiang, Y., Guo, Z., He, W., Jiang, H., and Wang, J. (2022). Aging-elevated inflammation promotes DNMT3A R878H-driven clonal hematopoiesis. Acta Pharmaceutica Sinica B 12, 678–691. 10.1016/j.apsb.2021.09.015.

108. Fuster, J.J., MacLauchlan, S., Zuriaga, M.A., Polackal, M.N., Ostriker, A.C., Chakraborty, R., Wu, C.-L., Sano, S., Muralidharan, S., Rius, C., et al. (2017). Clonal hematopoiesis associated with TET2 deficiency accelerates atherosclerosis development in mice. Science 355, 842–847. doi:10.1126/science.aag1381.

109. Fidler, T.P., Xue, C., Yalcinkaya, M., Hardaway, B., Abramowicz, S., Xiao, T., Liu, W., Thomas, D.G., Hajebrahimi, M.A., Pircher, J., et al. (2021). The AIM2 inflammasome exacerbates atherosclerosis in clonal haematopoiesis. Nature 592, 296–301. 10.1038/s41586-021-03341-5.

110. Sallman, D.A., Komrokji, R., Vaupel, C., Cluzeau, T., Geyer, S.M., McGraw, K.L., Al Ali, N.H., Lancet, J., McGinniss, M.J., Nahas, S., et al. (2016). Impact of TP53 mutation variant allele frequency on phenotype and outcomes in myelodysplastic syndromes. Leukemia 30, 666–673. 10.1038/leu.2015.304.

111. Yun, S., Komrokji, R., Al Ali, N., Song, J., Vaupel, C., Hussaini, M., Sweet, K., Lancet, J., Hall, J., List, A., et al. (2017). Prognostic Significance of Serial Molecular Annotation in Myelodysplastic Syndromes (MDS) and Secondary Acute Myeloid Leukemia (sAML). Clinical Lymphoma, Myeloma and Leukemia 17, S344–S345. 10.1016/j.clml.2017.07.166.

112. Feldman, A.T., and Wolfe, D. (2014). Tissue Processing and Hematoxylin and Eosin Staining. In Histopathology: Methods and Protocols, C.E. Day, ed. (Springer New York), pp. 31–43. 10.1007/978-1-4939-1050-2_3.

113. Yoshimi, A., Balasis, M.E., Vedder, A., Feldman, K., Ma, Y., Zhang, H., Lee, S.C.-W., Letson, C., Niyongere, S., Lu, S.X., et al. (2017). Robust patient-derived xenografts of MDS/MPN overlap syndromes capture the unique characteristics of CMML and JMML. Blood 130, 397–407. 10.1182/blood-2017-01-763219.

114. Yun, S., Vincelette, N.D., Knorr, K.L., Almada, L.L., Schneider, P.A., Peterson, K.L., Flatten, K.S., Dai, H., Pratz, K.W., Hess, A.D., et al. (2016). 4EBP1/c-MYC/PUMA and NFκB/EGR1/BIM pathways underlie cytotoxicity of mTOR dual inhibitors in malignant lymphoid cells. Blood. 10.1182/blood-2015-02-629485.

115. Satija, R., Farrell, J.A., Gennert, D., Schier, A.F., and Regev, A. (2015). Spatial reconstruction of single-cell gene expression data. Nat Biotechnol 33, 495–502. 10.1038/nbt.3192.

116. Wolock, S.L., Lopez, R., and Klein, A.M. (2019). Scrublet: Computational Identification of Cell Doublets in Single-Cell Transcriptomic Data. Cell Syst 8, 281–291 e289. 10.1016/j.cels.2018.11.005.

117. McGinnis, C.S., Murrow, L.M., and Gartner, Z.J. (2019). DoubletFinder: Doublet Detection in Single-Cell RNA Sequencing Data Using Artificial Nearest Neighbors. Cell Syst 8, 329–337 e324. 10.1016/j.cels.2019.03.003.

118. Germain, P., Lun, A., Macnair, W., and Robinson, M. (2021). Doublet identification in single-cell sequencing data using scDblFinder [version 1; peer review: 1 approved, 1 approved with reservations]. F1000Research 10. 10.12688/f1000research.73600.1.

119. Lun, A.T., McCarthy, D.J., and Marioni, J.C. (2016). A step-by-step workflow for low-level analysis of single-cell RNA-seq data with Bioconductor. F1000Res 5, 2122. 10.12688/f1000research.9501.2.

120. Hao, Y., Hao, S., Andersen-Nissen, E., Mauck, W.M., 3rd, Zheng, S., Butler, A., Lee, M.J., Wilk, A.J., Darby, C., Zager, M., et al. (2021). Integrated analysis of multimodal single-cell data. Cell 184, 3573–3587 e3529. 10.1016/j.cell.2021.04.048.

121. Aibar, S., González-Blas, C.B., Moerman, T., Huynh-Thu, V.A., Imrichova, H., Hulselmans, G., Rambow, F., Marine, J.-C., Geurts, P., Aerts, J., et al. (2017). SCENIC: single-cell regulatory network inference and clustering. Nature Methods 14, 1083–1086. 10.1038/nmeth.4463.

122. Stumpf, P.S., Du, X., Imanishi, H., Kunisaki, Y., Semba, Y., Noble, T., Smith, R.C.G., Rose-Zerili, M., West, J.J., Oreffo, R.O.C., et al. (2020). Transfer learning efficiently maps bone marrow cell types from mouse to human using single-cell RNA sequencing. Communications Biology 3, 736. 10.1038/s42003-020-01463-6.

123. van Galen, P., Hovestadt, V., Wadsworth Ii, M.H., Hughes, T.K., Griffin, G.K., Battaglia, S., Verga, J.A., Stephansky, J., Pastika, T.J., Lombardi Story, J., et al. (2019). Single-Cell RNA-Seq Reveals AML Hierarchies Relevant to Disease Progression and Immunity. Cell 176, 1265–1281.e1224. 10.1016/j.cell.2019.01.031.

124. Buenrostro, J.D., Corces, M.R., Lareau, C.A., Wu, B., Schep, A.N., Aryee, M.J., Majeti, R., Chang, H.Y., and Greenleaf, W.J. (2018). Integrated Single-Cell Analysis Maps the Continuous Regulatory Landscape of Human Hematopoietic Differentiation. Cell 173, 1535–1548.e1516. 10.1016/j.cell.2018.03.074.

125. Pishesha, N., Thiru, P., Shi, J., Eng, J.C., Sankaran, V.G., and Lodish, H.F. (2014). Transcriptional divergence and conservation of human and mouse erythropoiesis. Proceedings of the National Academy of Sciences 111, 4103–4108. doi:10.1073/pnas.1401598111.

126. Hay, S.B., Ferchen, K., Chetal, K., Grimes, H.L., and Salomonis, N. (2018). The Human Cell Atlas bone marrow single-cell interactive web portal. Experimental Hematology 68, 51–61. 10.1016/j.exphem.2018.09.004.

127. Wu, T., Hu, E., Xu, S., Chen, M., Guo, P., Dai, Z., Feng, T., Zhou, L., Tang, W., Zhan, L., et al. (2021). clusterProfiler 4.0: A universal enrichment tool for interpreting omics data. The Innovation 2. 10.1016/j.xinn.2021.100141.

128. Supek, F., Bošnjak, M., Škunca, N., and Šmuc, T. (2011). REVIGO Summarizes and Visualizes Long Lists of Gene Ontology Terms. PLOS ONE 6, e21800. 10.1371/journal.pone.0021800.

129. Jin, S., Guerrero-Juarez, C.F., Zhang, L., Chang, I., Ramos, R., Kuan, C.-H., Myung, P., Plikus, M.V., and Nie, Q. (2021). Inference and analysis of cell-cell communication using CellChat. Nature Communications 12, 1088. 10.1038/s41467-021-21246-9.

130. Türei, D., Korcsmáros, T., and Saez-Rodriguez, J. (2016). OmniPath: guidelines and gateway for literature-curated signaling pathway resources. Nature Methods 13, 966–967. 10.1038/nmeth.4077.

131. Dimitrov, D., Türei, D., Garrido-Rodriguez, M., Burmedi, P.L., Nagai, J.S., Boys, C., Ramirez Flores, R.O., Kim, H., Szalai, B., Costa, I.G., et al. (2022). Comparison of methods and resources for cell-cell communication inference from single-cell RNA-Seq data. Nature Communications 13, 3224. 10.1038/s41467-022-30755-0.

132. Türei, D., Valdeolivas, A., Gul, L., Palacio-Escat, N., Klein, M., Ivanova, O., Ölbei, M., Gábor, A., Theis, F., Módos, D., et al. (2021). Integrated intra- and intercellular signaling knowledge for multicellular omics analysis. Mol Syst Biol 17, e9923. 10.15252/msb.20209923.

133. Holland, C.H., Tanevski, J., Perales-Patón, J., Gleixner, J., Kumar, M.P., Mereu, E., Joughin, B.A., Stegle, O., Lauffenburger, D.A., Heyn, H., et al. (2020). Robustness and applicability of transcription factor and pathway analysis tools on single-cell RNA-seq data. Genome Biology 21, 36. 10.1186/s13059-020-1949-z.

134. Sabò, A., Kress, T.R., Pelizzola, M., de Pretis, S., Gorski, M.M., Tesi, A., Morelli, M.J., Bora, P., Doni, M., Verrecchia, A., et al. (2014). Selective transcriptional regulation by Myc in cellular growth control and lymphomagenesis. Nature 511, 488. 10.1038/nature13537https://www.nature.com/articles/nature13537#supplementary-information.

135. Rouillard, A.D., Gundersen, G.W., Fernandez, N.F., Wang, Z., Monteiro, C.D., McDermott, M.G., and Ma’ayan, A. (2016). The harmonizome: a collection of processed datasets gathered to serve and mine knowledge about genes and proteins. Database 2016. 10.1093/database/baw100.

136. Chan, J.K., Roth, J., Oppenheim, J.J., Tracey, K.J., Vogl, T., Feldmann, M., Horwood, N., and Nanchahal, J. (2012). Alarmins: awaiting a clinical response. The Journal of Clinical Investigation 122, 2711–2719. 10.1172/JCI62423.

137. Sergushichev, A.A. (2016). An algorithm for fast preranked gene set enrichment analysis using cumulative statistic calculation. bioRxiv, 060012. 10.1101/060012.

138. Gu, Z., Eils, R., and Schlesner, M. (2016). Complex heatmaps reveal patterns and correlations in multidimensional genomic data. Bioinformatics 32, 2847–2849. 10.1093/bioinformatics/btw313.

